# A molecular taxonomy of tumors independent of tissue-of-origin

**DOI:** 10.1101/2020.08.26.268987

**Authors:** Peter T. Nguyen, Simon G. Coetzee, Daniel L. Lakeland, Dennis J. Hazelett

**Affiliations:** The Center for Bioinformatics and Functional Genomics, Cedars-Sinai Medical Center, Los Angeles, California; Lakeland Applied Sciences LLC, Los Angeles, CA; Samuel Oschin Comprehensive Cancer Institute, Cedars-Sinai Medical Center, Los Angeles, CA

## Abstract

Cancer is a complex disease involving disrupted cellular metabolism, basic biochemical processes, and the microenvironment. However, despite some generally agreed upon unifying principles (Hanahan and Weinberg 2000, 2011), molecular signatures remain largely indistinguishable from tissue-of-origin, presenting a major barrier for precision health and individualized medicine. To address this challenge, we reduce mutation data to disruptions in a select set of pathways relevant to basic cell biology, from DNA replication to cellular communication. Using dimensionality reduction techniques, we assign tumor samples into ten clusters distinct from tissue-of-origin and largely free of bias from mutational burden or clinical stage. We show that the clusters vary in prognosis by modeling relative risk of death by cancer type and cluster. We identify cluster-specific mutations in different tissues, demonstrating that tissue-specific signatures contribute to common cellular phenotypes. Moreover, germline risk genes involved in replication fidelity and genome stability are equally distributed among clusters, contrary to the expectation that such genes are avatars of molecular subtype. We investigate metastatic and non-metastatic pathways, and show that most differences are cluster-specific. Some metastatic pathways from one cluster are cluster-specific pathways from non-metastatic tumors of another cluster, suggesting phenotypic convergence. Taken as a whole, our observations suggest that common driver genes combine with tissue-specific disruptions in tumor-promoting pathways to produce a limited number of distinct molecular phenotypes. Thus, we present a coherent view of global tumor biology, and explain how common cellular dysfunction might arise from tissue-specific mutations.

## Introduction

Recent advances in technology have greatly expanded our view into the mechanisms of cancer at the systems biology level. Array and next-generation sequencing technologies have made it possible to assay germline and somatic mutations, expression profiles, DNA methylation, and copy number variations from the same tissue samples. To take advantage of these tools, several large consortia, including the cancer genome atlas (TCGA) and pan-cancer analysis of whole genomes (PCAWG) sequenced large numbers of tumors and collected data from multiple assays that could be analyzed together, with the goal of integration and increased understanding of the mechanisms of cancer.

Considerable progress has been made analyzing these data. Statistical analyses identify hundreds of global and tissue-specific cancer driver genes (Dees et al. 2012; Tamborero, Gonzalez-Perez, and Lopez-Bigas 2013; Lawrence et al. 2014; Kumar et al. 2015; Tokheim et al. 2016; Jiang et al. 2019; Zhao et al. 2019) using approaches aimed at detecting when genes are mutated at a greater rate than expected due to chance. It has been estimated that fewer than five mutations in key oncogenes and/or tumor suppressors would be sufficient to transform a normal cell to a cancerous state (Vogelstein and Kinzler 2015; Iranzo, Martincorena, and Koonin 2018). Somewhat paradoxically, mutations in cancer driver genes are commonplace in healthy tissue and correlate with age and environmental exposures (Martincorena 2019).

Other studies provide a comprehensive view of mutations, gene expression and genomic signatures, with the express goal of understanding the common themes of all cancer independent of tissue of origin. Understanding cancer as a disease of the cell has long been a goal of the field as characterized in essays by Hanahan and Weinberg (2000); Hanahan and Weinberg (2011). The first of these studies in the genomic era identified 11 subtypes from 12 cancer types, using integrative analysis with co-equal weighting of gene expression, methylation, copy number and proteomics data (Hoadley et al. 2014). The principal finding was that tissue-of-origin is the predominant driving factor, though a significant proportion of tumors (~10%) could be reclassified independent of tissue-of-origin. In a second study involving 33 cancer types and a much greater number of tumors, the authors identify 28 clusters that could be further subdivided into organ specific groups, including pan-gastrointestinal, pan-gynecological, pan-squamous, pan-gynecological/squamous and pan-kidney (Hoadley et al. 2018).

More recently, attempts have been made to consider pan-cancer mutations and other data as disruptions of normal pathway activity. Sanchez-Vega et al. (2018) carried out near-exhaustive analysis of pathway enrichment using whole genome data. Network approaches address the sparseness of mutations, allowing genes to be influenced by mutations in the nearest network neighbors (Horn et al. 2018; Iranzo, Martincorena, and Koonin 2018). These methods are powerful especially when applied to a small number (on the order of hundreds) of tumors. One of the challenges in the field remains that beyond a couple hundred common driver genes and mutations, there may be thousands of moderate effect genes that occur at such low frequency–given tissue diversity–and low sample numbers that they are impossible to detect using positive selection theory. Some groups have attempted to address this challenge by applying machine learning to cancer data sets to discover groups of functionally related genes as they interact with larger pathways and networks (Kim and Kim 2018; Mourikis et al. 2019; Colaprico et al. 2020). These studies have proven very effective at highlighting fundamental disease phenotypes at the pathway level across cancers with different origins at the cellular and tissue level.

In this study, we attempt to understand how tissue-specific gene disruptions create common cancer phenotypes by focusing on discrete molecular pathways as the unit of disruption. Our approach strips all cell-type-specific information from the mutation data and equates gene-level mutations to cell-biological pathway disruptions. We use this heuristic to evaluate all cancers and show that, surprisingly, tumors that exhibit tissue-specific gene mutation patterns nonetheless fall into common categories of pathway disruption having unique prognoses in each cancer type.

## Results

### A taxonomy of tumors based on disrupted molecular pathways

To study cancer pathways we obtained a set of 7,607 solid tumor samples from The Cancer Genome Atlas (TCGA) through the Genomic Data Commons (GDC) portal. TCGA data were most appropriate for our study given the relative completeness of the patient metadata, particularly for survival and staging. We chose to analyze somatic mutations in exome sequencing data because the affected target gene is known unambiguously. Therefore, we selected all missense, nonsense, frameshift, stop-loss, untranslated region, and splicing mutations. In order to minimize bias from well-studied diseases and processes, we selected 377 Reactome pathways (see supplemental table ST1) of interest corresponding to basic cellular processes and biochemical pathways, excluding gene sets that correspond to miscellaneous categories (*e.g*. “transcription factors”) or disease associations (*e.g*. “mutated in colon cancer”) and filtered our gene list on membership in these pathways (total of 8,940 genes; see methods for details).

To avoid bias toward larger pathways (*i.e*. pathways with more member genes), we counted pathways as disrupted if one or more member genes were mutated. We do not attempt to calculate enrichment for mutations within a pathway. Data binarized by pathway are likely to be noisy for several reasons. First, point mutations can be deleterious (attenuating, hypomorphic or antimorphic) or activating (neomorphic or hypermorphic) in genes, and these can in turn act functionally as either oncogenes or tumor suppressors. For this study we assume a significant fraction of these mutations are generically disruptive to normal pathway activity since it is impossible to know the tumor promoting effects of all mutations, including rarely studied genes. Second, we know that low-expressed genes and non-expressed genes accumulate mutations at a higher rate due to transcription coupled repair (Kandoth et al. 2013; Kim and Kim 2018; Aitken et al. 2020; Cummings et al. 2020). To address this issue, we identified genes with low expression in each type of cancer and eliminated them for that cancer type only (see methods for details). Highly expressed genes could also have high mutation rates owing to transcription induced mutagenesis (Park, Qian, and Zhang 2012). We felt that such mechanisms result in cell-type-specific biases that might be biologically meaningful for predisposition to different classes of cancer however, and therefore chose not to exclude these genes from our analysis. After selecting our pathways and genes, we then compiled a matrix of the pathways, assigning a Boolean value of 1 to each pathway with one or more genes mutated and 0 for all others (**figure 1**).

**Figure 1:**
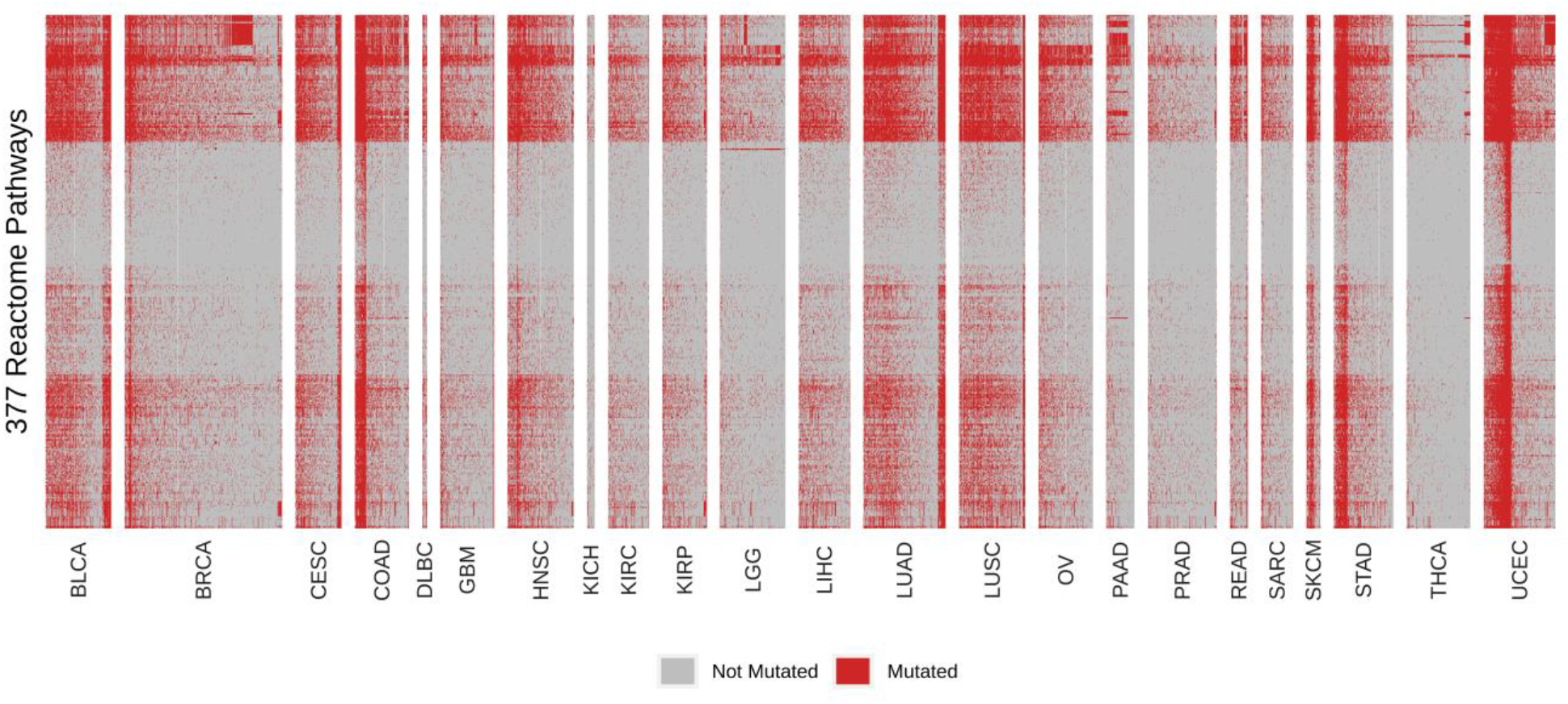
Molecular pathway profiles of tumor samples with one or more mutations. Each of 377 selected Reactome pathways (rows) is classified as disrupted if one or more genes is mutated in the tumor sample (columns) where red and grey represents the pathway is mutated and not, respectively. Tumors are grouped by tissue of origin using standardized abbreviations from the TCGA project.

We investigated this dataset further using multiple correspondence analysis (MCA) (Lê, Josse, and Husson 2008), and visually summarized the analysis with UMAP (**figure 2** and see interactive media from **supplemental file 1**) (McInnes, Healy, and Melville 2018). We used the resulting UMAP graph coordinates to perform density based clustering with HDBSCAN (McInnes and Healy 2017), which resulted in identification of 10 well-defined clusters capturing about 80% of the tumor samples. To capture the remaining samples into one of these 10 clusters we used kNN (see **Supplementary Methods** for details on clustering methods).

**Figure 2:**
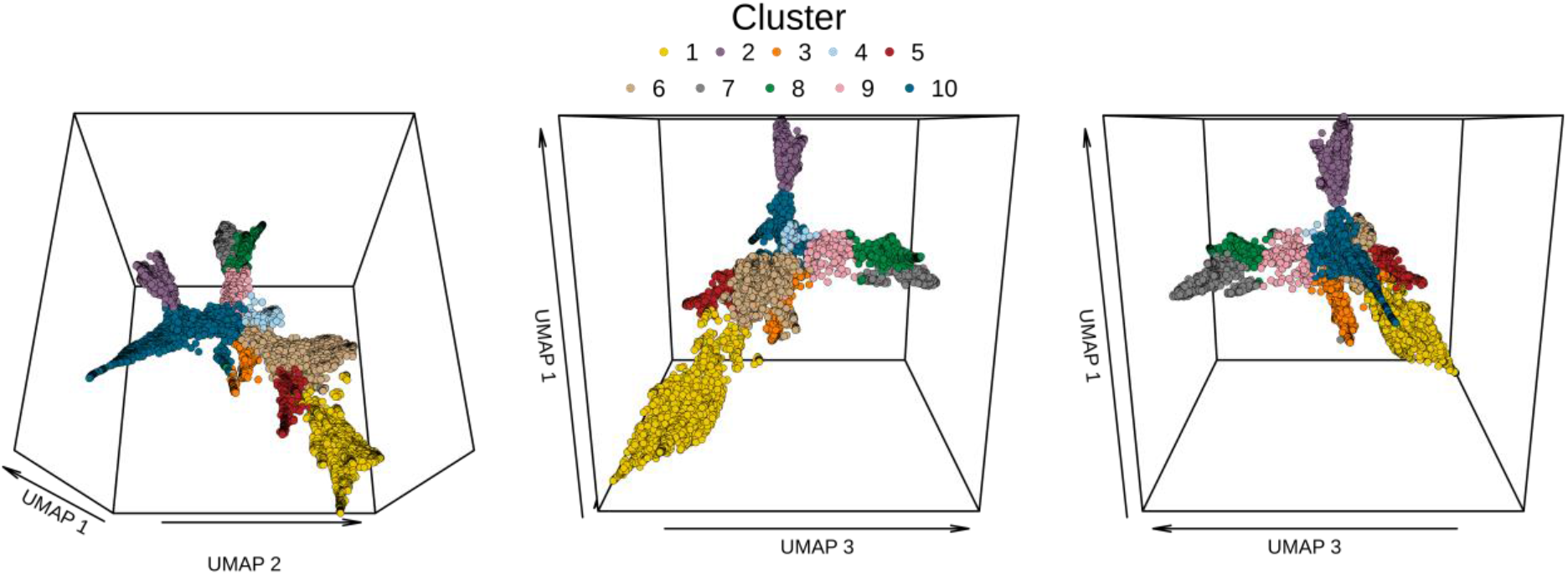
Clustering of tumor samples. Different rotational perspectives of the same MCA-based UMAP projection in three-dimensional space. Each dot corresponds to a tumor sample. The same colors indicate the tumor’s cluster identity throughout this manuscript.

#### Independence from tissue-of-origin

Having defined tumors in terms of their pathway disruption profile, we sought to understand whether different cancer types segregated into one or more predominant classes. To our surprise, most cancer types were not heavily biased in one class, and all well-represented cancer types had tumors in every class (see **figure 3A** and full tumor profiles in **supplementary figure S1**, see also interactive media from **supplementary file 1**), suggesting that, in principle at least, our pathway-disruptions identify clusters of molecular pathology largely independent of tissue-of-origin. As an example of one type of cancer that does have a biased pathway profile, pancreatic adenocarcinoma (PAAD) was predominantly found in class 8 (**figure 3A** and **supplementary figure S1** and **supplementary file 1**). But even PAAD comprises tumors from the nine remaining classes, meaning that patients suffering these tumors have potentially different underlying molecular pathologies.

**Figure 3:**
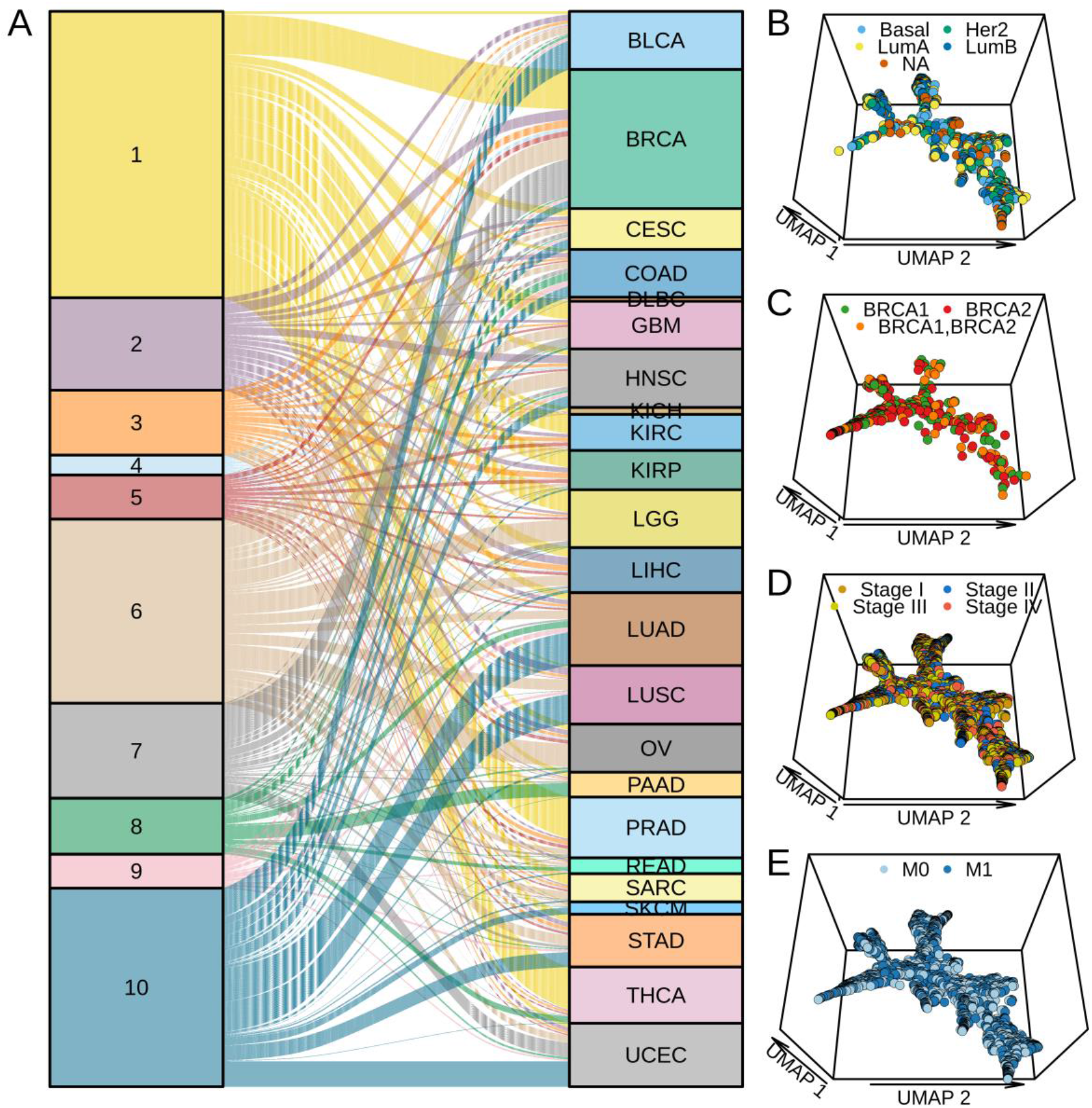
Pathway-based clustering independent of tissue-of-origin. A) Sankey plot showing correspondence between cancer type and cluster identity. B) Projection of breast cancer subtypes onto the UMAP. C) Projection of BRCA1/2 somatic mutation onto the UMAP. D) Projection of tumor stage onto the UMAP, regardless of cancer type. E) Projection of metastatic status onto the UMAP. Abbrevs: M0 = non-metastatic tumors, M1 = metastatic tumors.

#### Independence from molecular and histological subtype

Many cancers have molecular or histological subtypes defined based on gene expression or other -omics profiles or pathology lab results. These subtypes often have different standards of care owing to different overall drug sensitivity (or other factors). If the histological subtypes represent true molecular phenotypes, one predicts that histological subtypes should segregate with our pathway-based clusters, therefore providing support for the clusters as proxies for molecular pathology sub-typing. To our surprise, we find a similar result to the previous analysis of cancer types projected onto the UMAP of pathway disruptions. To illustrate this, we projected annotations for each of the breast cancer subtypes, composed of Triple-negative/Basal-like, Her2 positive, normal-like, and luminal A and B subtypes onto the UMAP. These are among the most heavily studied molecular subtypes in cancer, which each have different prognoses and standards of care. We did not observe any exclusive segregation by pathway for these subtype annotations (**Figure 3B**). We also projected histological subtype data for the remaining cancers (see **supplementary figure S2** for the full set of projections and see interactive media from **supplemental file 1**); we find that the subtypes, though often biased towards one or more classes, are almost never exclusive. We interpret these data in aggregate to mean that our pathway disruption classes do not correspond to previously identified molecular subtypes within the parent cancer type.

#### Independence from drivers of genome instability

There are several well-known familial cancer-causing mutations that have been interrogated extensively for differences in basic biology, survival and treatment outcomes. However, the functions of these genes are related to risk factors such as genome stability generally, proof-reading and DNA damage repair, and telomere length. *BRCA1/2* genes for example are key for DNA double-stranded break repair (Moynihan et al. 1999; Davies et al. 2001) and germline mutations in these genes confer elevated risk for breast, prostate and ovarian cancers. The mechanism of risk is thought to involve loss of heterozygosity resulting in loss of the wildtype, functional allele (Gudmundsson et al. 1995), so we projected the somatic mutations for *BRCA1* and *BRCA2* genes onto the UMAP, but did not observe segregation of these mutations into specific clusters (**figure 3C** and see interactive media from **supplemental file 1**). We made similar projections for the mismatch repair (MMR) genes *MSH2, MSH6, MLH1, MLH3, PMS1* and *PMS2; BRIP1, RAD51, CHEK2* and *APC*. None of these genes except for *APC* exhibited any remarkable specificity with respect to cluster assignment (**supplementary figure S3**). To look at other risk factors such as maintenance of DNA methylation levels and telomere length, we projected somatic mutations of the *TET2* and *TET3* genes, plus *TERT, TEP1*, and *DKC1*, and observed similar lack of segregation by cluster (**supplementary figure S3**).

#### Independence of stage, mutation count and mutation profile

Tumor staging is based on physico-pathological criteria, including tumor diameter, which can vary greatly in importance between different tissues. Stage is used clinically as a proxy for advancement toward a more deadly state and metastasis. Given these assumptions, it is possible that more advanced tumors have common pathway disruption profiles. The UMAP, which features a series of lobe-like structures on a common backbone of tumor samples (**figure 2**) could in principle reflect progression through a series of stages. The backbone starts with a cluster of tumors (class 1) that has the fewest point mutations and culminates in a cluster (class 10) which has nearly every pathway disrupted (**figure 4**). However, outside of class 10 we don’t observe an obvious trend in the overall mutation burden across the backbone of the UMAP. Nonetheless, to test the hypothesis that the molecular-pathway disruption clusters represent advancement through stages, we projected staging data onto the UMAP. Similar to tissue of origin and other categories of tumor, we did not observe any bias among the stages to specific pathway disruption clusters (**figure 3D**), suggesting that stage is not a contributing factor to cluster identity.

**Figure 4:**
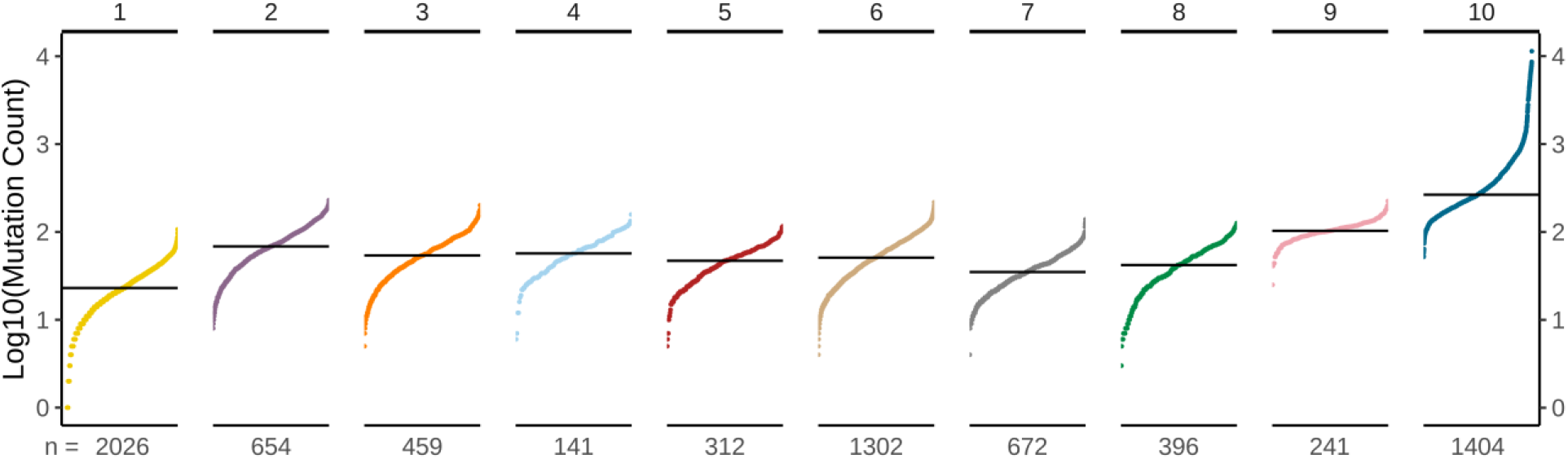
Somatic mutation frequencies for tumors in each class. Vertical axis shows log mutation count, horizontal axis is cluster identity. Each dot represents an individual tumor sample, ranked lowest to highest by mutation count. The median mutation count in each cluster is indicated by the horizontal line.

Finally, as a measure of tumor advancement, metastasis is the condition in which certain phenotypic criteria are met. These phenotypes include loss of differentiation, cell-cell contacts, epithelial to mesenchyme transition, immune system evasion and tissue invasiveness (Hanahan and Weinberg 2011). To determine whether any of our clusters correspond to an especially advanced stage of cancer across tissue types, we projected the metastasis data onto our UMAP, and surprisingly we observed an even distribution of metastases across classes (**figure 3E**). This final observation suggests that our pathway-disruption classification is dependent on particular combinations of gene mutations affecting different pathways that can each give rise to advanced stages of disease and metastasis, regardless and independent of overall mutational burden.

### Tissue specific genes define cluster membership

In order to identify pathway disruption enrichment across all cancers (pan-cancer), we created a list of pathway disruptions with percent mutated samples and top genes (**supplementary table ST1**). As expected, these analyses reveal the broad importance of many well known pathways that are disrupted in cancer, including “PIP3 activates Akt signaling” (77% of samples), “MAP1K/MAP3K signaling” (70% of samples), “Mitotic G2-G2/M phases” (67% of samples), “Cellular senescence” (64% of samples), “G2/M Checkpoints” (62% of samples), *etc*.

To discover what pathways are most important for clustering, we calculated percent enrichment for each pathway *within cluster* relative to all other clusters combined (see methods) and ranked pathways for each cluster from highest to lowest enrichment. We visualized the enrichment as a heatmap (**figure 5A**). Using this approach, we identified twenty-five pathways highly enriched (enrichment score ? 0.3, 95% confidence; see methods) in cluster 2, nine pathways enriched in cluster 3, twenty-four pathways in cluster 4, ten pathways in cluster 5, ten pathways in cluster 6, twenty-four pathways in cluster 7, twenty-four pathways in cluster 8, ten pathways in cluster 9, and no pathways for clusters 1 and 10 (**supplementary table ST2**). Clusters 7, 8, and 9 had several pathways in common. To explore the specific pathways marking each cluster, we projected disruptions for each of the 377 pathways onto the UMAP (**supplementary figure S4** and see interactive media from **supplemental file 1**). Clusters 3 and 5 were distinguished by metabolic pathways including RNA and protein biosynthesis (**supplementary figure S4**). Similarly, cluster 4 was distinguished by mutations affecting regulation of DNA and histone methylation (“DNA methylation”, “PRC2 methylates histones and DNA”, and “Nucleosome assembly”). Clusters 7-9 have in common mutations in extracellular, intracellular, and immune-related signaling pathways (see **figure 5B** and **supplementary figure S4**). Cluster 2 had the highest pathway enrichment levels of the three, having mutations in hedgehog signaling, “β-catenin degradation”, “cellular response to hypoxia”, “regulation of cell cycle” and “apoptosis” among others.

**Figure 5:**
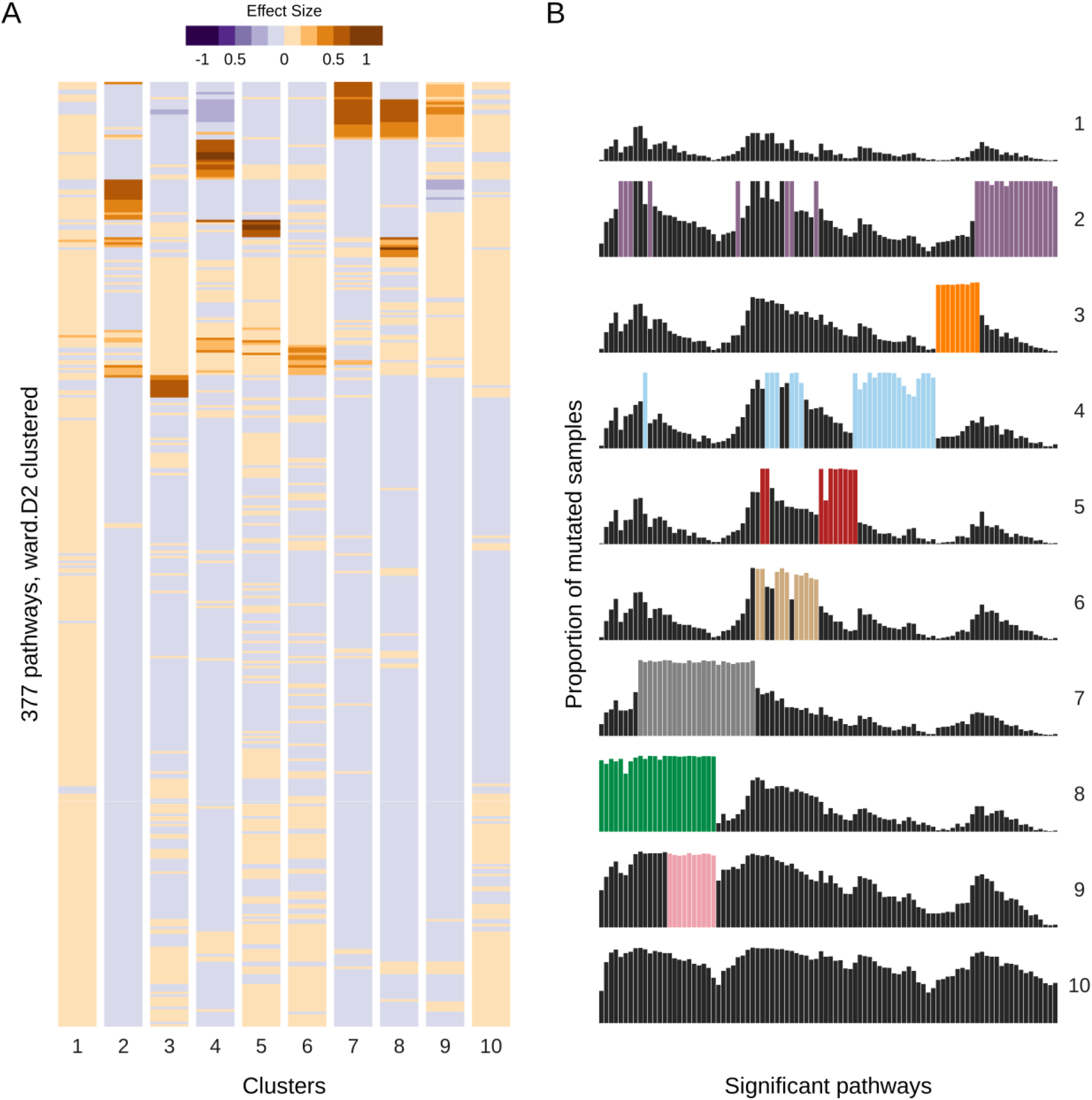
Pan-cancer enrichment of pathway disruptions. A) Heatmap shows the relative enrichment of each pathway (rows) within each numbered cluster (columns). Effect size is displayed as colors representing percent enrichment. B) Proportion of mutated samples in the each significant pathway (columns; union set of pathways with effect size ? 0.30 in each cluster) within each numbered cluster (rows).

Prior efforts to extract common signatures from pan-cancer datasets met with difficulty in distinguishing tumor samples from their tissue-specific-omics data signatures. Given our pathway-disruption based clustering, this raises the question, are tumor phenotypes driven entirely by common driver genes, or by “silent” tissue-specific effectors (*i.e*. too few samples to detect above statistical significance thresholds), or a combination of both? To answer this question, we compared top pathway genes for each cluster relative to the TCGA background to find differentially mutated genes. We ranked odds ratios and selected the top ten enriched and depleted genes (pvalue < 0.01) for each cluster (**Figure 6**; odds ratios plot). Clusters 7 and 8, which shared multiple enrichment in signaling pathways, are largely driven by mutations in *PI3K* and its orthologs and *Ras* genes, respectively (compare *PIK3CA* and *KRAS* panels of **Supplementary figure S5** and see interactive media from **supplemental file 1**). Interestingly, cluster 9, which also shared multiple enrichment in signaling pathways with clusters 7 and 8, is enriched for both *PIK3CA* and *KRAS*. Clusters 3 and 5, defined by enrichment in metabolic pathways, had mutations in ribosomal proteins and nuclear pore complex, respectively. Cluster 4 had mutations in genes responsible for nucleosome structure. Cluster 2 had mutations in proteasomal subunit genes involved in protein degradation. We also observed that genes that were enriched for one cluster are depleted from others (*i.e*. is enriched in cluster 6, but depleted in cluster 7; *PIK3CA* is enriched in cluster 7, but depleted in clusters 3 and 8). Next, we investigated the proportion of samples per cancer type for the significant genes within a cluster (**Figure 6**; heatmap). Surprisingly, cluster-specific tumors were not predominated by one or more highly mutated genes across all cancers. Instead, when observing the mutation rate for these genes within samples that belong to a cluster, the mutation rate is heterogeneous across tumors by tissue origin (*e.g*. In cluster 4, CESC was enriched for *H2AFX*, OV was enriched for *HIST1H2BD*, and UCEC was enriched for *HIST1H2AC*). Even among the top most mutationally enriched genes within clusters there is no global pattern, indicating that our clusters are not driven by individual genes, but rather networks as a whole. Taken together, our data identifies a framework of cancer type-specific mutations associated with specific clusters.

**Figure 6:**
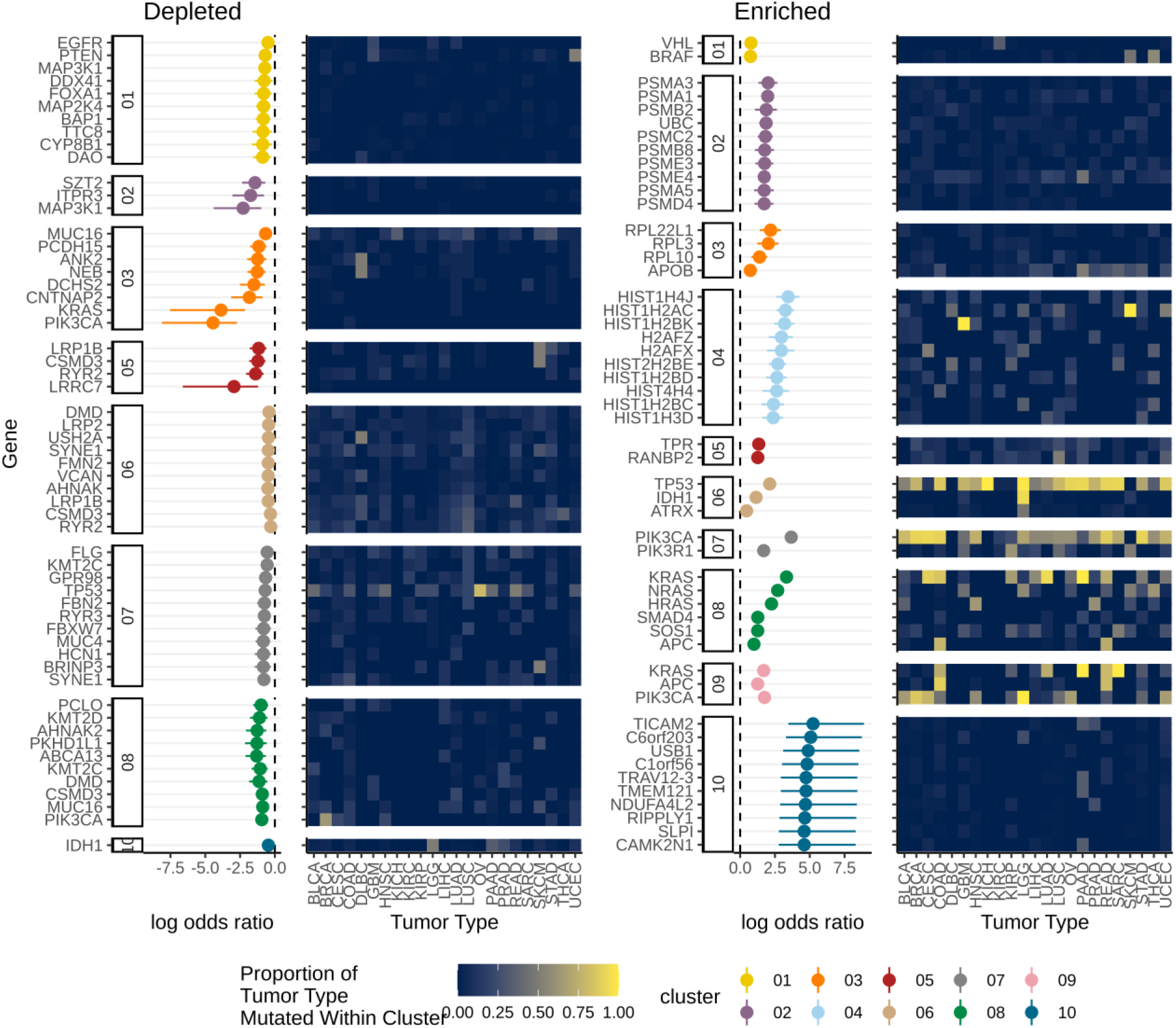
Gene level analysis reveals tissue-specific class signatures. Odds ratio plot; Column uses a logarithmic axis to represent odds ratio with a 95% confidence interval. Row represents the significant genes from each cluster. Each cluster was compared against the background (all other clusters) to find differentially mutated genes. Significant genes (pvalue < 0.01) were selected and limited to the top ten results for each clusters. Heatmap; Column represents the cancer type. Row represents the significant genes as described above. The heatmap shows the proportion of samples per cluster and cancer type that are mutated for each gene. Depleted significant genes (left). Enriched significant genes (right).

### Enrichment of pathways in metastasis is cluster-specific

Following the same logic we used to investigate cluster-specific enrichment of pathways we compared metastatic *vs*. non-metastatic tumors, as we find that metastatic tumors are distributed across all ten clusters (*e.g*. **figure 3E**). Using all non-metastatic tumors as background, we found very low levels of enrichment (< 10%) in a handful of pathways. We reasoned that the individual clusters might be too different to detect global metastasis enrichment signal given the small sample size (n = 215 metastatic tumor samples).

Therefore, we calculated cluster-specific enrichment in metastatic tumors and found a total of 31 enriched pathways (significant with enrichment score ≥ 0.3) across all clusters (**table 1**). A number of enrichments were found in multiple clusters represented by pathways that were already shown to be enriched in a neighboring cluster. For example, “Signaling by *PTK6”* is enriched in non-metastatic samples of cluster 8 (see **supplementary figure S4**) but not in 7 and 9. This pathway is enriched in metastatic tumors of clusters 7 and 9 (*p* < 10^−3^, **table 1**). This is also true of “Erythropoietin activates *RAS*”, which is enriched in non-metastatic tumors of cluster 8 (**supplementary figure S4**) and also in metastatic tumors of clusters 7 and 9. Cluster 4 metastatic samples were enriched for “Fc epsilon receptor (FCERI) signaling”, a key neutrophil pathway, which is also specific to clusters 2, 7 and 8 non-metastatic tumors. Thus, metastatic-enriched pathways from one cluster are often enriched in non-metastatic tumors of other clusters.

**Table 1:**
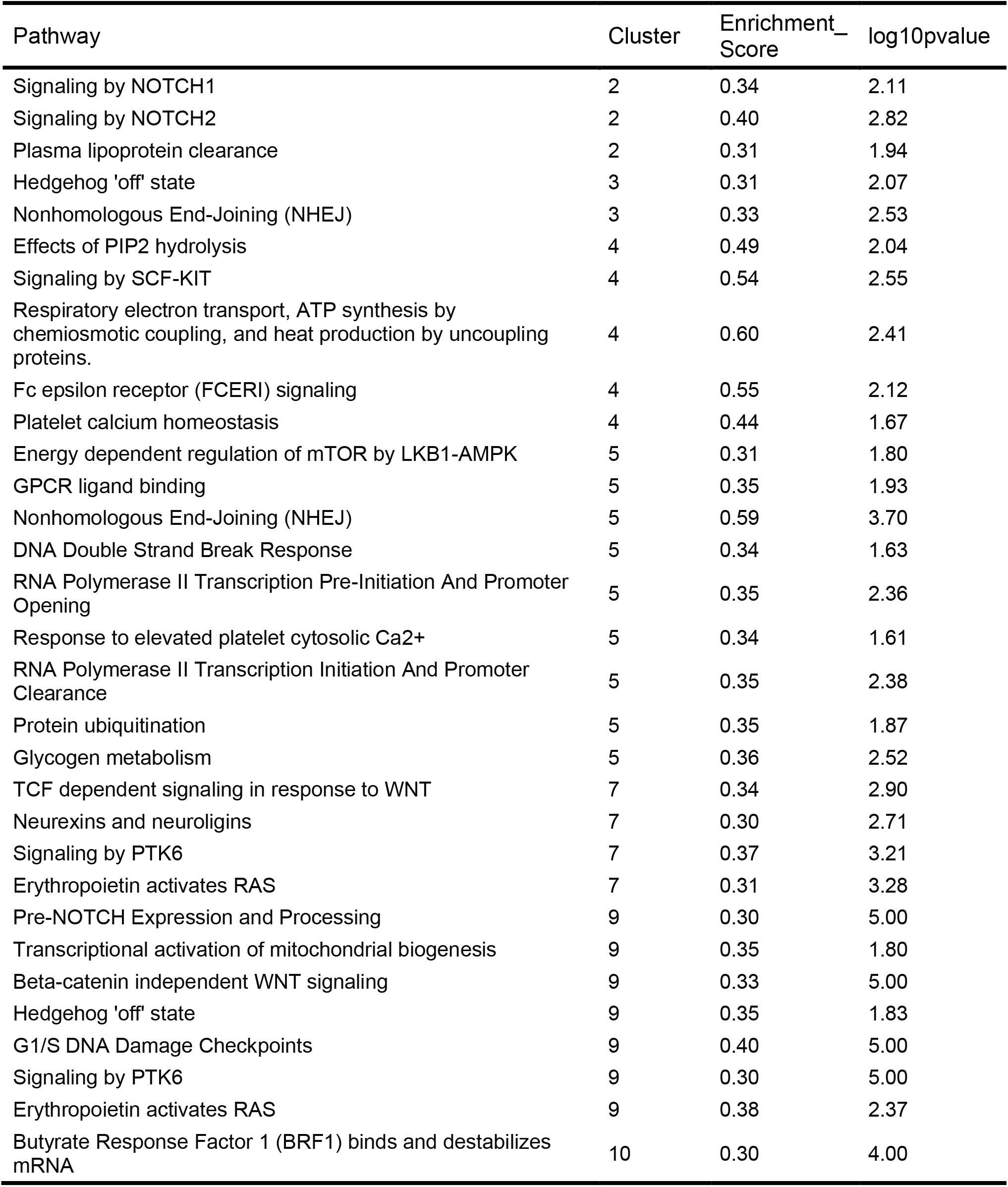
Cluster-specific enriched pathways (effect size ≥ 0.30) in metastasis.

| Pathway | Cluster | Enrichment_<br>Score | log10pvalue |
| --- | --- | --- | --- |
| Signaling by NOTCH1 | 2 | 0.34 | 2.11 |
| Signaling by NOTCH2 | 2 | 0.40 | 2.82 |
| Plasma lipoprotein clearance | 2 | 0.31 | 1.94 |
| Hedgehog 'off' state | 3 | 0.31 | 2.07 |
| Nonhomologous End-Joining (NHEJ) | 3 | 0.33 | 2.53 |
| Effects of PIP2 hydrolysis | 4 | 0.49 | 2.04 |
| Signaling by SCF-KIT | 4 | 0.54 | 2.55 |
| Respiratory electron transport, ATP synthesis by chemiosmotic coupling, and heat production by uncoupling proteins. | 4 | 0.60 | 2.41 |
| Fc epsilon receptor (FCERI) signaling | 4 | 0.55 | 2.12 |
| Platelet calcium homeostasis | 4 | 0.44 | 1.67 |
| Energy dependent regulation of mTOR by LKB1-AMPK | 5 | 0.31 | 1.80 |
| GPCR ligand binding | 5 | 0.35 | 1.93 |
| Nonhomologous End-Joining (NHEJ) | 5 | 0.59 | 3.70 |
| DNA Double Strand Break Response | 5 | 0.34 | 1.63 |
| RNA Polymerase II Transcription Pre-Initiation And Promoter Opening | 5 | 0.35 | 2.36 |
| Response to elevated platelet cytosolic Ca2+ | 5 | 0.34 | 1.61 |
| RNA Polymerase II Transcription Initiation And Promoter Clearance | 5 | 0.35 | 2.38 |
| Protein ubiquitination | 5 | 0.35 | 1.87 |
| Glycogen metabolism | 5 | 0.36 | 2.52 |
| TCF dependent signaling in response to WNT | 7 | 0.34 | 2.90 |
| Neurexins and neuroligins | 7 | 0.30 | 2.71 |
| Signaling by PTK6 | 7 | 0.37 | 3.21 |
| Erythropoietin activates RAS | 7 | 0.31 | 3.28 |
| Pre-NOTCH Expression and Processing | 9 | 0.30 | 5.00 |
| Transcriptional activation of mitochondrial biogenesis | 9 | 0.35 | 1.80 |
| Beta-catenin independent WNT signaling | 9 | 0.33 | 5.00 |
| Hedgehog 'off' state | 9 | 0.35 | 1.83 |
| G1/S DNA Damage Checkpoints | 9 | 0.40 | 5.00 |
| Signaling by PTK6 | 9 | 0.30 | 5.00 |
| Erythropoietin activates RAS | 9 | 0.38 | 2.37 |
| Butyrate Response Factor 1 (BRF1) binds and destabilizes mRNA | 10 | 0.30 | 4.00 |

### Pathway disruption clusters vary in short-term prognosis of survival

If our molecular pathway disruption clusters represent biological states distinct from tissue of origin, they may have different prognoses within cancer types or across all cancers. These analyses are limited by confounding factors of age and stage at diagnosis, sex, ethnicity, and tissue-specific disease progression. To explore these ideas, we used Bayesian inference to test models (see Methods and **Supplemental Methods** for greater detail) of survival using public longevity data from the CDC, accounting for age, gender and ethnicity. Our model estimates the effect of cancer type and cluster-specific cancer effects independently, resulting in a cancer and cluster-specific estimate of the aging rate multiplier, *k*.

To understand the model, we show in **Figure 7A** simulated survival time curves for four related groups. We start with a cohort of women aged 30 and show their expected survival (light blue). By setting their cancer rate multiplier to *k* = 2 we can simulate the effect of a moderately deadly cancer (yellow). Compare this to a randomly selected group of women ranging in age from 30 to 70 years old (dark blue). Immediately the survival curve changes due to the mixture of ages, without any malignancy. Adding a *k* = 2 malignancy further reduces expected lifespan (red). We can see that for different tissues, simply due to the change in distribution of age-at-diagnosis, we should expect equally deadly cancers to have *different* survival curves. The model takes this into account and the aggressiveness of the cancer can be estimated accurately without the confounding effect of age.

**Figure 7:**
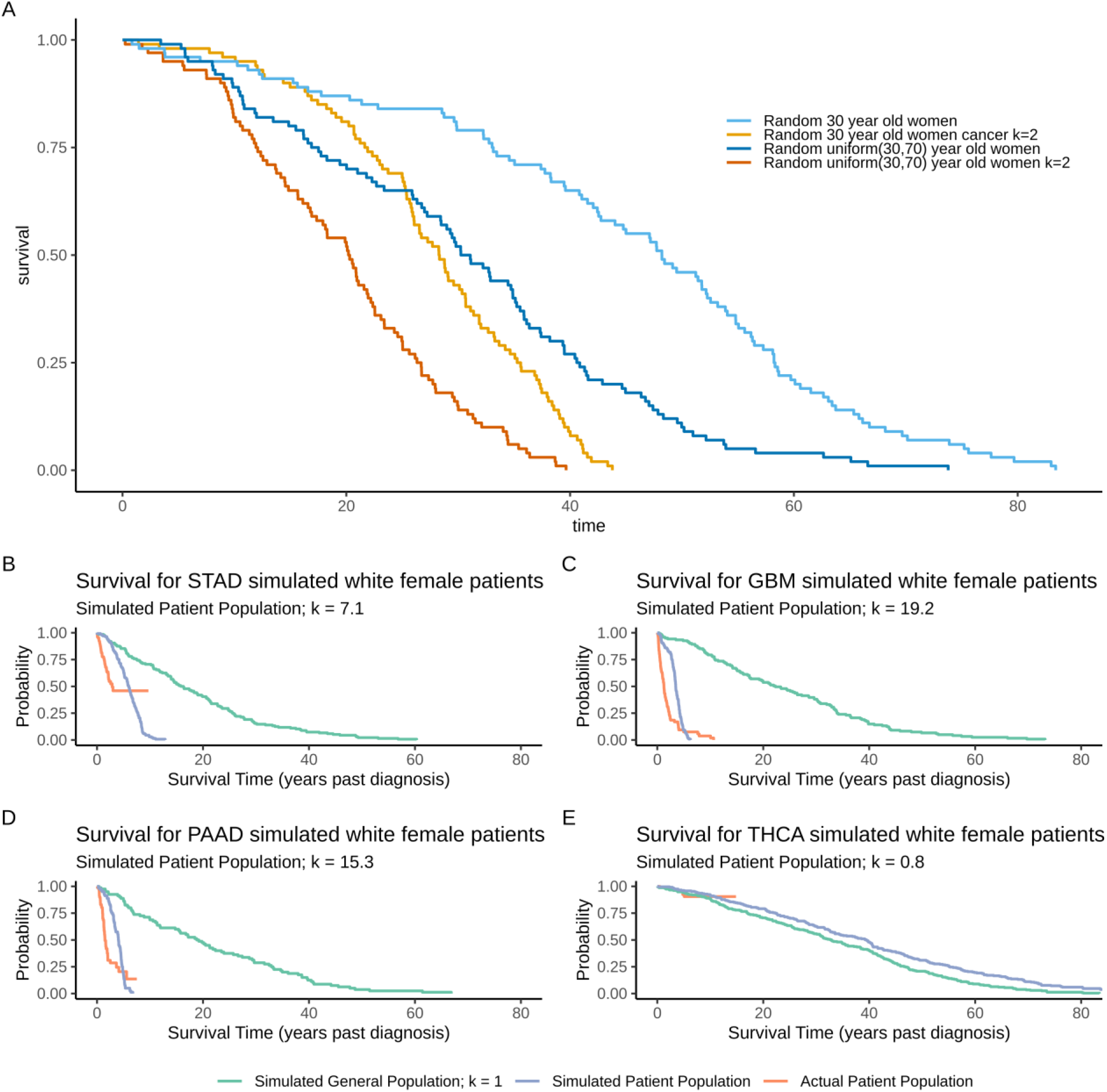
Comparison of different cancers, simulated and actual results results. To illustrate the risk model, we compare survival of four simulated cohorts. A) A group consisting of randomly selected 30 year old women (light blue) vs. an equal sized group of 30 year old women with cancer, k = 2 (light orange). To illustrate how the mix of ages alters the survival curve, we compare a randomly selected group of people with ages uniformly distributed from 30 to 70 (dark blue) and a randomly selected group with uniformly distributed ages and cancer k = 2 (dark orange). B,C,D,E) We compare a randomly selected group with correct age distribution for the given cancer (turqoise) to the model predicted survival for the given average k (blue) and the actual survival for the patients in our dataset (orange). As can be seen, Stomach Cancer (STAD), Glioblastoma (GBM), and Pancreatic Cancer (PAAD) are all very deadly. The modeled results fit the general trend for the actual patients. Thyroid Cancer (THCA) on the other hand has very little if any effect on our estimates of life expectancy, in fact there may be a slight benefit which may be associated with a bias in diagnosis towards patients who are more health conscious, or higher socioeconomic status, with more access to care compared to random population members.

We found that cancer types, as expected, have a range of prognoses relative to the general population. For example in **Figure 7B-E** we see three particularly deadly cancers (Stomach:STAD, Glioblastoma:GBM, and Pancreatic:PAAD), and one cancer where diagnosis apparently decreases risk relative to background (Thyroid: THCA). Cancers with posterior probability for relative risk of less than 1 such as THCA should be interpreted carefully. This Bayesian model is a model for a state of information. The information that a person is diagnosed with cancer may lead us to expect that they will live a shorter time than the general population of matched age (k > 1), or a longer time than the general population of matched age (k < 1). One mechanism for a shorter result is that the cancer is aggressive and we can expect it to rapidly injure the body, causing death. One explanation for longer results may be that the cancer is relatively mild, and therefore diagnosis is potentially an informational signal that the patient is health conscious, with the comparison group having more people whose cancers go undiagnosed. It’s important to note that the diagnosis can increase our expectation of life relative to the comparison group, even if it decreases the expectation of life of the individual relative to the counterfactual where they did not have cancer.

The degree to which a given cancer accelerates aging can be determined by the *k_tls_* multiplier (**Figure 8**) multiplied by the cluster modifier *k_cl_* (**Figure 9**). Looking at the tissue specific multiplier, the least deadly was prostate cancer (PRAD), and the most deadly was glioblastoma (GBM) which has a risk multiplier of between 15 and 22 relative to the background risk of death in the population. This undoubtedly is influenced by the fact that relatively young patients are affected by GBM and that it is extremely deadly. Among the deadliest cancers outside of GBM were stomach (STAD), melanoma (SKCM) and pancreatic (PAAD) cancers. Apart from these top 4 cancers which ranged from 6-22 in relative risk of age-adjusted death, the remaining cancer types ranged from about 1 to 5 in magnitude.

**Figure 8:**
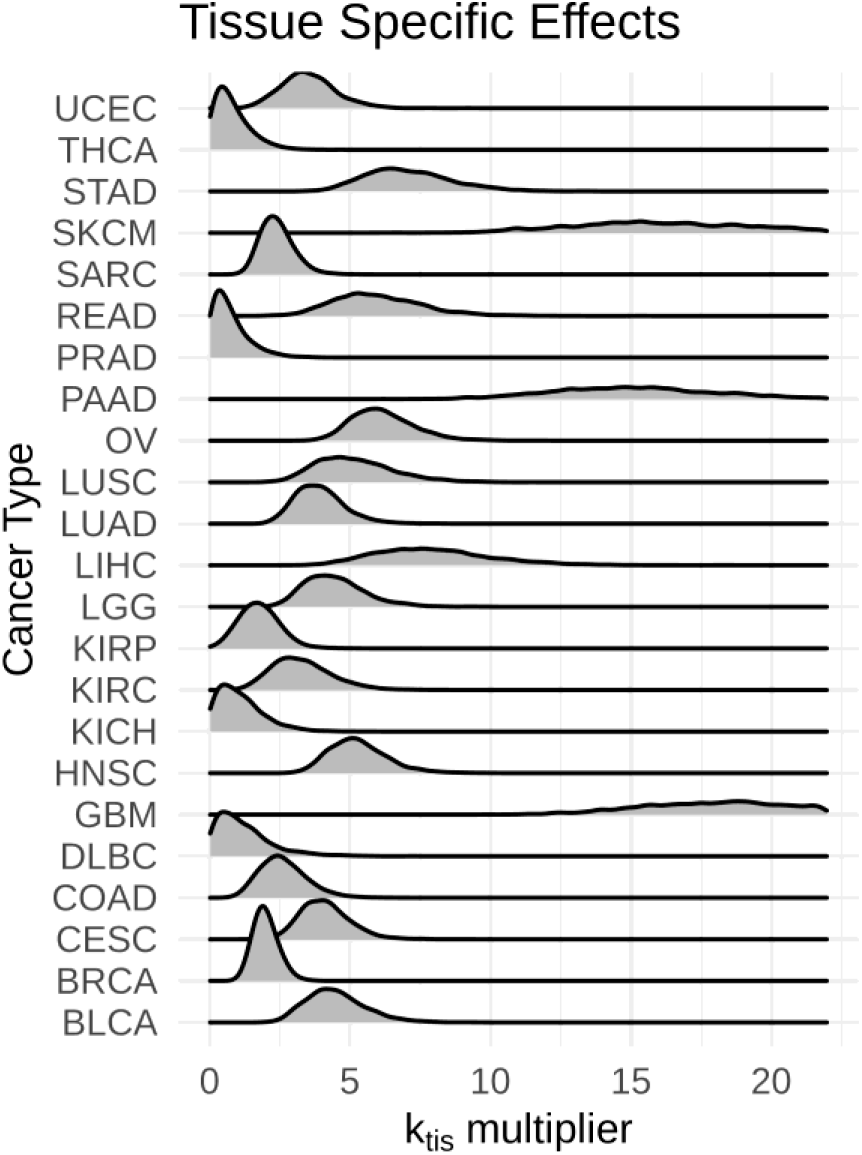
Tissue specific effects. Effects are compared between cancer type by showing the probability density for the posterior value of the *k_tis_* multiplier. Larger values correspond to decreased age-independent life expectancy.

**Figure 9:**
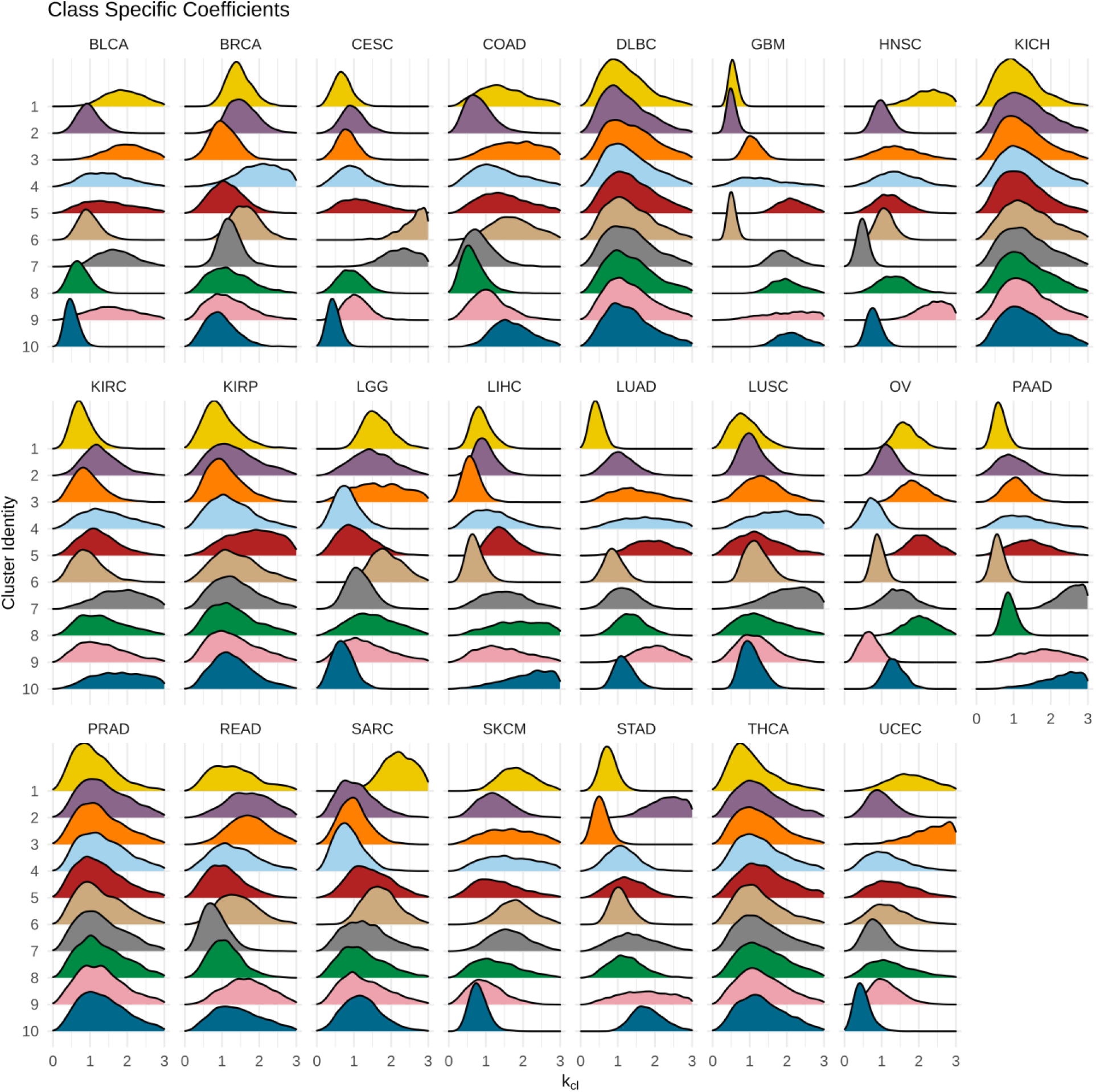
Cluster specific *k_cl_* values for each cancer type. The overall *k* value is the product of *k_tis_* × *k_cl_*. The cluster-specific value represents the relative aggressiveness of each cluster within the cancer type. For example clusters 1, 2, and 6 Glioblastoma (GBM) are apparently much less deadly than other types of GBM. A similar situation can be seen with Pancreatic Cancer (PAAD) where 1,6, and 8 are less deadly compared to clusters 7, 9, 10. For some cancer types there are few data points and little ability to estimate values with precision (DLBC and KICH for example).

Our estimates of tissue-specific cluster effects, in contrast, ranged from less than 1 up to about 3 or 4, reflecting that some clusters are either less deadly or more deadly than other clusters within each cancer type (**Figure 9**). A cluster-specific rate of 1 represents the typical rate for this tissue type. For several cancers (*e.g*. PRAD, kidney chromophobe (KICH), diffuse large B-cell lymphoma (DBLC), thyroid cancer (THCA)) the posterior estimates are largely indistinguishable from the prior, reflecting that either there were too few mortalities in the data to make an estimate (as expected for PRAD and THCA) or two few samples, period. We did not observe cluster-specific trends that held true across cancer types, which could result from different cancers having different standards of care for example. In support of this interpretation, we also tested a factored model which considered both cancer type and each cluster independent of cancer type (not shown). Though we were able to successfully fit this model, it is a special case of the more general model where cluster and tissue independently affect longevity, and there is no reason to believe that cluster specific effects would necessarily remain constant across tissue types given how widely the patients vary across tissue types, drug and surgical treatments.

## Discussion

### Classification of tumors independent of tissue-of-origin

One of the biggest hurdles in cancer research is the sparsity of data; ~20,000 protein-coding genes is comparable with the number of tumor samples, even with multiple mutations per sample. This also does not account for the fact that ~30,000 non-coding genes and hundreds of thousands of non-coding regulatory elements exist with oncogenic or tumor suppressor activity (Dees et al. 2012; Tamborero, Gonzalez-Perez, and Lopez-Bigas 2013; Lawrence et al. 2014; Kumar et al. 2015; Jiang et al. 2019; Zhao et al. 2019; Zhang et al. 2020). Hence, we sought to simplify the problem by employing a “knowledge-base driven analysis” (Khatri, Sirota, and Butte 2012), investigating cancer as a disease of basic cellular and biochemical pathways, which we accomplished by translating gene-level mutations into pathway level disruptions. Our approach differs from previously described methods (Creixell et al. 2015) in that we chose to focus on the pathway (defined in the methods) as the unit of disruption instead of the gene, where individual mutations may be sufficient to disrupt or misdirect pathway activity. Rather than evaluating the burden of mutations on each pathway within or across tumors, which can be biased by the number of genes or the coverage of pathway-linked genes in the genome, we scored each pathway in binary fashion, allowing one mutation or multiple mutations to count equally for each sample.

To our knowledge, this approach has not previously been attempted in spite of its relative simplicity. We limited our analysis to mutations with likely deleterious effects, including missense, nonsense, frame-shift, stop-loss, untranslated region and splice site mutations in genes that are actively expressed in each cancer type, thus avoiding bias from transcription coupled repair. Our method of filtration differs from the “rank-and-cut” method of (PCAWG Consortium 2020) but represents a reasonable attempt to account for the same biases. We also filtered genes on the basis of pathway membership in biochemical pathways, excluding curated gene sets related to diseases, syndromes, or classes of proteins with shared catalytic activity or conserved domains which are potentially problematic (Khatri, Sirota, and Butte 2012). We chose this approach to limit redundancy and exclude biologically unrelated collections of genes. Our pathway-centric analysis allows us to circumvent the problem that some cancer-promoting mutations likely occur at such low frequencies as to be indistinguishable from background.

The hypothesis that cancer is the result of dysfunction from a limited number of basic cellular processes common to all living human cells was introduced and later expanded on in a pair of essays by Hanahan and Weinberg (Hanahan and Weinberg 2000, 2011). In this view, all tumors regardless of tissue of origin acquire six essential phenotypes; sustaining proliferative signaling, evading growth suppressors, replicative immortality, evading apoptosis, supportive angiogenesis, tissue invasion and metastasis (Hanahan and Weinberg 2000). Advancements in the field over the ensuing decade led to the inclusion of new tumor promoting processes, namely deregulation of cellular energy cycle, tumor promoting inflammation, evasion of the immune system, genome instability, and on top of these acquired traits a new appreciation for the concept of the tumor microenvironment and its contribution to the evolution and promotion of metastatic disease (Hanahan and Weinberg 2011).

An alternative hypothesis is that every tumor belongs to one of a large number of syndromes which are unique to each tissue-of-origin, that share some mechanisms and treatment strategies. Recent publication of TCGA consortium papers present a view largely, and surprisingly, consistent with this hypothesis (Hoadley et al. 2014, 2018). Perhaps owing to the intractable complexity of genomics, proteomics, and patient metadata in all its forms, the inescapable conclusion thus far is that tissue-of-origin remains the most important driver of tumor characteristics at every scale and by every measure. Our observations contrast with this view, and instead support an interpretation of publicly available data in which all tumors manifest one of a limited number of metastable phenotypes resulting from disruptions in basic pathways as represented by our clusters.

We attempted to account for our clusters in terms of more trivial explanations. For example, it could be that the clusters we identified are consistent with a signature of overall disease progression, such that successive, adjacent clusters on the UMAP projection exhibit increasing specificity. We identified no such trend in the number of mutations, the relative staging or metastasis; and each cluster instead was associated with unique combinations of pathways.

We find that most tumors segregate into one of ten biologically distinct clusters, regardless of the cancer type. Some cancer types are unevenly distributed, though we could not identify any cancers that were exclusive to a single tumor cluster. Only cancer types with the fewest samples were found to be absent from one or more clusters at all. Somewhat surprisingly to us, this finding extends to histological subtypes of breast, head and neck cancers, leukemias, *etc*. This result implies that many, if not most, histological subtypes could reflect differences in cell-of-origin, rather than fundamental differences in proximal cause. The four major subtypes of breast cancer correspond to histological and molecular expression profiles that define them and how they respond to experimental stimuli (Prat and Perou 2010). It has been hypothesized that differences in the molecular regulators of development in the precursor cell types present in the breast epithelium drive the various histological phenotypes (Skibinski and Kuperwasser 2015; Zhang, Lee, and Rosen 2017). Consistent with this view we found that breast tumor samples of the Luminal A subtype were heavily biased toward membership in clusters 1 and 7, and basal tumor samples were biased toward cluster 6. However, both subtypes also contained samples in every other cluster (without exception), and Luminal B and Her2 positive samples are distributed across clusters. We interpret these data to mean that inherited cell-of-origin molecular profiles may predispose certain precursor cell types within the breast epithelium to forming tumors of one cluster or another but are not determinative; this view is compatible with the previously stated hypothesis but opens the way for a more granular view of individual tumors. It would be surprising if we did not observe bias for some cancers and subtypes amongst our classes, since some treatment regimens have greater efficacy for patients of a given cancer or histological subtype (Prat and Perou 2010). Nonetheless, the basis for some tumors being treatment-refractory in spite of receiving the standard of clinical care for diagnostic markers remains elusive. Doubtless some of this is due to chance events, as tumors can metastasize and remain dormant years before they are detected at distal sites, or resistant clones may have already arisen at undetectable levels (Hanahan and Weinberg 2011), but our analysis suggests the possibility of identifying more informative molecular, histological or cellular subtypes that could form a basis for future stratification of patients into different precision treatment regimens.

### Tissue specific manifestation of pathway-centric disruptions

Our results illustrate how unique combinations of mutations in pan-cancer driver genes with tissue-specific pathway disruptions result in common categories when viewed at the level of the pathway knowledge-base. Top cancer driver genes (*e.g. PIK3CA* and *TP53*) are found in most of the clusters, in spite of the fact that they contribute to many cluster-specific pathways. This can only be explained as a result of other less common driver genes complementing the pathway disruption of the unique combination of driver genes that are disrupted in each tumor, and we speculate that many of these are relatively tissue-specific in terms of their sensitivity to mutation. Consistent with our observations, Colaprico et al. (2020) showed how tissue- and cancer-specific driver genes and druggable targets of pathway moduli can be discovered by machine learning on integrated multi-omics datasets. Marrying these approaches in future work will greatly enhance our understanding of tumor biology.

### Incompleteness of the pathway disruption data

Finally, we must remark on the limitations of our work exemplified in two of the clusters, 1 and 10, for which we did not find many distinctive associations with pathways. Cluster 1 had a relatively low proportion of mutated pathways across the board, although it is broadly enriched in many of the same tumor-promoting pathways common to the other groups. This likely does not reflect a true difference in stages of tumor evolution, as our data show clearly that this cluster is as likely to contain stage IV metastatic tumors as it is to contain those of stage I. This cluster likely represents a group of tumors with aberrations in methylation, key deletions, or other structural variants. Consistent with this, kidney chromophobe and thyroid cancers have high proportions of structural variations *vs*. other variant types (PCAWG Consortium 2020) and are heavily skewed to cluster 1 membership. Likewise, cluster 10 represents a group of hyper-mutated tumors that harbor so many mutations that virtually no pathway is unaffected. It seems likely that a significant fraction of the “mutant” samples for each pathway are burdened with excess passenger mutations. This could be addressed with application of ever-increasingly sophisticated filtering of likely passenger mutations (*e.g*. Jaganathan et al. 2019; Sundaram et al. 2018). In the future, we hope to incorporate these other data into a comprehensive pathway-centric analysis as we have done here for point mutations and indels.

### Estimates of survival reveal pathway-dependent differences

By modeling CDC longevity data as a baseline risk function we showed that each of our pathway disruption clusters exhibit cancer-type specific effects on expected survival. Contrary to our initial expectation, we found that models assuming different survival based on cluster × cancer type fit better than models in which they are independent factors. However, considering that within each cancer type there are different clinical standards of care, and even within classes of drugs the preferred treatment can very between cancers, it makes sense that we observe tissue-specific cluster effects. Contrast the situation with ovarian *vs*. breast cancer, which are both hormonally driven cancers, for example. Ovarian cancer has but one main treatment axis, platinum, whereas breast cancer patients have a variety of treatment regimens based on molecular subtype and other factors. Standard of care for one cancer and cluster could produce different outcomes than the same cluster of a different cancer type. Unfortunately, given the diversity of drug classes and treatments (including non-chemical treatments such as surgery and palliative care), we lack sufficient power to explore these variables in the TCGA data. It is our hope that future studies will help to distinguish between treatment-specific effects on survival given different pathway disruption clusters.

### Implications for the evolution of cancer

Our findings imply that there are separate processes in the etiology of cancer that can be broadly thought of as general cancer promoting, cluster-specific mutations and metastasis. General cancer promoting processes must include factors that relate to genome stability and immortality, as “enabling characteristics” of the cancer phenotype (Hanahan and Weinberg 2011). Such pathways are disrupted in all or most of the defined clusters and are frequently the result of aberrations involving common driver genes such as BRCA1/2, MMR genes, mitotic checkpoints, cohesion complexes, *etc*. Cluster-specific evolution must involve the acquisition of disruptions to pathways that may individually be harmful (*e.g*. highly proliferative cells are more likely to senesce) but together produce more specialized cancer phenotype and increased fitness. Importantly, our observations do not imply the order in which these mutations should accumulate. This could be addressed in a future study by integrating our analysis with evolutionary analysis of clonality, drawing inferences from variant allele frequencies as in Gerstung et al. (2020). However, since many of the genes in the non-cluster-specific pathways involve the known driver genes, it is reasonable to hypothesize that these mutations promote or enable the subsequent acquisition of cluster-specific defects via random mutation and natural selection, thus producing the clusters we observed. In support of this, the pan-cancer analysis of whole genomes consortium (PCAWG) found that oncogenic driver mutations are highly enriched in early arising clones, whereas later arising clones have much greater diversity in driver mutations (Gerstung et al. 2020). Moreover, driver genes that are known to be responsible for discrete mutation signatures such as APOBEC, BRCA1 and BRCA2 produce mutational hotspots reflecting the varying selective pressures in different tissues (PCAWG Consortium 2020).

One of the drawbacks of bulk tumor whole genome sequencing data is the problem of tumor heterogeneity. Consortium samples are likely to contain contamination from support tissue, stroma, inflammatory cells, immune cells of the innate and adaptive immune systems, all potentially harboring cancer supporting mutations (Tripathi, Billet, and Bhowmick 2012). We think it will be instructive to explore these ideas in the context of single cell experiments.

### On metastasis as a convergence of phenotypes

We report that pathway enrichment in metastatic tumors across all clusters (pooled) yielded generally lower effect sizes and larger p-values than the cluster specific analysis, suggesting that signal is diluted when the samples are analyzed together, and supporting the view that metastasis has cluster-specific requirements. Since the number of metastatic samples is relatively low, this part of our analysis is likely underpowered and subject to expanded analysis with larger cohorts. The fact that most metastatic enrichment is cluster-specific and has a tendency to overlap with cluster-specific pathways from non-metastatic tumors of neighboring clusters suggests a mechanism of complementarity, in which newly acquired mutations result in similar clusters converging on one or more deadly phenotypes that capture most of the critical features of end-stage cancer. Thus, even at relatively low power, our analysis of metastasis further supports the pathway-centric approach as a fruitful way to uncover biologically meaningful differences between tumors against the noisy backdrop of tissue specific profiles. As a major caveat to this, we acknowledge that many therapies *do* target highly-specific driver genes, markers, and signaling pathways (*e.g*. TP53, EGFR or HER2), but understanding the broader context of the genetic background and pathway vulnerability of tumors containing such markers may aid in creating smarter combination therapies. We submit that when we discover the requirements of each cluster with respect to pathway disruptions and metastasis we may be able to target them therapeutically and prevent further adaptation.

## Materials & Methods

All code for producing the analyses and figures herein are included in this fully reproducible manuscript in R markdown format. R markdown files and all other models are available from our repositories on the distributed version control site, GitHub.

### Selection of pathways

To understand the molecular mechanism of cancer at a pathway level, we used Reactome (https://reactome.org/), a knowledge-based pathway database. The mapping files of ENSEMBL genes to pathways, pathway hierarchy relationships, and complexes to top pathways were downloaded from https://reactome.org/download-data. Using these data, we imposed pathway criteria to define basic cellular processes and biochemical pathways: (1) human-derived pathways (*e.g*. ‘Beta-catenin independent WNT signaling’ in ‘Signal Transduction’) (3) exclusion of pathways in the parent pathway: “Disease”, “Muscle contraction”, and “Reproduction” or pathway names that include any of the following keywords: “disease”, “defect”, “cancer”, “oncogenic”, “disorder”, “mutant”, “loss”, “infection”, “bacteria”, or “listeria”. While some of the excluded pathways have been shown to play an important role in cancer, they are highly specialized (*e.g*. “PI3K/AKT Signaling in Cancer”). Additionally, for most of the excluded pathways, a neutral version pathway of the pathway exists (*e.g*. “PIP3 activates Akt signaling”). Finally, we mapped the 18,577 Ensembl IDs from the TCGA dataset to the highly selected Reactome pathways. This operation produced a lookup table that consisted of 377 pathways mapped to 8,940 genes.

### Filtering genes

We filtered likely erroneous mutations due to transcription coupled repair. Our approach was to determine the status of each gene (*i.e*. expressed or not expressed) in each tissue type in order to exclude gene mutations low expressed enes. To do this, we obtained the TCGA RNA sequencing data adjusted for batch effect dataset (https://pancanatlas.xenahubs.net). Using the data, we removed the genes and tumor samples that were not included in our analysis, grouped the tumor samples by tissue type and computed the mean expression value for each gene. A minimum threshold of 10 transcripts per million was set for expressed genes based on an inflection point observed when plotting the mean expression values of genes ranked by expression in each tissue type. Genes that did not meet this threshold were considered not expressed. This operation produced a lookup table for gene expression status in each tumor sample for 18,127 genes.

### Class and stage-specific Enrichment Calculations

We calculated the enrichment of pathways in one set of tumors as the relative fraction of tumors (with estimated uncertainty) with a mutated gene in the pathway to all tumors, inclusive of the category of interest. To do this operation, we use the beta distribution to permute a posterior distribution on the fraction of tumors with a pathway mutated for each category based on the observed set, and compared this to the posterior obtained from the full set of tumors (all tumors) as the distribution of differences between all permuted samples. We considered a pathway enriched if the 95% range of credible differences thus obtained excludes 0 and the mean credible difference was greater than or equal to 30% enrichment, which excludes a large number of small differences that are not likely to be biologically relevant.

### Models of survival based on population-specific longevity data

We used Bayesian inference to explore the relative impact of cancer diagnosis on survival at the time of diagnosis. Our model assumes that at birth each person has a small initial probability per unit time to die, and that this probability grows exponentially in time at some baseline rate that matches the observed CDC data. The rate of growth in risk is assumed to jump to a higher level at diagnosis, which we take as a proxy for onset. This represents an accelerated aging type model. To fit the model, we constrain the baseline risks using the available CDC life tables (ftp://ftp.cdc.gov/pub/Health_Statistics/NCHS/Publications/NVSR/68_07/). We also constrain the cancer specific multiplier using the longevity data within the cancer tumor dataset. Our model estimates the effect of cancer type and class-specific cancer effects independently, resulting in a cancer and class specific estimate of the aging rate multiplier. A detailed description of the CDC longevity model and likelihood is provided in the **Supplementary methods**.

## Acknowledgements

We wish to thank David Van Valen, Kate Lawrenson, Simon Knott, and Megan Hitchens for critical reading of this manuscript, and Ivetth Corona for early discussions and feedback.

## Declarations

The authors have no conflicts of interest to declare. This work was supported by a grant from the Cedars-Sinai Precision Health Initiative.

## Notes

https://junkdnalab.shinyapps.io/PANCAN_supplemental/

https://github.com/dennishazelett/TTmanu

### Competing Interest Statement

The authors have declared no competing interest.

