## Supplemental Methods for "A molecular taxonomy of tumors independent of tissue-of-origin"

8/24/2020

### Supplementary Methods

#### Clustering

In order to classify tumors using this dataset, we used multiple correspondence analysis (MCA) (Lê, Josse, and Husson 2008). First, we determined the number of dimensions containing useful information by selecting the eigenvalue with the most explanatory power, using the average of 100 permutations of the data as baseline (**figure SM1A**). We then chose the maximum eigenvalue for which the p-value remained ≤ 0.05 (see cutoff in **figure SM1B**). Then we performed a UMAP analysis (McInnes, Healy, and Melville 2018), both in order to summarize the MCA graphically, and as a preprocessing step to boost the performance of density based clustering. The resulting map was notable for its lobed structure, with several reproducible projections regardless of random seed setting. A representative version of this 3D UMAP is shown in **figure SM1C**, rotated to enhance the visibility of the major features. Following this spatial mapping we attempted to define groupings of similar tumors within the spatial map using HDBSCAN, which performs hierarchical clustering and provides metrics of cluster stability and probabilities of cluster membership for each node (McInnes and Healy 2017). However, HDBSCAN is sensitive to several parameters; key for our analysis are the minimum number of tumor samples in a cluster that capture the maximum number of tumor samples, measured by probability of membership of ≥ 5% in at least one cluster. Thus, we created a score metric as the fraction of classified tumors with max probability < 5% in one cluster and chose a cluster size of 92 to minimize the score function (**figure SM1D**). HDBSCAN with these settings resulted in ten distinct high-density clusters which we then projected onto the UMAP (**figure SM1E**). This classified 6,038 out of 7,607 tumors but still resulted in a significant fraction of unclassified tumor samples. Since we ultimately wish to be able to classify any tumor using this scheme, we performed k Nearest Neighbors (kNN) analysis, which computes a similarity metric to every tumor in the set and then lets the *k* most similar tumors "vote" as to the identity of the query tumor sample based on their cluster labels. We set *k* to be the square root of the number of tumor samples (87). Using this method, we assigned cluster membership to the remaining tumors (**figure SM1F**).

*Figure SM1*

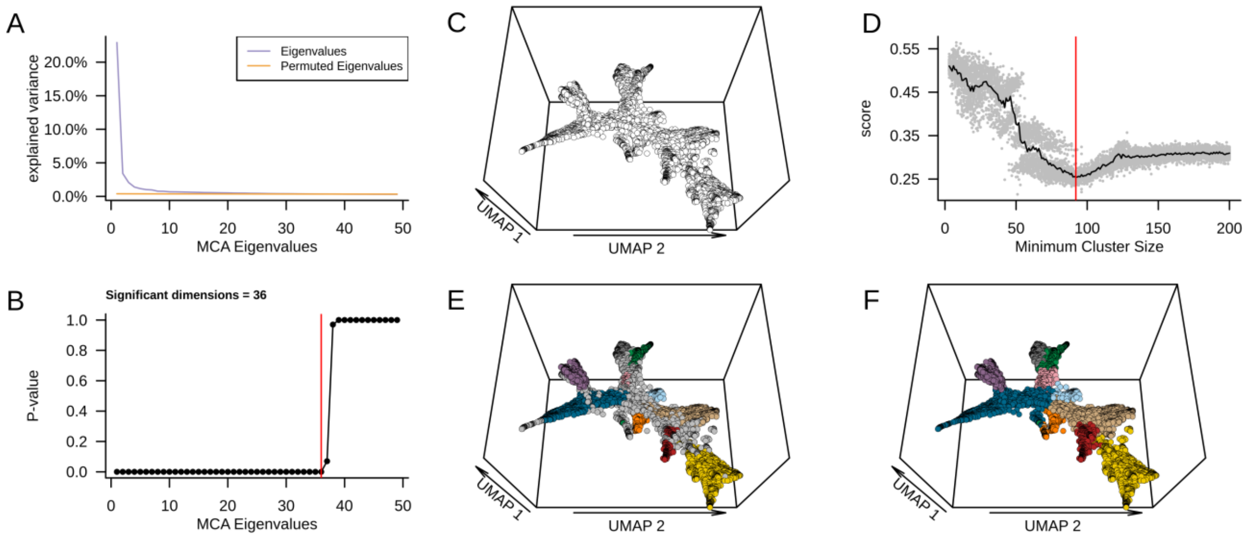


**Figure SM1: Method for tumor classification using pathway aberration data** A) MCA analysis was performed to determine the eigenvalue with the most explanatory power from the categorical data. Average of 100 permutations of the data was used as a baseline. B) The p-value of each eigenvalue was plotted and the max eigenvalue with a pvalue < 0.05 was selected. C) The resulting MCA was projected down to 3 dimensions for the purpose of density based clustering (only the first 2 dimensions are shown). Each dot represents a tumor D) HDBSCAN was used to perform hierarchical density based clustering tuned to minimize the fraction of unassigned tumors. E) HDBSCAN clustered tumors were visualized on the UMAP projection; tumors are colored by cluster membership (see legend). This results in some tumors remaining unclassified (grey points). F) kNN was used to assign membership to the remaining tumors.

**Survival**

In order to understand how the diagnosis of cancer changes the expectation for longevity of patients, we first modeled the longevity of the overall population of the U.S. based on CDC life tables. Most TCGA cases are diagnosed late in life, so we fit a model that emphasizes accuracy in the right tail of the distribution. We modeled baseline longevity using a risk rate function:

$$R(t) = R_{20}\exp(a(A-20) + D(t)ka(\frac{t}{365}))$$

Time is split into two components. $A$ going from 0 up to the age of the patient at diagnosis and then remaining fixed, whereas $t$ starts at 0 upon diagnosis and represents the days since diagnosis, a field recorded in the dataset. $D(t)$ is an indicator for whether diagnosis has occurred or not (before diagnosis $D(t) = 0$, after $D(t) = 1$). The model assumes that the risk of death is $R_{20}$ at age 20 years and increases exponentially with a constant rate $a$ up to the age at diagnosis. Risk of death assumes a new rate $ka$ thereafter. Though the model extends to earlier ages, relatively few patients were under age 20.

Given this risk per unit time, the probability of death at time $T + dT$ is the probability to survive to time $T$ which is $(1 - P(T))$ and then die in the remaining interval which is $R(T)dT$

$$dP = (1 - P(T))R(T)dT$$

leading to the differential equation for the cumulative probability of death $P(T)$

$$\frac{dP}{dT} = (1 - P(T))R(T)$$

The solution of this ODE is:

$$P(T) = 1 - \exp(-\int_0^T R\,(t)dt)$$

And the density of deaths per unit time is the derivative with respect to T:

$$p(T) = R(T)\exp(-\int_0^T R\,(t)dt)$$

The $R(T)$ function has two important parameters, the $a$ value which represents the background risk rates, and the $k$ value, which is a function of both cancer tissue type, and the cancer cluster identity.

$$k = k_{tis}k_{cl}$$

The product $k_{tis}k_{cl}$ is obviously symmetric between the two factors. To disambiguate the meaning of the two, the prior distribution of $k_{cl}$ has peak probability at 1 and a relatively narrow width, due to the fact that it is a tissue specific multiplicative modifying factor. This allows the overall magnitude of $k$ to be primarily determined by the $k_{tis}$ value whose prior is significantly less constrained so that there is a wider range of risk across tissue types.

The $k$ in our model can be taken as, roughly, the aging rate relative to the baseline rates. Thus, a value of $k = 2$ means that a given cancer causes you to experience the same risk in time $dt$ as a non-cancer patient would experience at the same age, in time $2dt$.

Using this model, we split the data by Male/Female sex, and by White *vs.* Black race as these were the main well-identified categories available to us in the dataset. Sex is known to be a risk factor for death, with males dying at slightly higher rates for all ages of interest in our dataset. Black *vs.* White race is an important category due to differences in access to care and other socioeconomic and education related factors. However, due to the low number of Black patient tumors, we use ethnicity only to identify the baseline risk function, while the tissue- and cluster-specific multipliers are not ethnicity specific. There were too few tumors from other races to obtain good inference so we excluded them from our analysis.

**Supplementary figure SM2** gives a comparison between the post-fit survival curves for men and women vs the CDC data it is based on. This comparison does not distinguish on ethnicity for simplicity in visualization.

*Figure SM2*

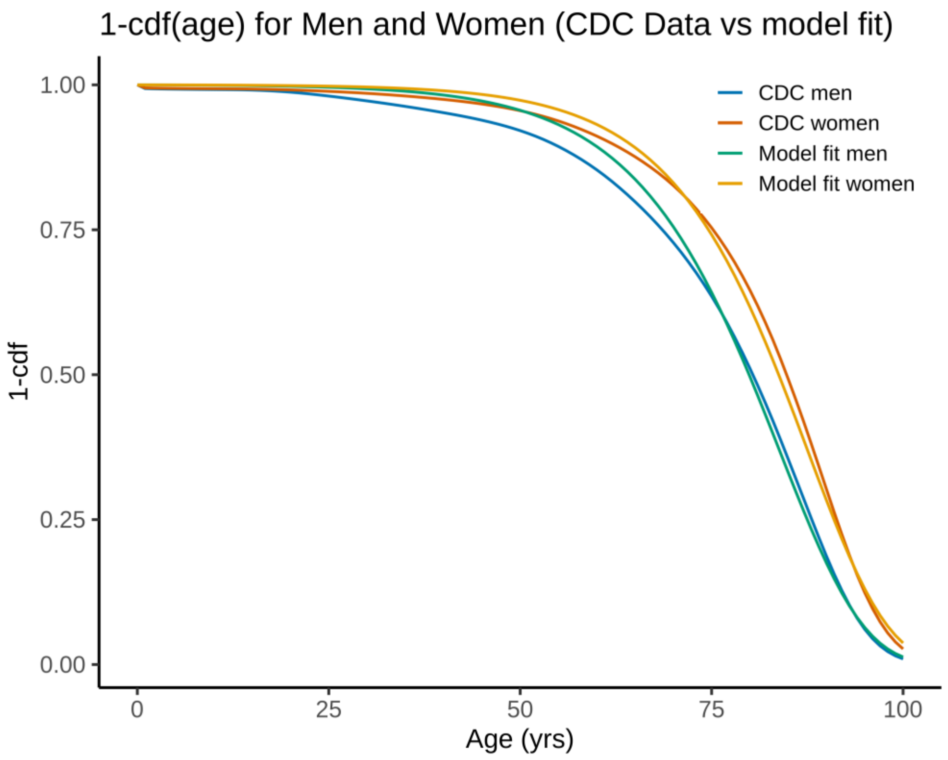


**Supplementary Figure SM2** Comparison between CDC data and ODE model fit for males and females of European descent.

As an aside, we did not explicitly account for stage at diagnosis in our model for the following reasons. In the tumor sample data, stage at diagnosis is confounded with cancer type because some cancers are screened aggressively ( *e.g.* colorectal and prostate cancers) while others are diagnosed typically after they become problematic for the patient's lifestyle ( *e.g.* ovarian and pancreatic cancers). Secondly, such a model specification would suffer from added noise because the staging data are not well standardized across cancer types, have different criteria, and because it is unclear what the relationship between stage and advancement of disease is (for example some sub-stage 4 tumors are metastatic). To compound this latter issue, our tumor set is vastly under-powered given the uneven representation of stage across cancer types.
