## Supplemental Figures for "A molecular taxonomy of tumors independent of tissue-of-origin"

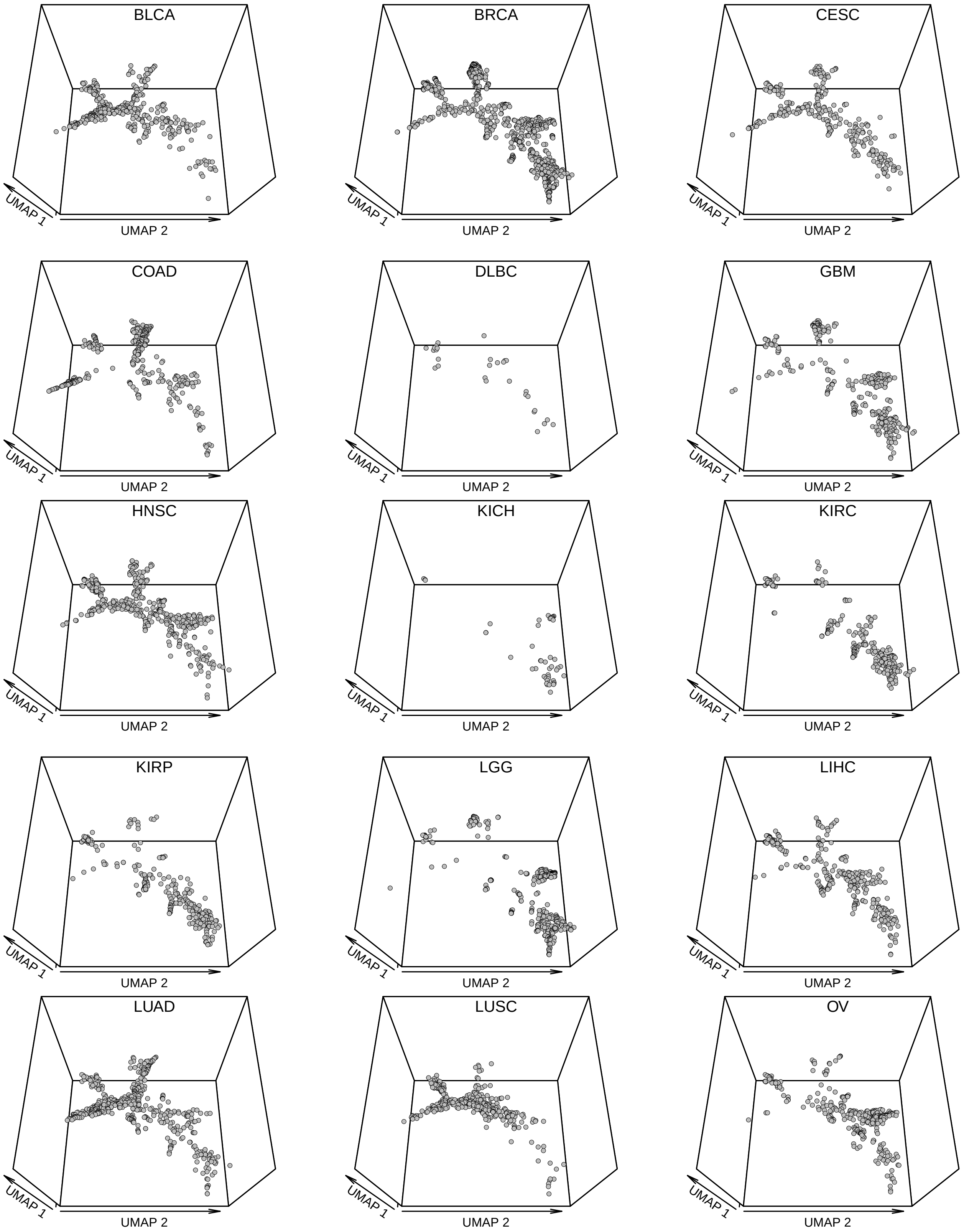
BLCA
BRCA
CESC
COAD
DLBC
GBM
HNSC
KICH
KIRC
KIRP
LGG
LIHC
LUAD
LUSC
OV
UMAP 1
UMAP 2

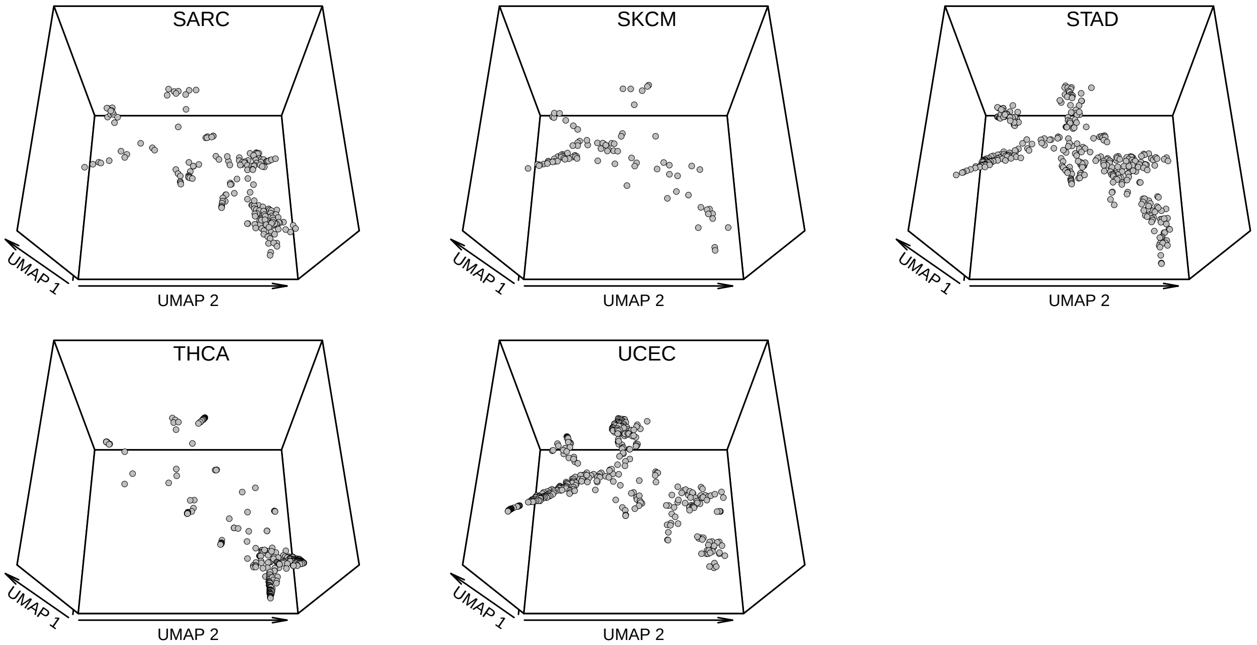


**Supplementary figure S1. Distribution of cancer type on UMAP clusters.** Each panel shows the identity of individual tumors by one of 23 cancer types projected onto the cluster UMAP identified in **Figure 2 of the main text**.

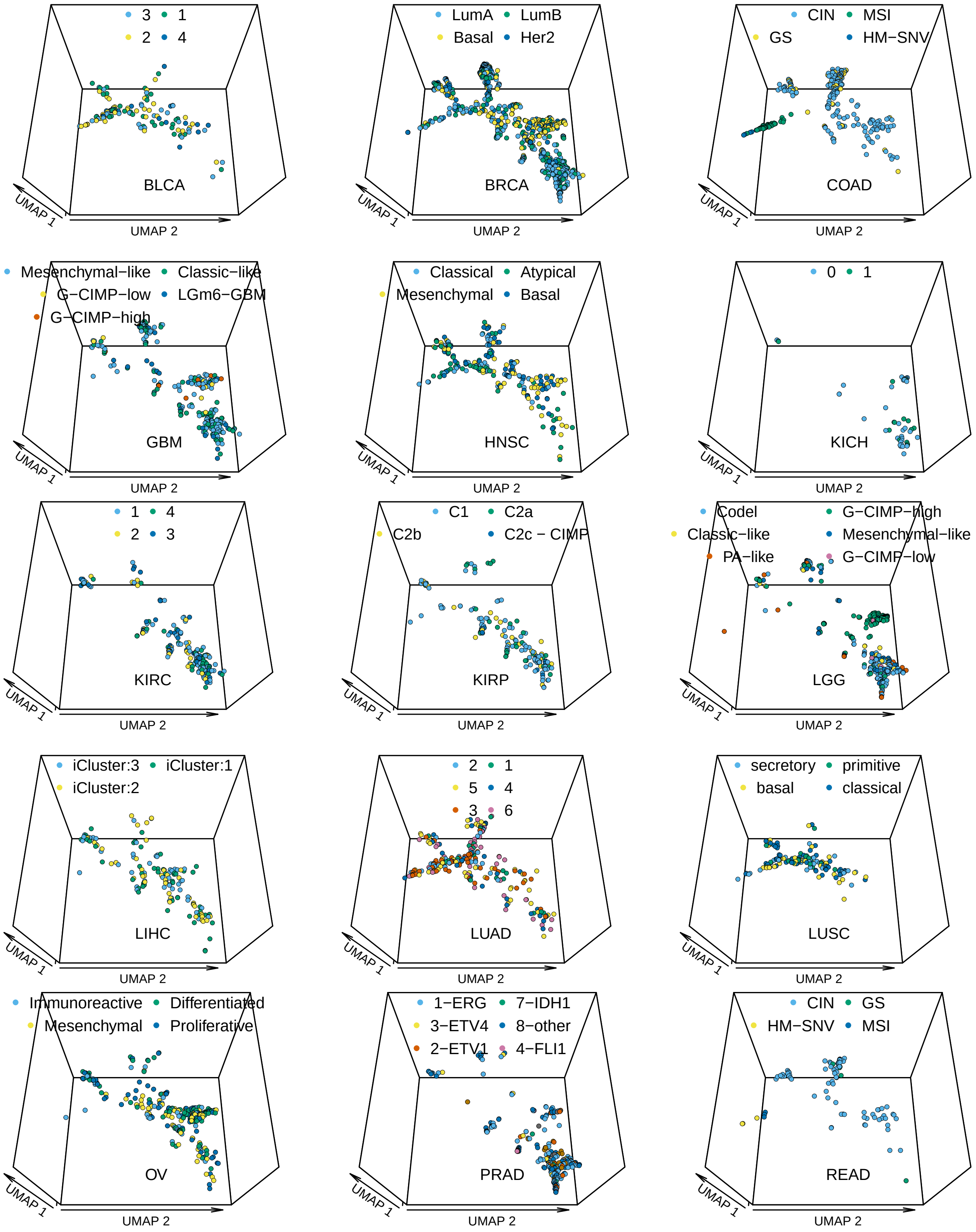
3
1
2
4
BLCA
UMAP 1
UMAP 2
LumA
LumB
Basal
Her2
BRCA
UMAP 1
UMAP 2
CIN
MSI
GS
HM−SNV
COAD
UMAP 1
UMAP 2
Mesenchymal−like
Classic−like
G−CIMP−low
LGm6−GBM
G−CIMP−high
GBM
UMAP 1
UMAP 2
Classical
Atypical
Mesenchymal
Basal
HNSC
UMAP 1
UMAP 2
0
1
KICH
UMAP 1
UMAP 2
1
4
2
3
KIRC
UMAP 1
UMAP 2
C1
C2a
C2b
C2c − CIMP
KIRP
UMAP 1
UMAP 2
Codel
G−CIMP−high
Classic−like
Mesenchymal−like
PA−like
G−CIMP−low
LGG
UMAP 1
UMAP 2
iCluster:3
iCluster:1
iCluster:2
LIHC
UMAP 1
UMAP 2
2
1
5
4
3
6
LUAD
UMAP 1
UMAP 2
secretory
primitive
basal
classical
LUSC
UMAP 1
UMAP 2
Immunoreactive
Differentiated
Mesenchymal
Proliferative
OV
UMAP 1
UMAP 2
1−ERG
7−IDH1
3−ETV4
8−other
2−ETV1
4−FLI1
PRAD
UMAP 1
UMAP 2
CIN
GS
HM−SNV
MSI
READ
UMAP 1
UMAP 2

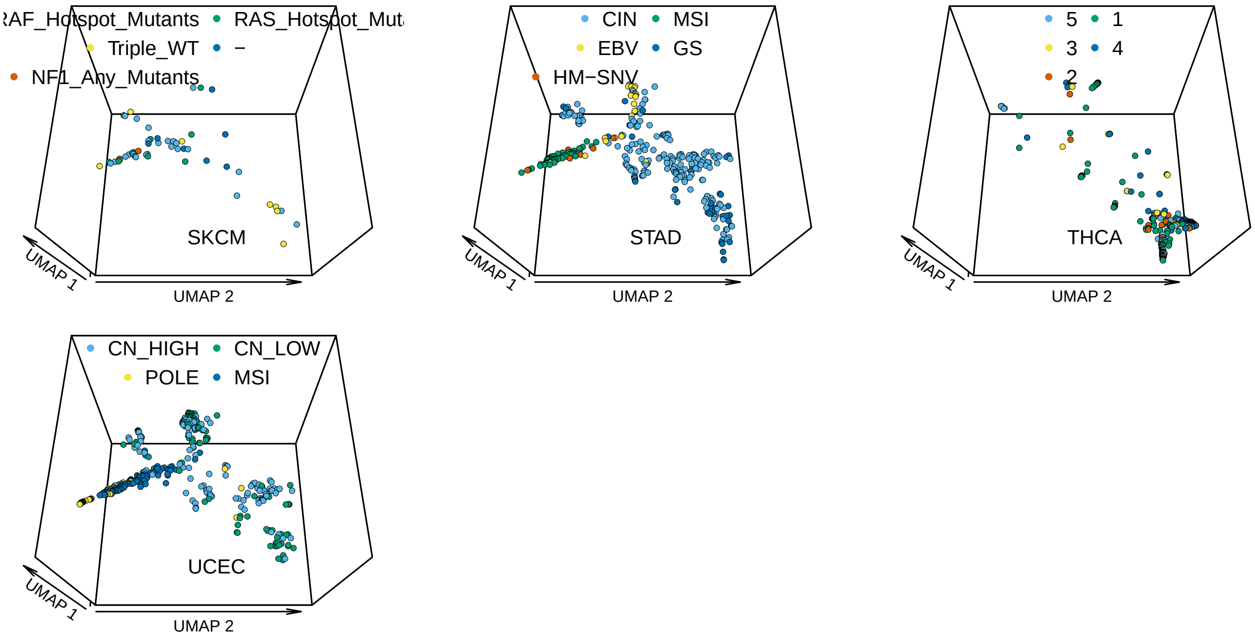


**Supplementary Figure S2. Histological subtypes.** Each panel shows the histological subtypes of one of the 23 cancers projected onto the tumors of the UMAP from **Figure 2 of the main text**.

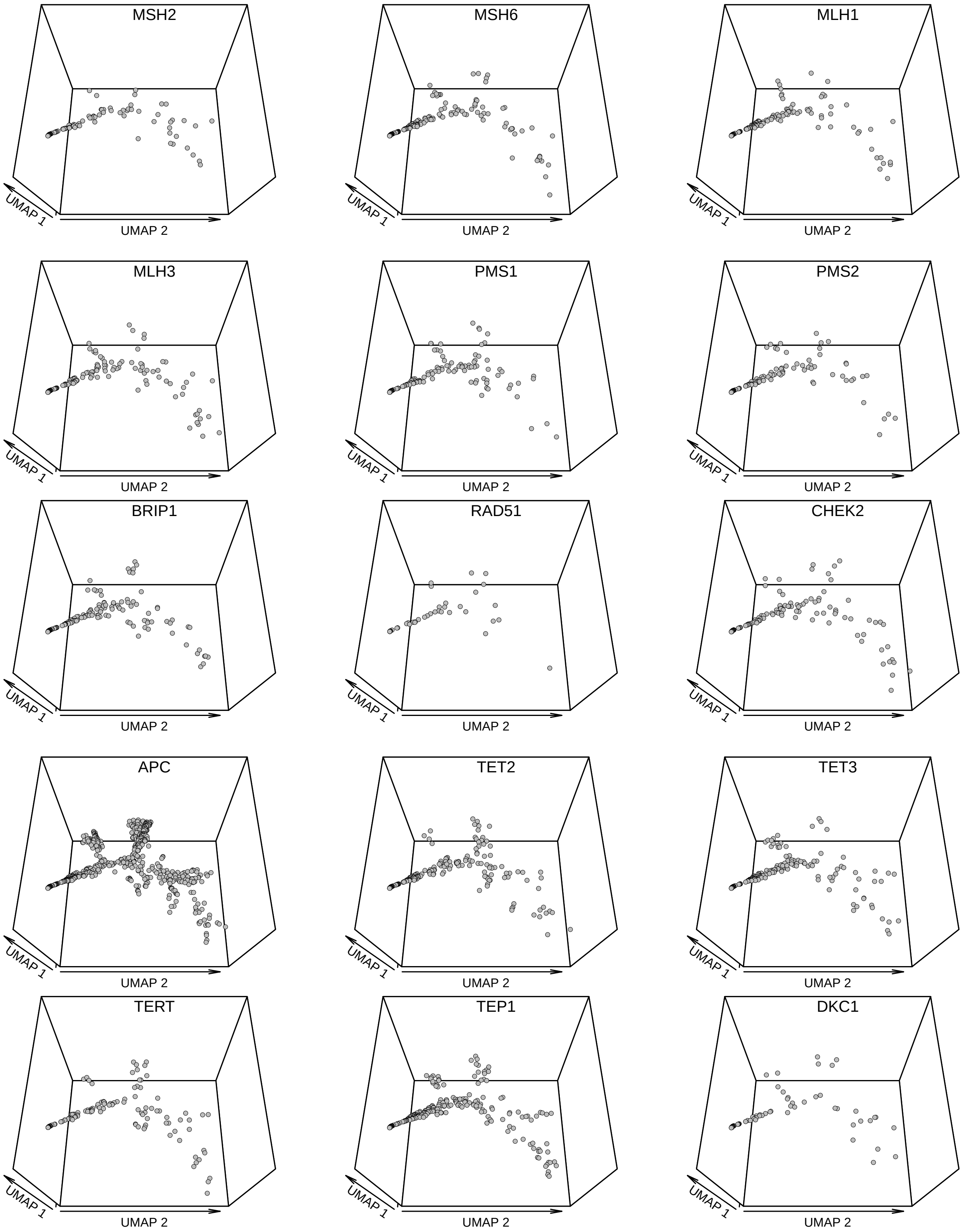


**Supplementary Figure S3. Distribution of mismatch repair genes on UMAP clusters.** Each panel shows one of 15 genes involved in genome stability projected on the UMAP from **Figure 2 of the main text**.

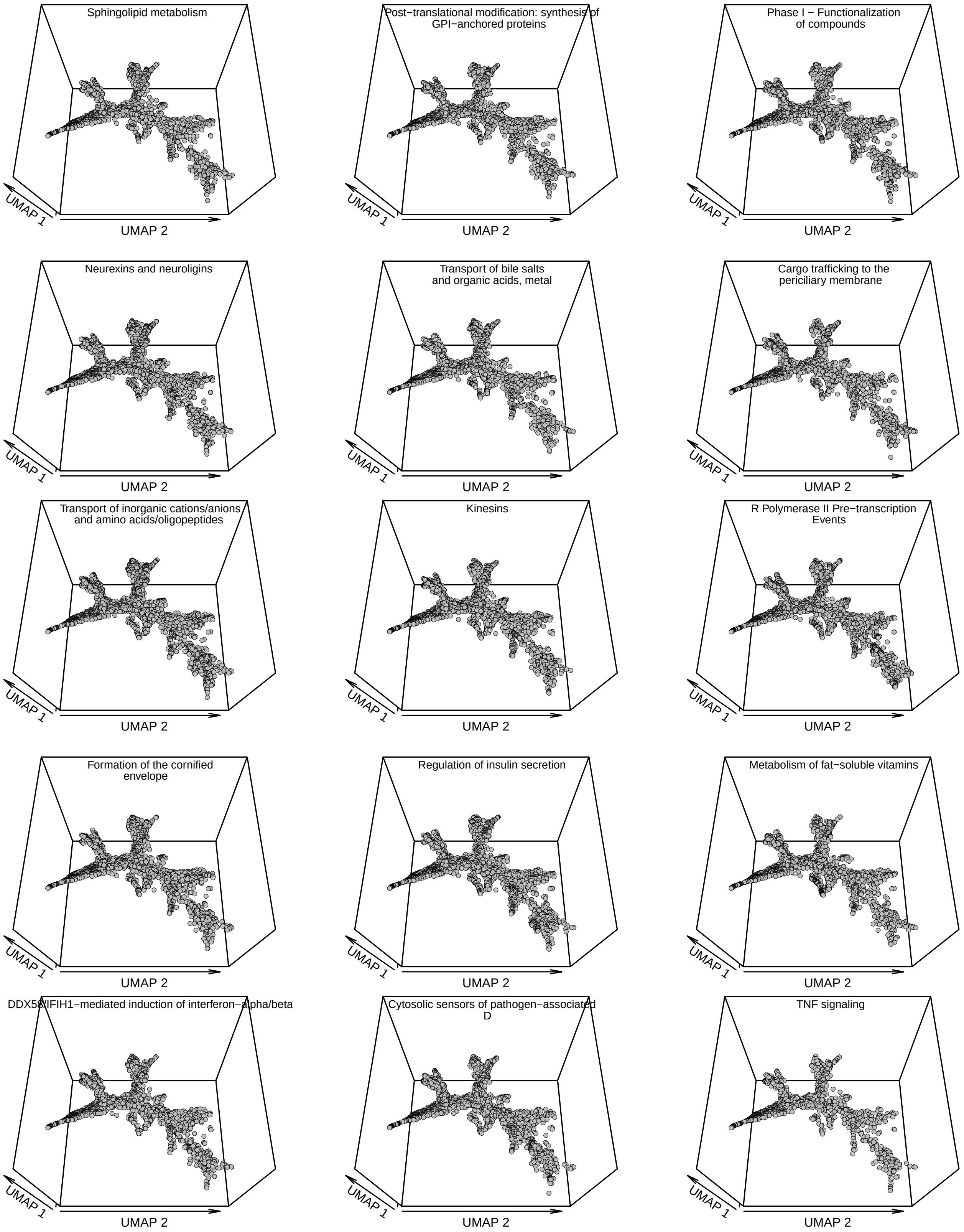
Sphingolipid metabolism
UMAP 1
UMAP 2
Post−translational modification: synthesis of GPI−anchored proteins
UMAP 1
UMAP 2
Phase I − Functionalization of compounds
UMAP 1
UMAP 2
Neurexins and neuroligins
UMAP 1
UMAP 2
Transport of bile salts and organic acids, metal
UMAP 1
UMAP 2
Cargo trafficking to the periciliary membrane
UMAP 1
UMAP 2
Transport of inorganic cations/anions and amino acids/oligopeptides
UMAP 1
UMAP 2
Kinesins
UMAP 1
UMAP 2
R Polymerase II Pre−transcription Events
UMAP 1
UMAP 2
Formation of the cornified envelope
UMAP 1
UMAP 2
Regulation of insulin secretion
UMAP 1
UMAP 2
Metabolism of fat−soluble vitamins
UMAP 1
UMAP 2
DDX58/IFIH1−mediated induction of interferon−alpha/beta
UMAP 1
UMAP 2
Cytosolic sensors of pathogen−associated D
UMAP 1
UMAP 2
TNF signaling
UMAP 1
UMAP 2

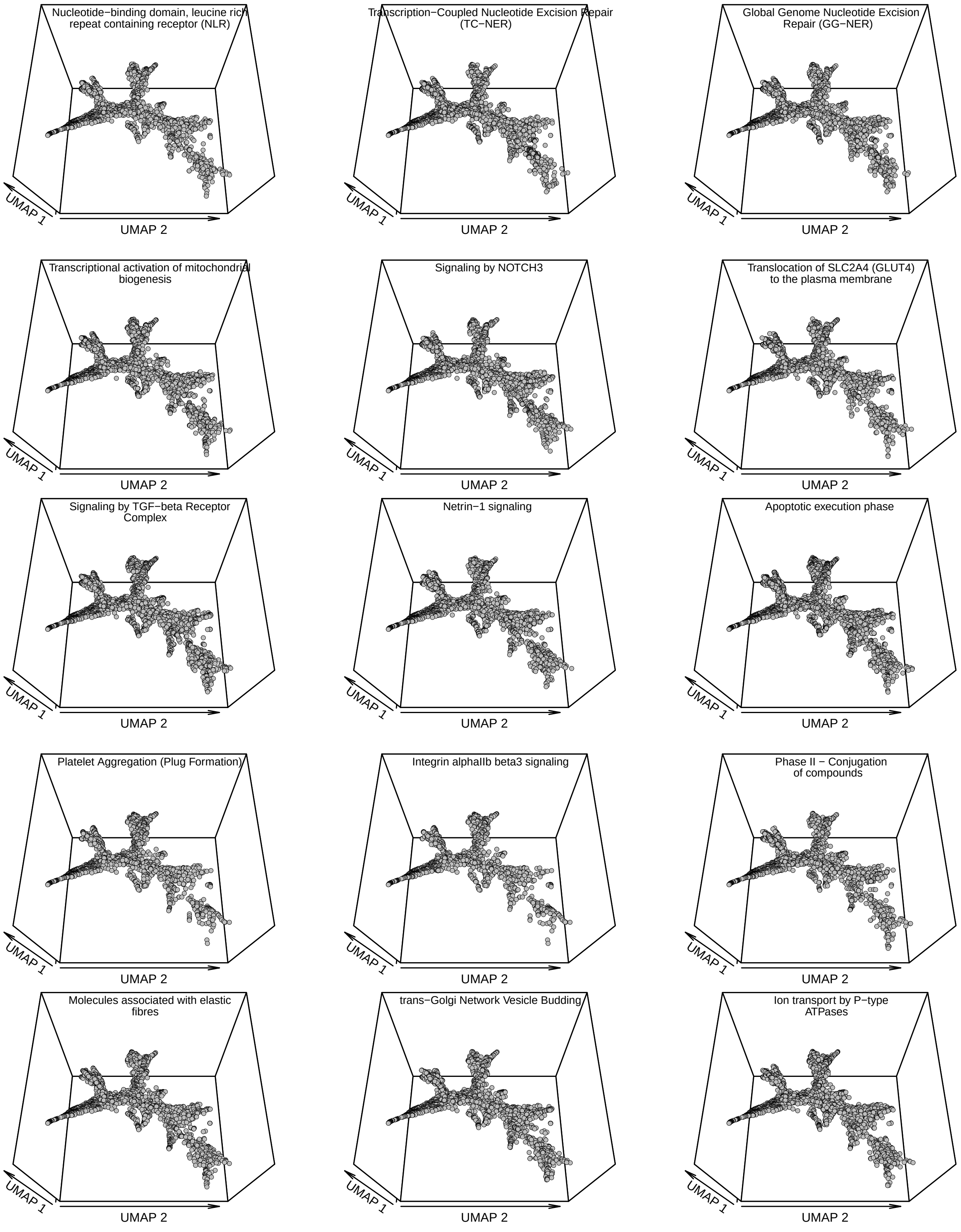
Nucleotide−binding domain, leucine rich repeat containing receptor (NLR)
UMAP 1
UMAP 2
Transcription−Coupled Nucleotide Excision Repair (TC−NER)
UMAP 1
UMAP 2
Global Genome Nucleotide Excision Repair (GG−NER)
UMAP 1
UMAP 2
Transcriptional activation of mitochondrial biogenesis
UMAP 1
UMAP 2
Signaling by NOTCH3
UMAP 1
UMAP 2
Translocation of SLC2A4 (GLUT4) to the plasma membrane
UMAP 1
UMAP 2
Signaling by TGF−beta Receptor Complex
UMAP 1
UMAP 2
Netrin−1 signaling
UMAP 1
UMAP 2
Apoptotic execution phase
UMAP 1
UMAP 2
Platelet Aggregation (Plug Formation)
UMAP 1
UMAP 2
Integrin alphaIIb beta3 signaling
UMAP 1
UMAP 2
Phase II − Conjugation of compounds
UMAP 1
UMAP 2
Molecules associated with elastic fibres
UMAP 1
UMAP 2
trans−Golgi Network Vesicle Budding
UMAP 1
UMAP 2
Ion transport by P−type ATPases
UMAP 1
UMAP 2

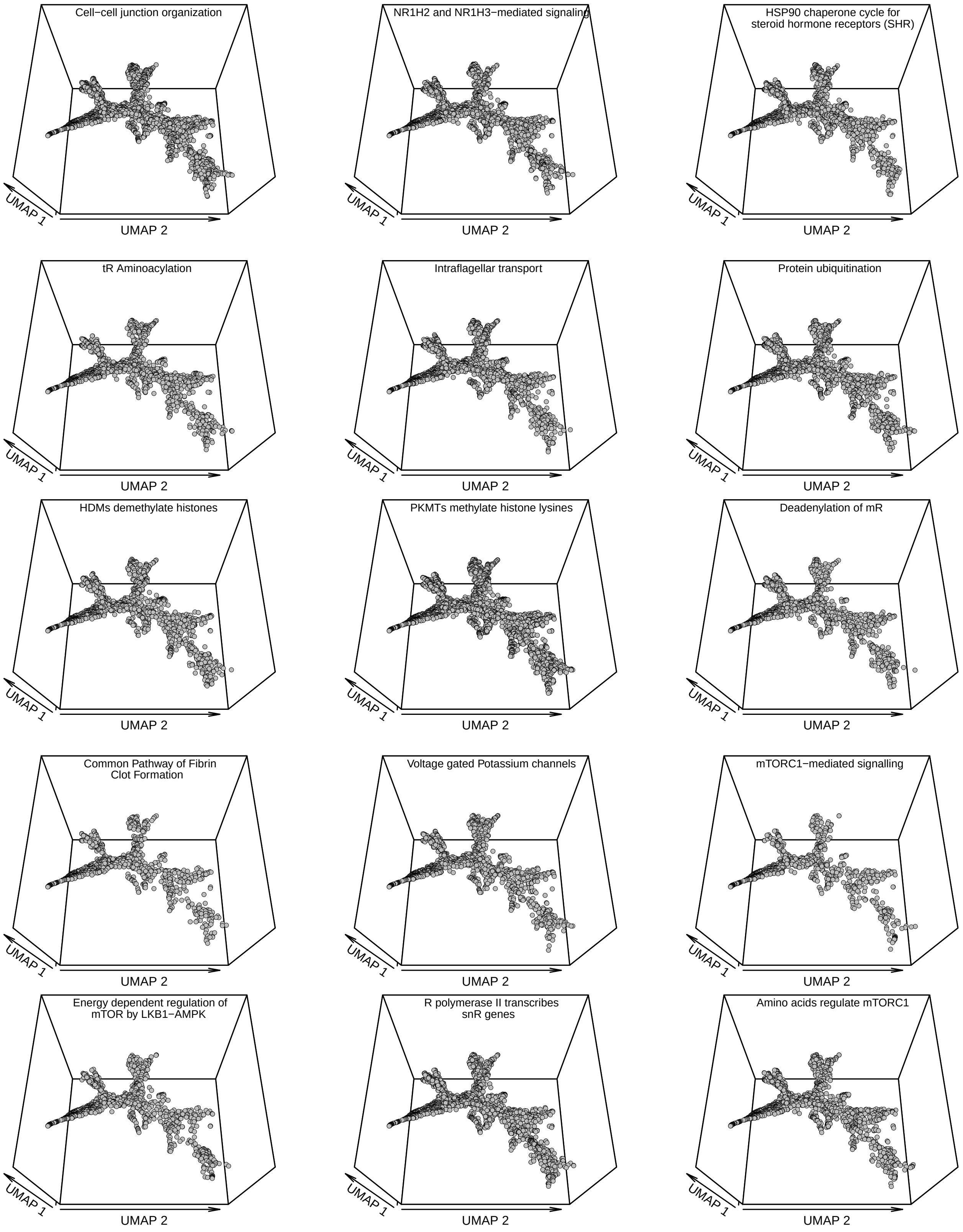
Cell−cell junction organization
UMAP 1
UMAP 2
NR1H2 and NR1H3−mediated signaling
UMAP 1
UMAP 2
HSP90 chaperone cycle for
steroid hormone receptors (SHR)
UMAP 1
UMAP 2
tR Aminoacylation
UMAP 1
UMAP 2
Intraflagellar transport
UMAP 1
UMAP 2
Protein ubiquitination
UMAP 1
UMAP 2
HDMs demethylate histones
UMAP 1
UMAP 2
PKMTs methylate histone lysines
UMAP 1
UMAP 2
Deadenylation of mR
UMAP 1
UMAP 2
Common Pathway of Fibrin
Clot Formation
UMAP 1
UMAP 2
Voltage gated Potassium channels
UMAP 1
UMAP 2
mTORC1−mediated signalling
UMAP 1
UMAP 2
Energy dependent regulation of
mTOR by LKB1−AMPK
UMAP 1
UMAP 2
R polymerase II transcribes
snR genes
UMAP 1
UMAP 2
Amino acids regulate mTORC1
UMAP 1
UMAP 2

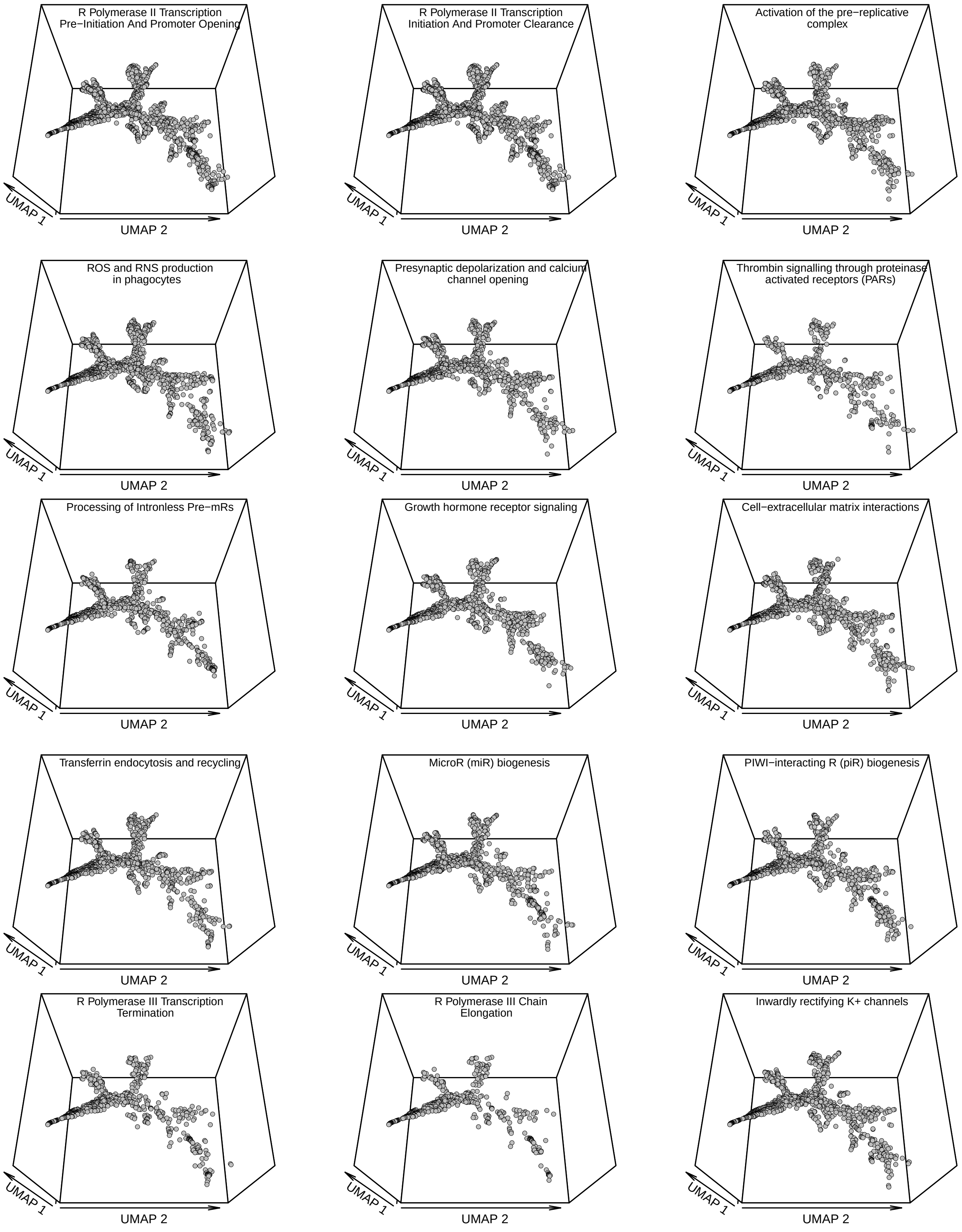
R Polymerase II Transcription Pre−Initiation And Promoter Opening
UMAP 1
UMAP 2
R Polymerase II Transcription Initiation And Promoter Clearance
UMAP 1
UMAP 2
Activation of the pre−replicative complex
UMAP 1
UMAP 2
ROS and RNS production in phagocytes
UMAP 1
UMAP 2
Presynaptic depolarization and calcium channel opening
UMAP 1
UMAP 2
Thrombin signalling through proteinase activated receptors (PARs)
UMAP 1
UMAP 2
Processing of Intronless Pre−mRs
UMAP 1
UMAP 2
Growth hormone receptor signaling
UMAP 1
UMAP 2
Cell−extracellular matrix interactions
UMAP 1
UMAP 2
Transferrin endocytosis and recycling
UMAP 1
UMAP 2
MicroR (miR) biogenesis
UMAP 1
UMAP 2
PIWI−interacting R (piR) biogenesis
UMAP 1
UMAP 2
R Polymerase III Transcription Termination
UMAP 1
UMAP 2
R Polymerase III Chain Elongation
UMAP 1
UMAP 2
Inwardly rectifying K+ channels
UMAP 1
UMAP 2

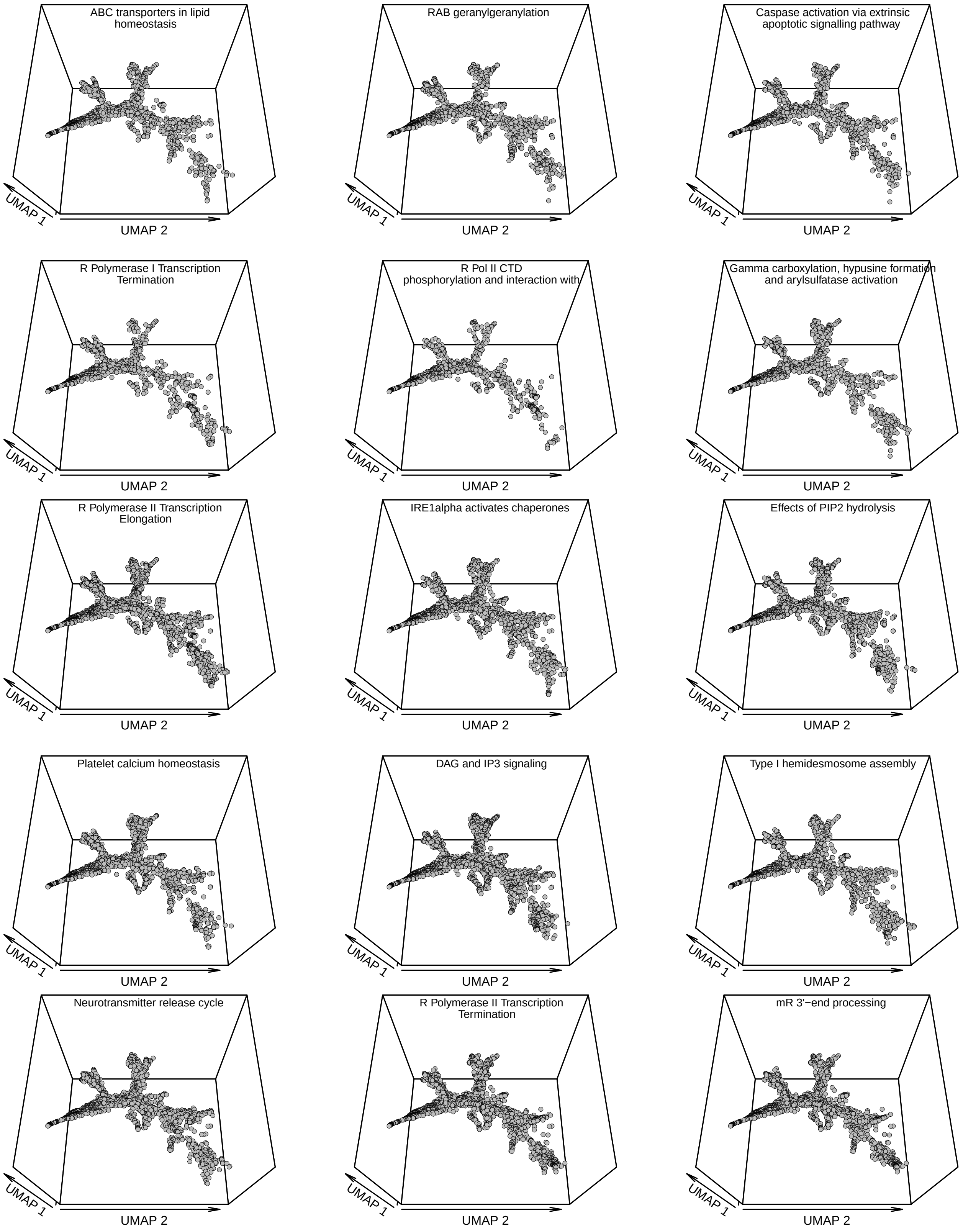
ABC transporters in lipid homeostasis
RAB geranylgeranylation
Caspase activation via extrinsic apoptotic signalling pathway
R Polymerase I Transcription Termination
R Pol II CTD phosphorylation and interaction with
Gamma carboxylation, hypusine formation and arylsulfatase activation
R Polymerase II Transcription Elongation
IRE1alpha activates chaperones
Effects of PIP2 hydrolysis
Platelet calcium homeostasis
DAG and IP3 signaling
Type I hemidesmosome assembly
Neurotransmitter release cycle
R Polymerase II Transcription Termination
mR 3'−end processing
UMAP 1
UMAP 2

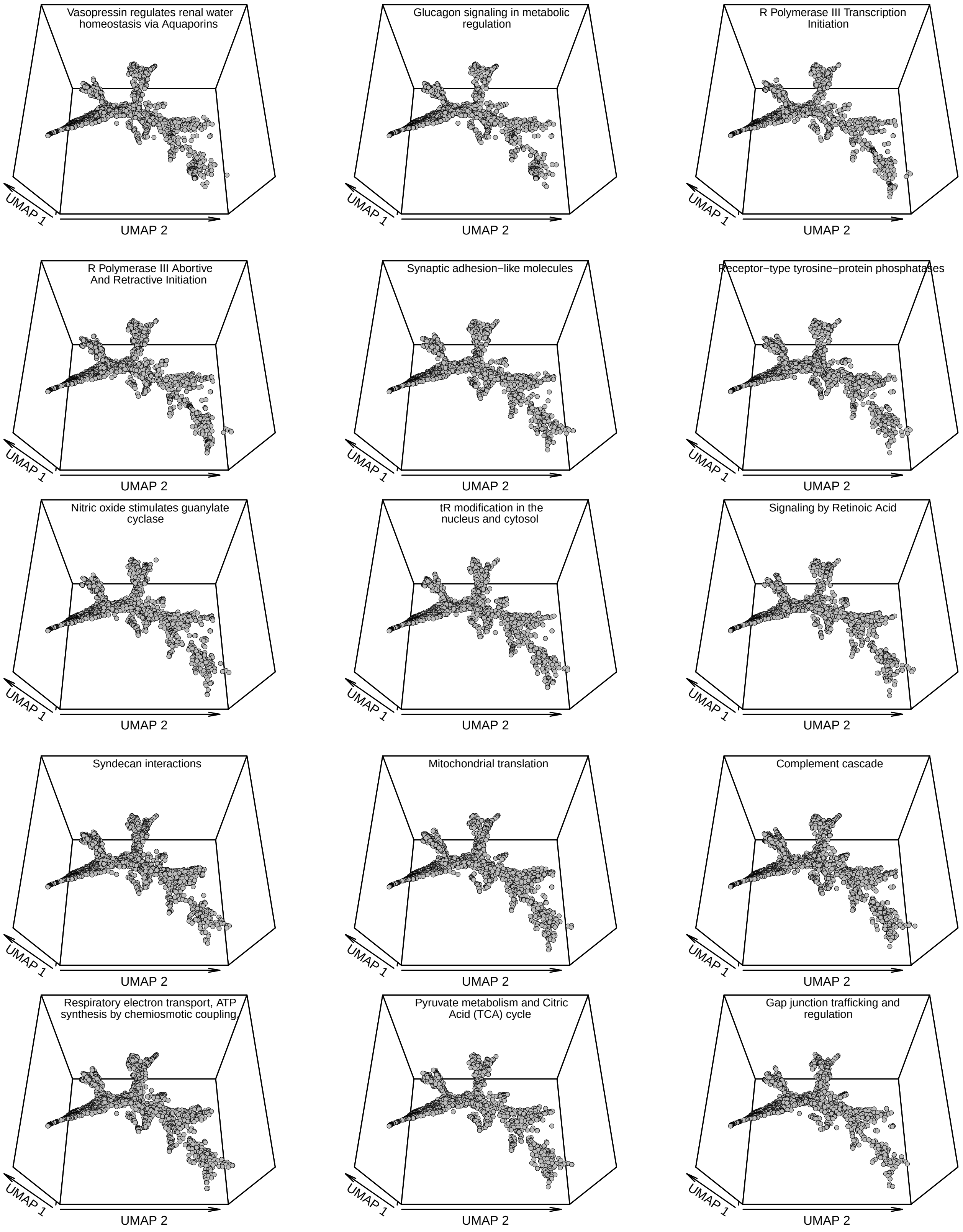
Vasopressin regulates renal water homeostasis via Aquaporins
Glucagon signaling in metabolic regulation
R Polymerase III Transcription Initiation
R Polymerase III Abortive And Retractive Initiation
Synaptic adhesion−like molecules
Receptor−type tyrosine−protein phosphatases
Nitric oxide stimulates guanylate cyclase
tR modification in the nucleus and cytosol
Signaling by Retinoic Acid
Syndecan interactions
Mitochondrial translation
Complement cascade
Respiratory electron transport, ATP synthesis by chemiosmotic coupling
Pyruvate metabolism and Citric Acid (TCA) cycle
Gap junction trafficking and regulation
UMAP 1
UMAP 2

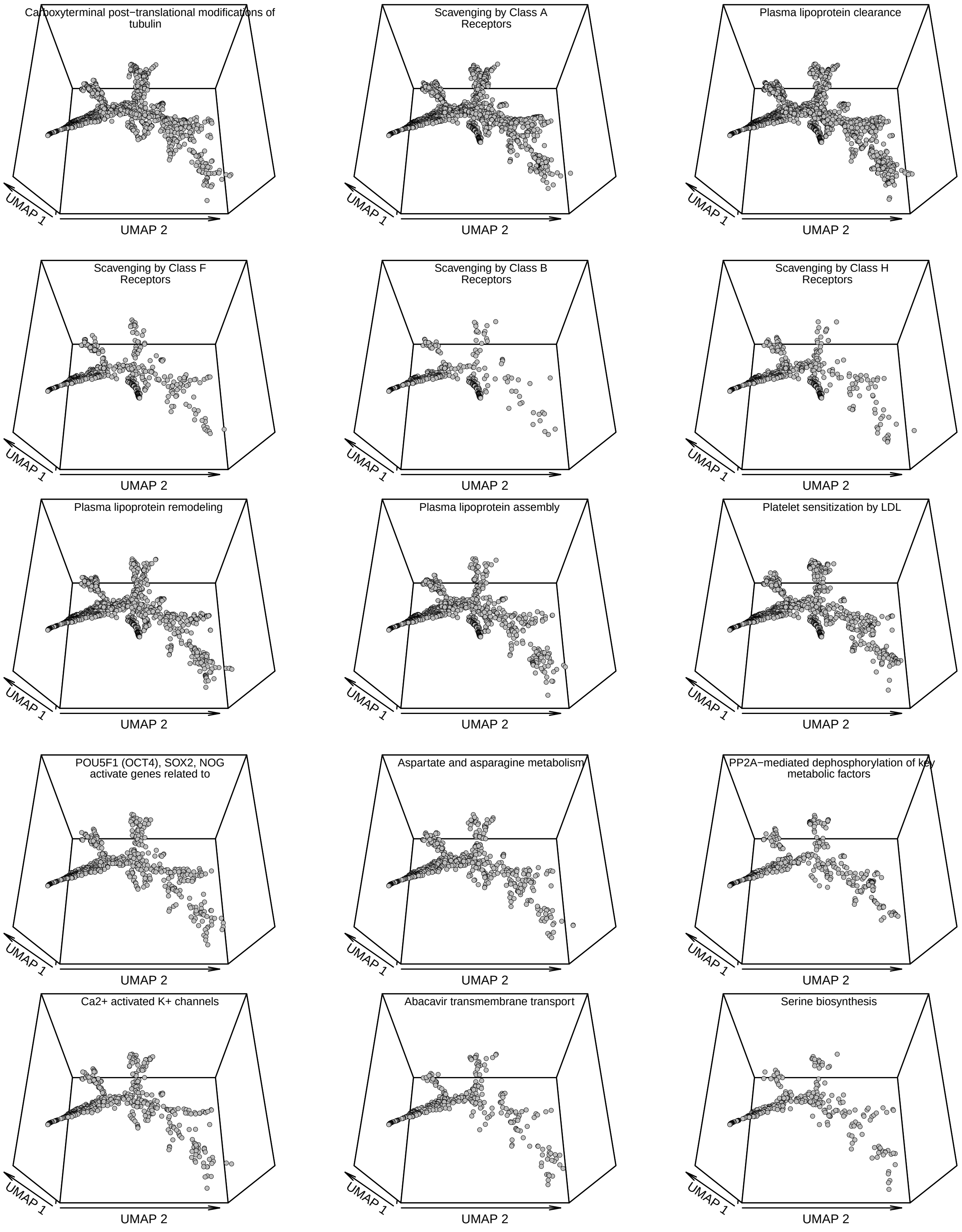
Carboxyterminal post−translational modifications of tubulin
UMAP 1
UMAP 2
Scavenging by Class A Receptors
UMAP 1
UMAP 2
Plasma lipoprotein clearance
UMAP 1
UMAP 2
Scavenging by Class F Receptors
UMAP 1
UMAP 2
Scavenging by Class B Receptors
UMAP 1
UMAP 2
Scavenging by Class H Receptors
UMAP 1
UMAP 2
Plasma lipoprotein remodeling
UMAP 1
UMAP 2
Plasma lipoprotein assembly
UMAP 1
UMAP 2
Platelet sensitization by LDL
UMAP 1
UMAP 2
POU5F1 (OCT4), SOX2, NOG activate genes related to
UMAP 1
UMAP 2
Aspartate and asparagine metabolism
UMAP 1
UMAP 2
PP2A−mediated dephosphorylation of key metabolic factors
UMAP 1
UMAP 2
Ca2+ activated K+ channels
UMAP 1
UMAP 2
Abacavir transmembrane transport
UMAP 1
UMAP 2
Serine biosynthesis
UMAP 1
UMAP 2

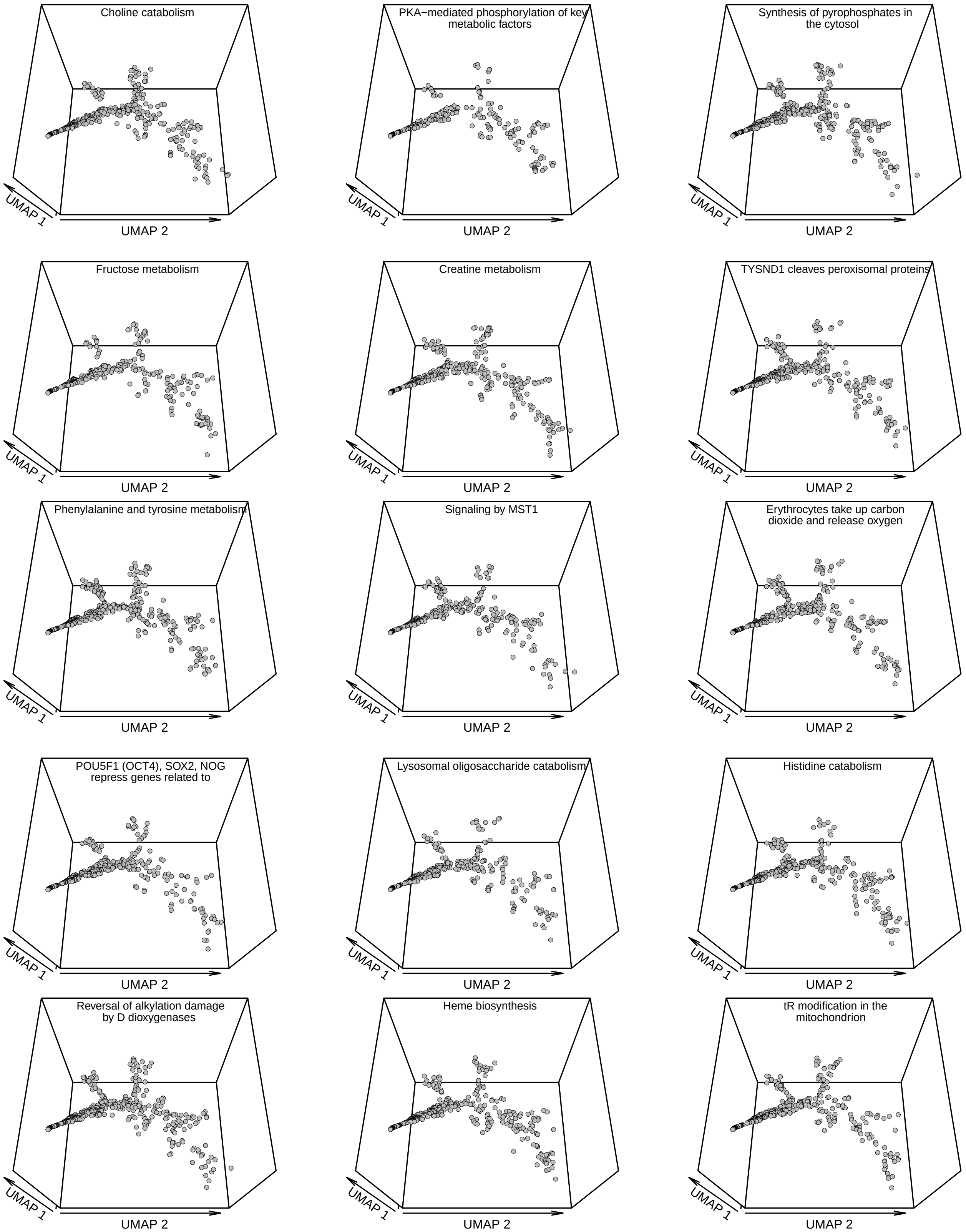
Choline catabolism
UMAP 1
UMAP 2
PKA−mediated phosphorylation of key metabolic factors
UMAP 1
UMAP 2
Synthesis of pyrophosphates in the cytosol
UMAP 1
UMAP 2
Fructose metabolism
UMAP 1
UMAP 2
Creatine metabolism
UMAP 1
UMAP 2
TYSND1 cleaves peroxisomal proteins
UMAP 1
UMAP 2
Phenylalanine and tyrosine metabolism
UMAP 1
UMAP 2
Signaling by MST1
UMAP 1
UMAP 2
Erythrocytes take up carbon dioxide and release oxygen
UMAP 1
UMAP 2
POU5F1 (OCT4), SOX2, NOG repress genes related to
UMAP 1
UMAP 2
Lysosomal oligosaccharide catabolism
UMAP 1
UMAP 2
Histidine catabolism
UMAP 1
UMAP 2
Reversal of alkylation damage by D dioxygenases
UMAP 1
UMAP 2
Heme biosynthesis
UMAP 1
UMAP 2
tR modification in the mitochondrion
UMAP 1
UMAP 2

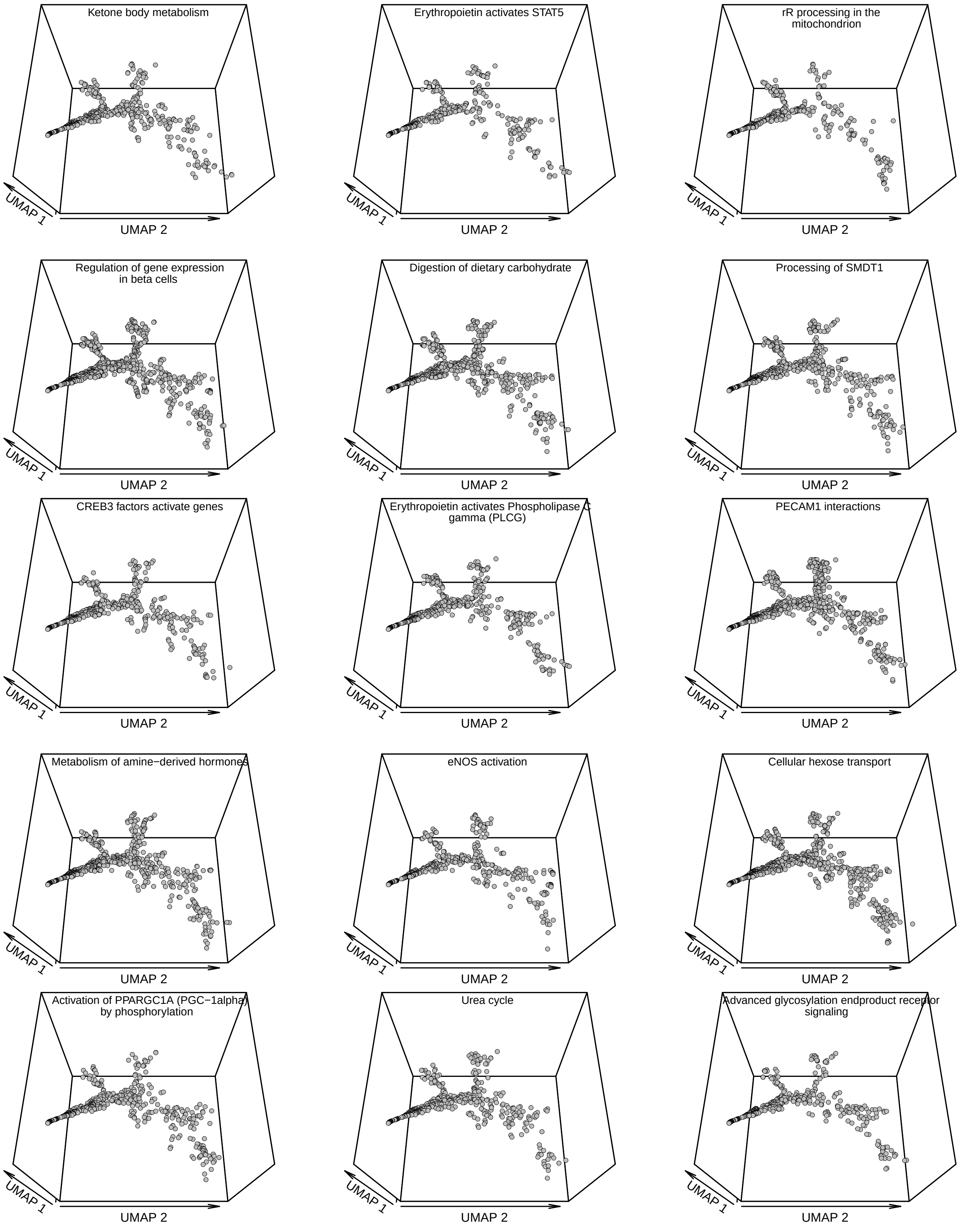

Ketone body metabolism
UMAP 1
UMAP 2
Erythropoietin activates STAT5
UMAP 1
UMAP 2
rR processing in the mitochondrion
UMAP 1
UMAP 2
Regulation of gene expression in beta cells
UMAP 1
UMAP 2
Digestion of dietary carbohydrate
UMAP 1
UMAP 2
Processing of SMDT1
UMAP 1
UMAP 2
CREB3 factors activate genes
UMAP 1
UMAP 2
Erythropoietin activates Phospholipase C gamma (PLCG)
UMAP 1
UMAP 2
PECAM1 interactions
UMAP 1
UMAP 2
Metabolism of amine−derived hormones
UMAP 1
UMAP 2
eNOS activation
UMAP 1
UMAP 2
Cellular hexose transport
UMAP 1
UMAP 2
Activation of PPARGC1A (PGC−1alpha) by phosphorylation
UMAP 1
UMAP 2
Urea cycle
UMAP 1
UMAP 2
Advanced glycosylation endproduct receptor signaling
UMAP 1
UMAP 2

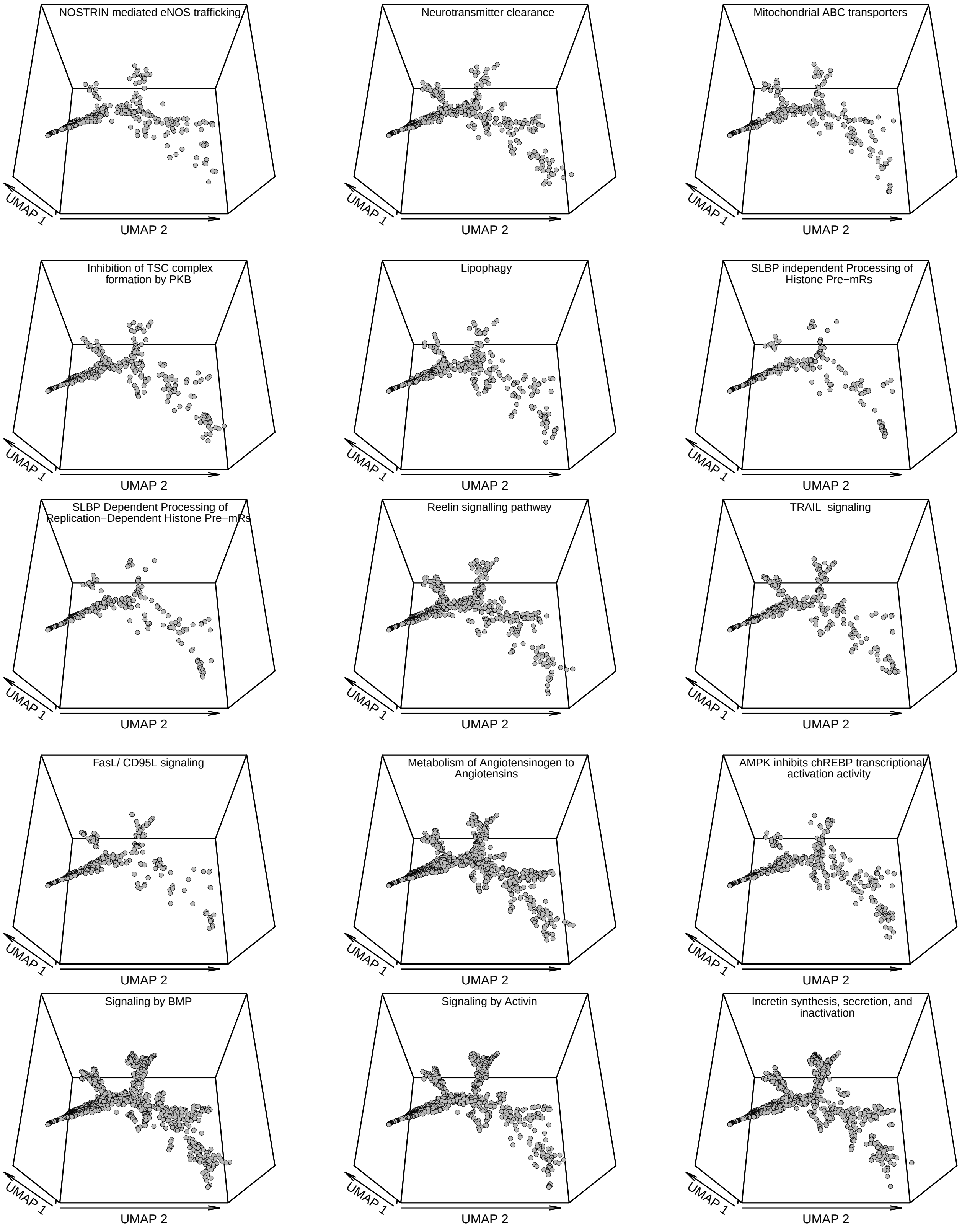
NOSTRIN mediated eNOS trafficking
Neurotransmitter clearance
Mitochondrial ABC transporters
Inhibition of TSC complex formation by PKB
Lipophagy
SLBP independent Processing of Histone Pre−mRs
SLBP Dependent Processing of Replication−Dependent Histone Pre−mRs
Reelin signalling pathway
TRAIL signaling
FasL/ CD95L signaling
Metabolism of Angiotensinogen to Angiotensins
AMPK inhibits chREBP transcriptional activation activity
Signaling by BMP
Signaling by Activin
Incretin synthesis, secretion, and inactivation
UMAP 1
UMAP 2

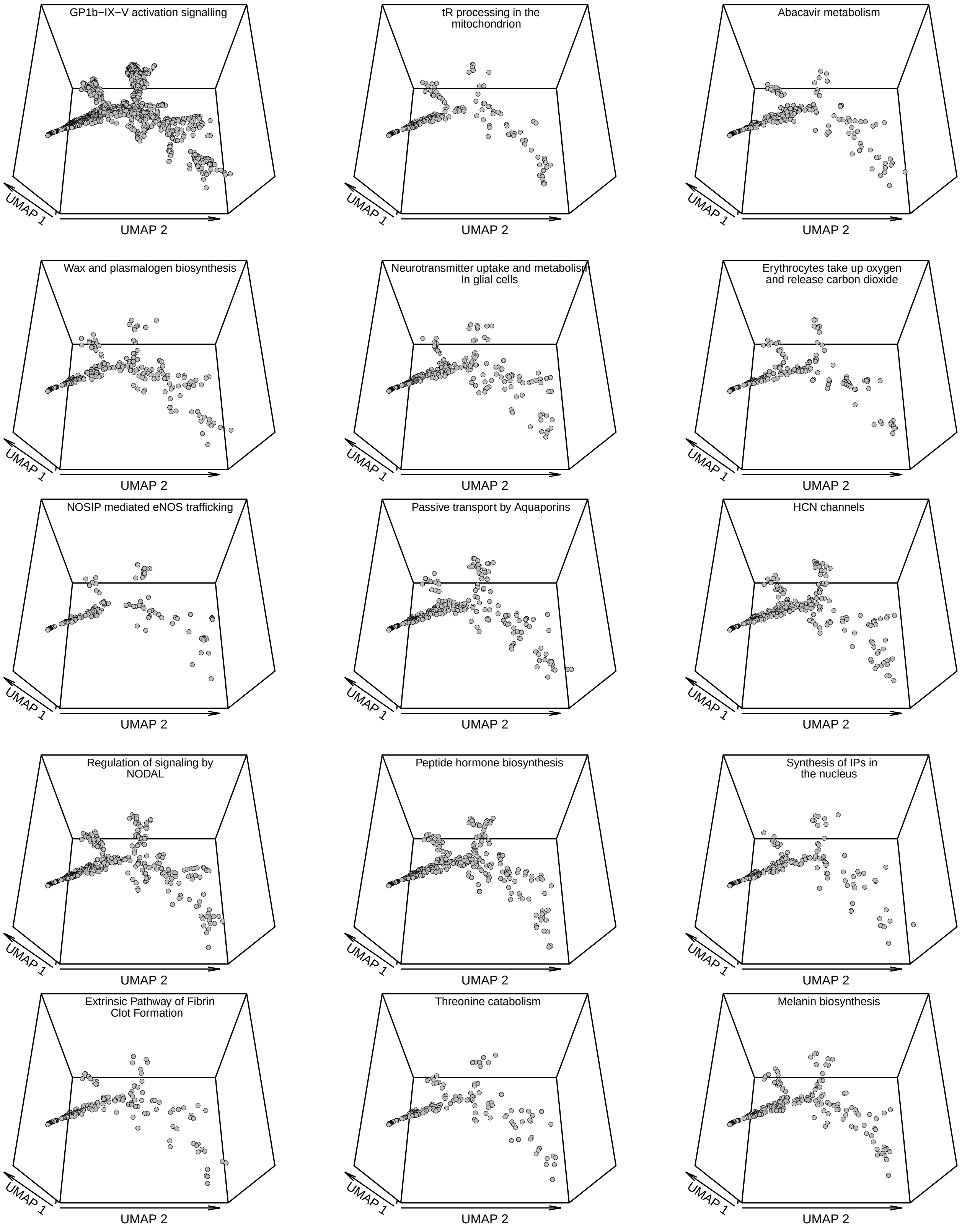
GP1b−IX−V activation signalling
UMAP 1
UMAP 2
tR processing in the mitochondrion
UMAP 1
UMAP 2
Abacavir metabolism
UMAP 1
UMAP 2
Wax and plasmalogen biosynthesis
UMAP 1
UMAP 2
Neurotransmitter uptake and metabolism In glial cells
UMAP 1
UMAP 2
Erythrocytes take up oxygen and release carbon dioxide
UMAP 1
UMAP 2
NOSIP mediated eNOS trafficking
UMAP 1
UMAP 2
Passive transport by Aquaporins
UMAP 1
UMAP 2
HCN channels
UMAP 1
UMAP 2
Regulation of signaling by NODAL
UMAP 1
UMAP 2
Peptide hormone biosynthesis
UMAP 1
UMAP 2
Synthesis of IPs in the nucleus
UMAP 1
UMAP 2
Extrinsic Pathway of Fibrin Clot Formation
UMAP 1
UMAP 2
Threonine catabolism
UMAP 1
UMAP 2
Melanin biosynthesis
UMAP 1
UMAP 2

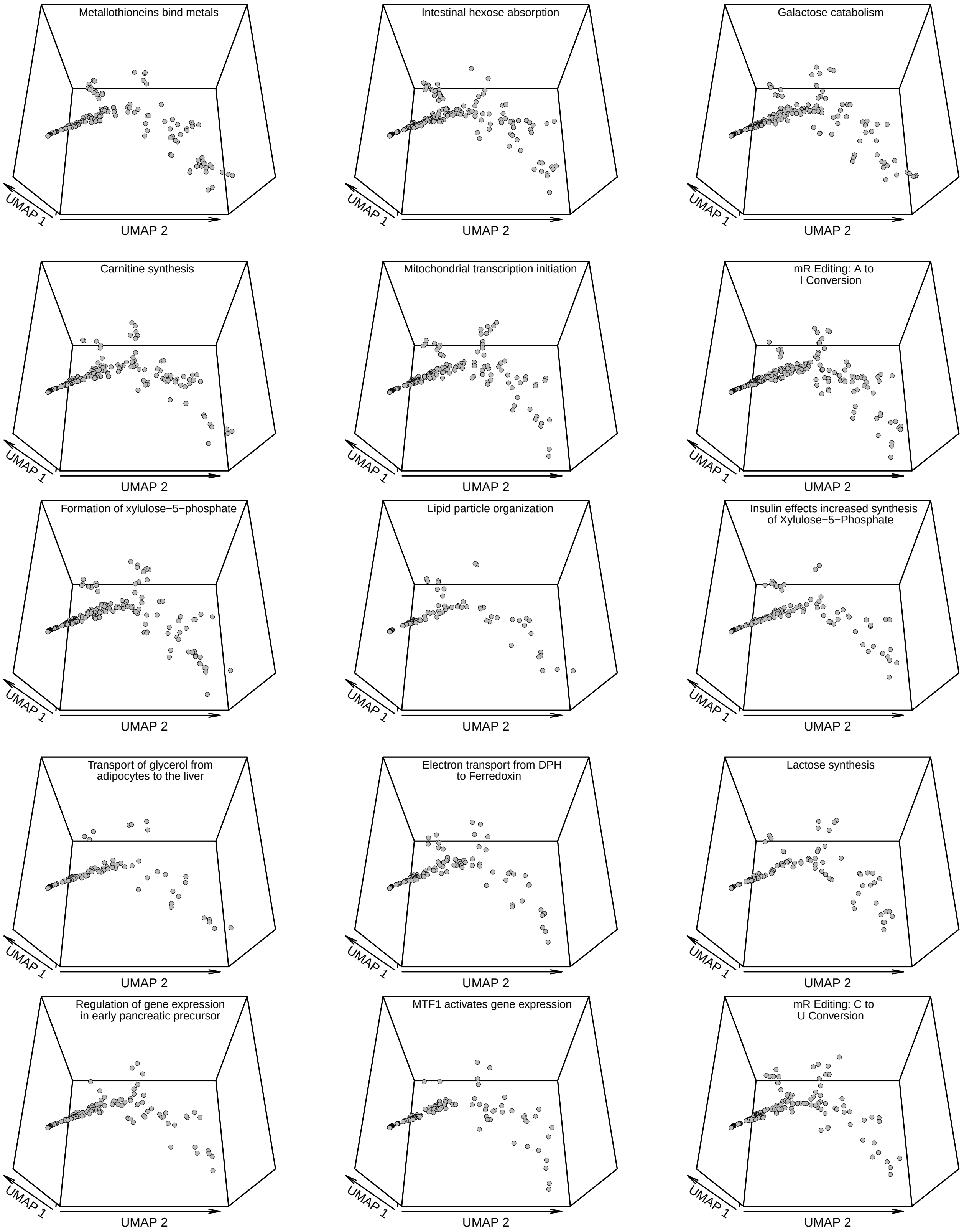
Metallothioneins bind metals
Intestinal hexose absorption
Galactose catabolism
Carnitine synthesis
Mitochondrial transcription initiation
mR Editing: A to I Conversion
Formation of xylulose−5−phosphate
Lipid particle organization
Insulin effects increased synthesis of Xylulose−5−Phosphate
Transport of glycerol from adipocytes to the liver
Electron transport from DPH to Ferredoxin
Lactose synthesis
Regulation of gene expression in early pancreatic precursor
MTF1 activates gene expression
mR Editing: C to U Conversion
UMAP 1
UMAP 2

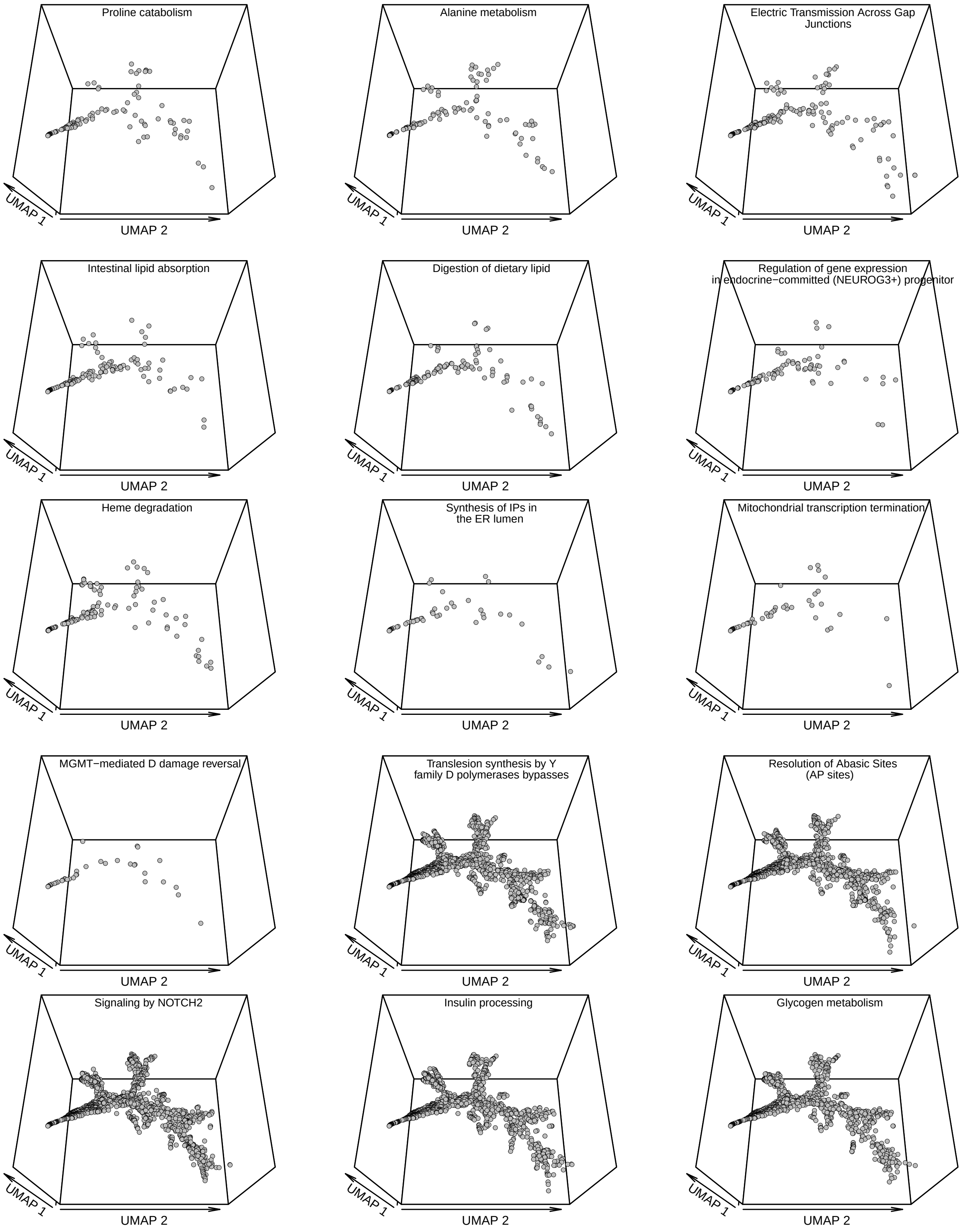
Proline catabolism
Alanine metabolism
Electric Transmission Across Gap Junctions
Intestinal lipid absorption
Digestion of dietary lipid
Regulation of gene expression in endocrine−committed (NEUROG3+) progenitor
Heme degradation
Synthesis of IPs in the ER lumen
Mitochondrial transcription termination
MGMT−mediated D damage reversal
Translesion synthesis by Y family D polymerases bypasses
Resolution of Abasic Sites (AP sites)
Signaling by NOTCH2
Insulin processing
Glycogen metabolism
UMAP 1
UMAP 2

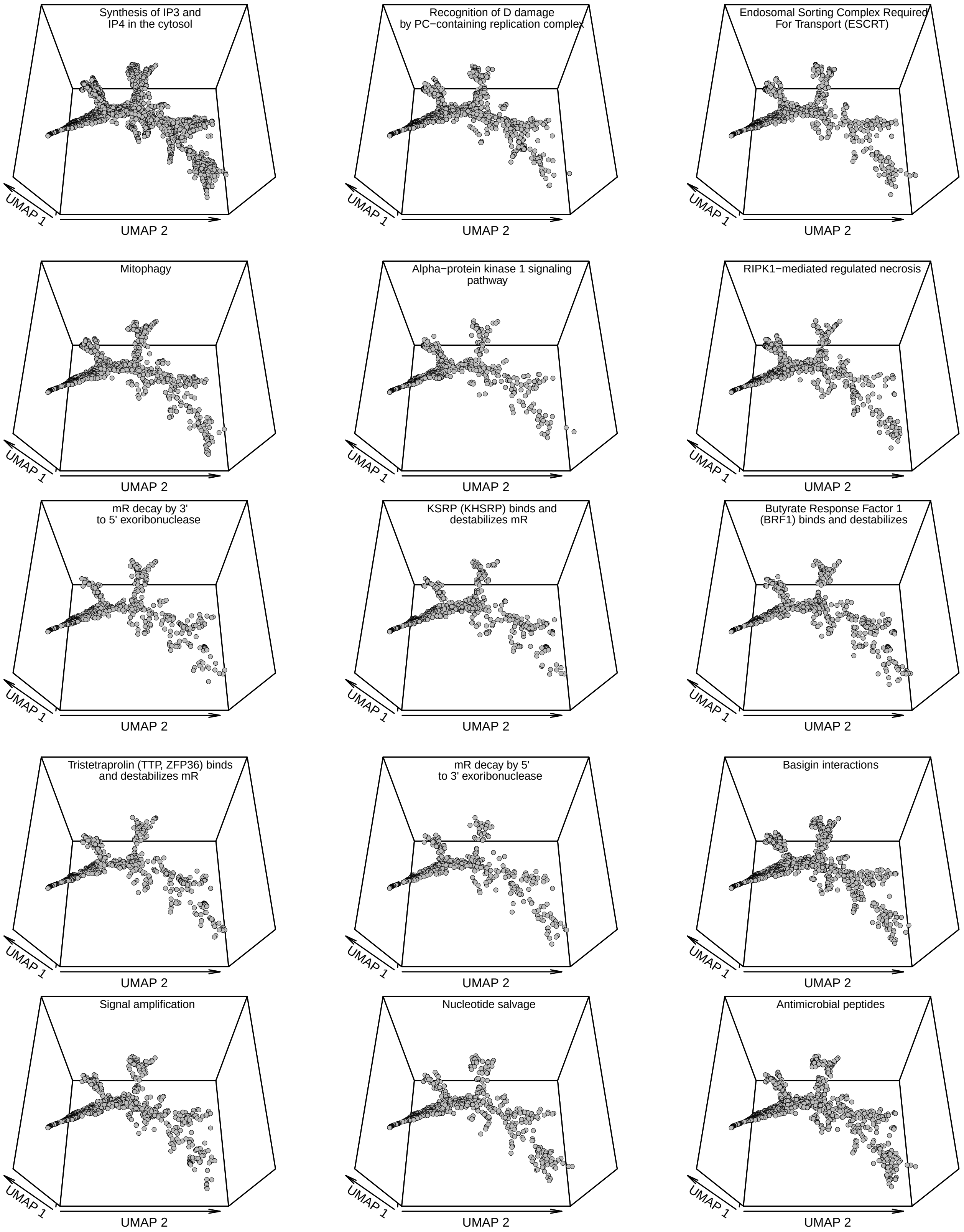
Synthesis of IP3 and
IP4 in the cytosol
UMAP 1
UMAP 2
Recognition of D damage
by PC−containing replication complex
UMAP 1
UMAP 2
Endosomal Sorting Complex Required
For Transport (ESCRT)
UMAP 1
UMAP 2
Mitophagy
UMAP 1
UMAP 2
Alpha−protein kinase 1 signaling
pathway
UMAP 1
UMAP 2
RIPK1−mediated regulated necrosis
UMAP 1
UMAP 2
mR decay by 3'
to 5' exoribonuclease
UMAP 1
UMAP 2
KSRP (KHSRP) binds and
destabilizes mR
UMAP 1
UMAP 2
Butyrate Response Factor 1
(BRF1) binds and destabilizes
UMAP 1
UMAP 2
Tristetraprolin (TTP, ZFP36) binds
and destabilizes mR
UMAP 1
UMAP 2
mR decay by 5'
to 3' exoribonuclease
UMAP 1
UMAP 2
Basigin interactions
UMAP 1
UMAP 2
Signal amplification
UMAP 1
UMAP 2
Nucleotide salvage
UMAP 1
UMAP 2
Antimicrobial peptides
UMAP 1
UMAP 2

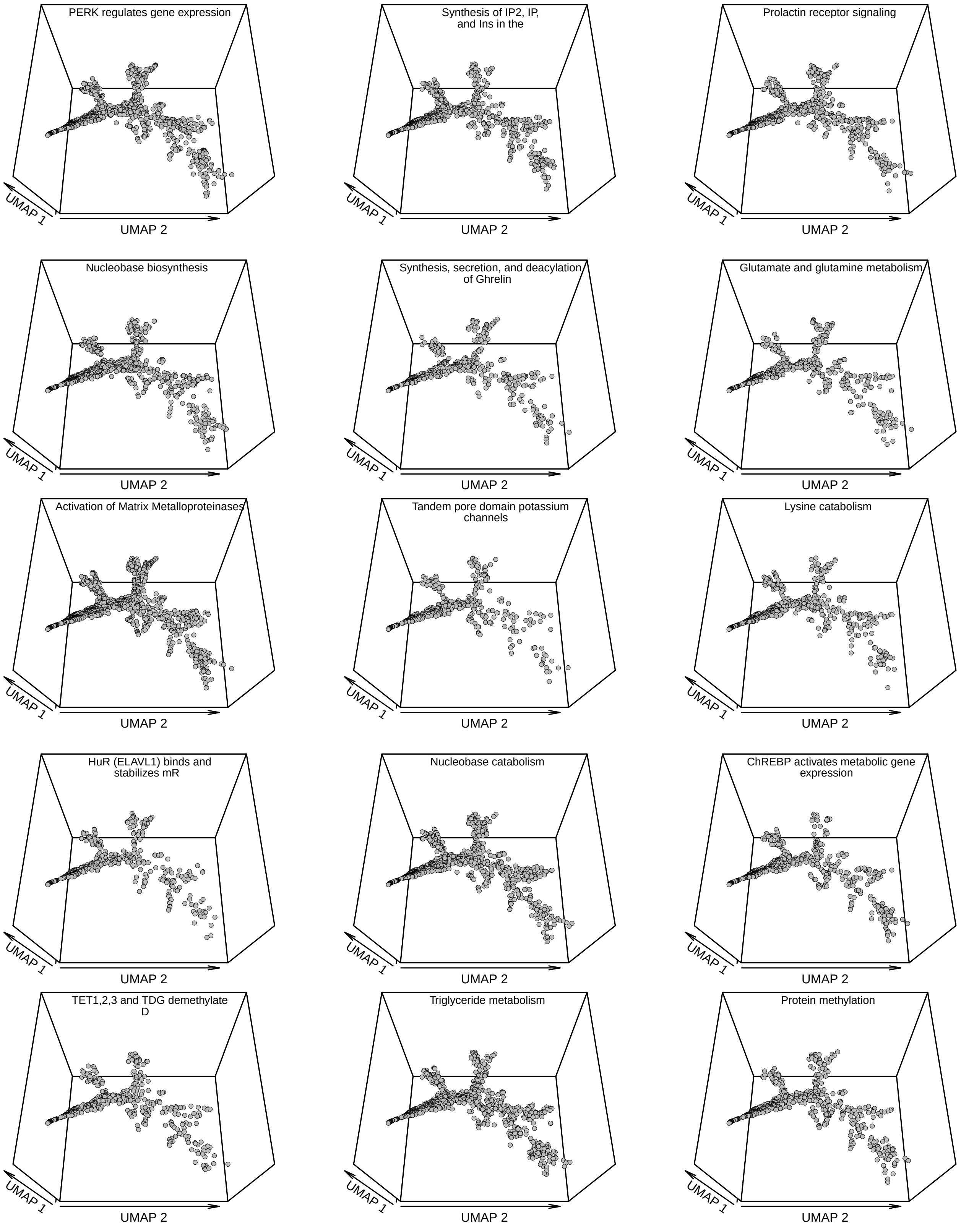
PERK regulates gene expression
UMAP 1
UMAP 2
Synthesis of IP2, IP, and Ins in the
UMAP 1
UMAP 2
Prolactin receptor signaling
UMAP 1
UMAP 2
Nucleobase biosynthesis
UMAP 1
UMAP 2
Synthesis, secretion, and deacylation of Ghrelin
UMAP 1
UMAP 2
Glutamate and glutamine metabolism
UMAP 1
UMAP 2
Activation of Matrix Metalloproteinases
UMAP 1
UMAP 2
Tandem pore domain potassium channels
UMAP 1
UMAP 2
Lysine catabolism
UMAP 1
UMAP 2
HuR (ELAVL1) binds and stabilizes mR
UMAP 1
UMAP 2
Nucleobase catabolism
UMAP 1
UMAP 2
ChREBP activates metabolic gene expression
UMAP 1
UMAP 2
TET1,2,3 and TDG demethylate D
UMAP 1
UMAP 2
Triglyceride metabolism
UMAP 1
UMAP 2
Protein methylation
UMAP 1
UMAP 2

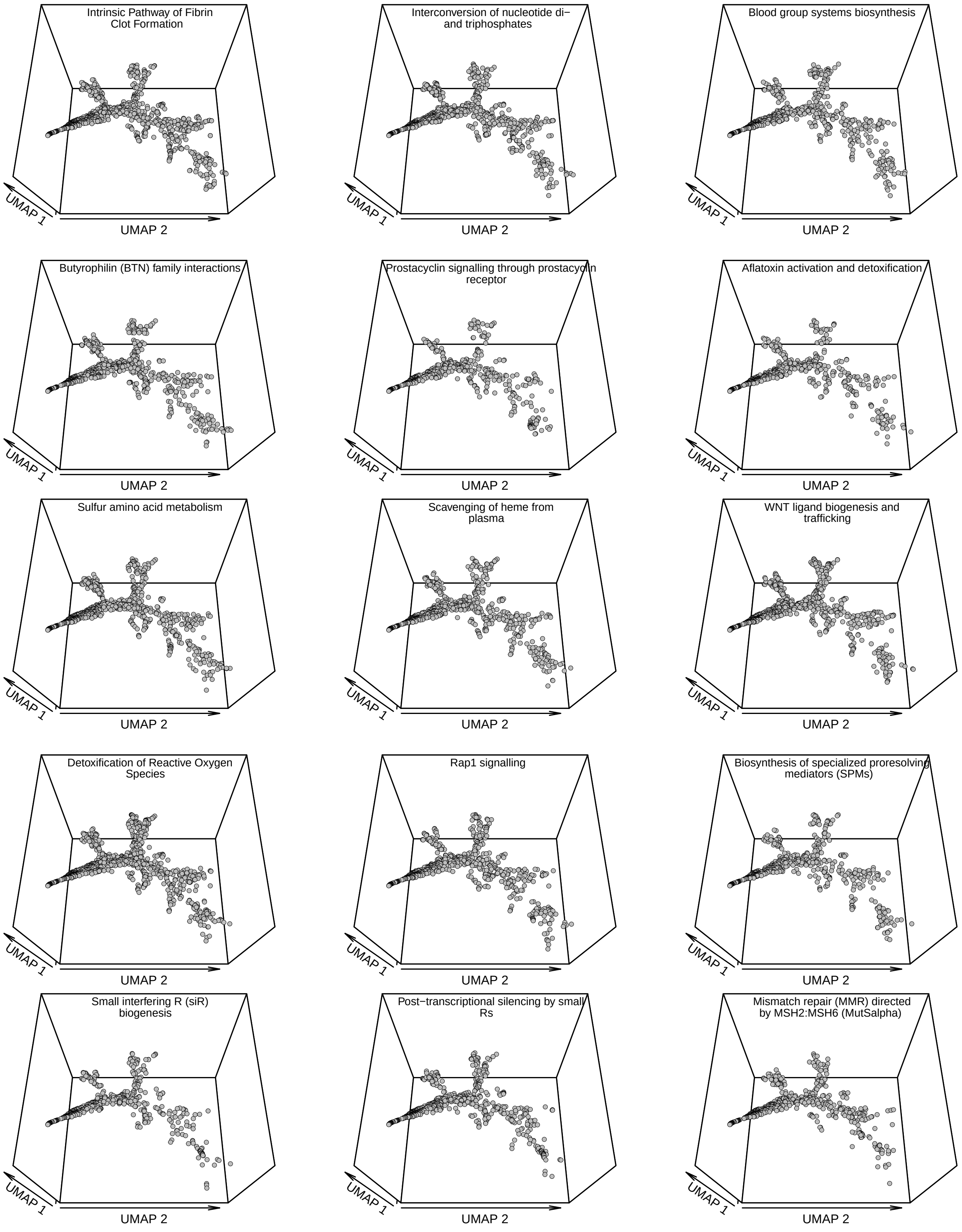
Intrinsic Pathway of Fibrin Clot Formation
UMAP 1
UMAP 2
Interconversion of nucleotide di− and triphosphates
UMAP 1
UMAP 2
Blood group systems biosynthesis
UMAP 1
UMAP 2
Butyrophilin (BTN) family interactions
UMAP 1
UMAP 2
Prostacyclin signalling through prostacyclin receptor
UMAP 1
UMAP 2
Aflatoxin activation and detoxification
UMAP 1
UMAP 2
Sulfur amino acid metabolism
UMAP 1
UMAP 2
Scavenging of heme from plasma
UMAP 1
UMAP 2
WNT ligand biogenesis and trafficking
UMAP 1
UMAP 2
Detoxification of Reactive Oxygen Species
UMAP 1
UMAP 2
Rap1 signalling
UMAP 1
UMAP 2
Biosynthesis of specialized proresolving mediators (SPMs)
UMAP 1
UMAP 2
Small interfering R (siR) biogenesis
UMAP 1
UMAP 2
Post−transcriptional silencing by small Rs
UMAP 1
UMAP 2
Mismatch repair (MMR) directed by MSH2:MSH6 (MutSalpha)
UMAP 1
UMAP 2

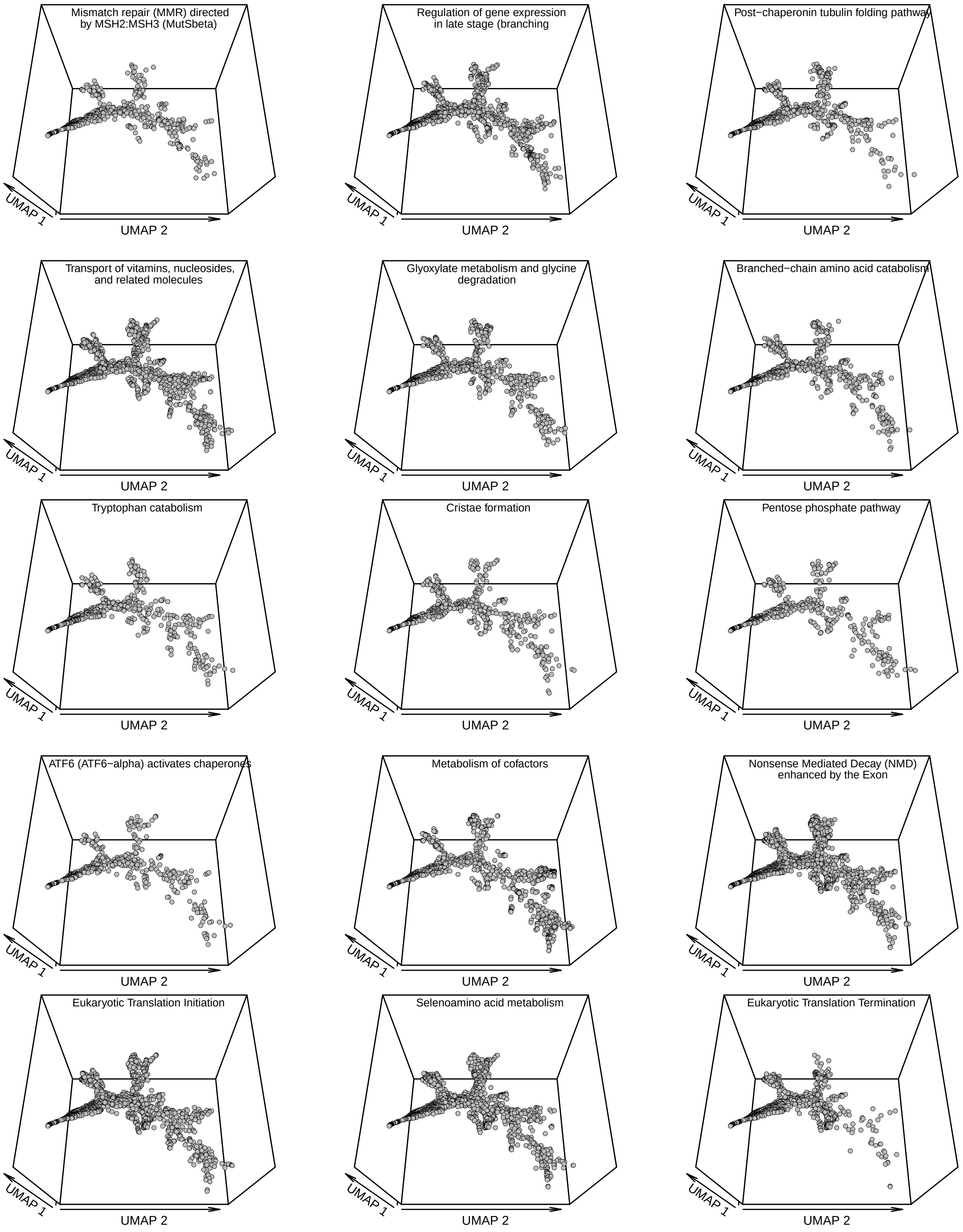
Mismatch repair (MMR) directed by MSH2:MSH3 (MutSbeta)
Regulation of gene expression in late stage (branching
Post−chaperonin tubulin folding pathway
Transport of vitamins, nucleosides, and related molecules
Glyoxylate metabolism and glycine degradation
Branched−chain amino acid catabolism
Tryptophan catabolism
Cristae formation
Pentose phosphate pathway
ATF6 (ATF6−alpha) activates chaperones
Metabolism of cofactors
Nonsense Mediated Decay (NMD) enhanced by the Exon
Eukaryotic Translation Initiation
Selenoamino acid metabolism
Eukaryotic Translation Termination
UMAP 1
UMAP 2

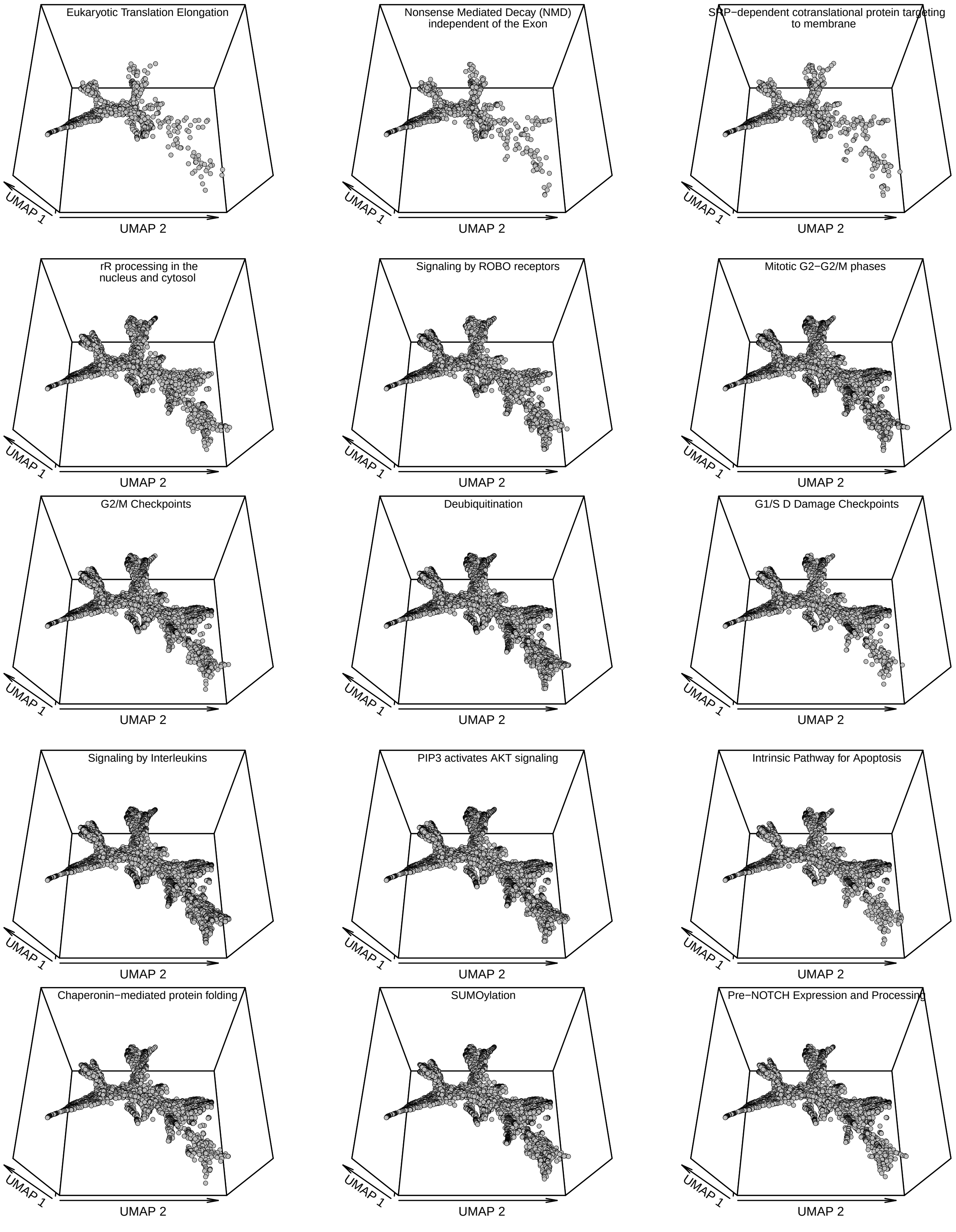
Eukaryotic Translation Elongation
Nonsense Mediated Decay (NMD) independent of the Exon
SRP−dependent cotranslational protein targeting to membrane
rR processing in the nucleus and cytosol
Signaling by ROBO receptors
Mitotic G2−G2/M phases
G2/M Checkpoints
Deubiquitination
G1/S D Damage Checkpoints
Signaling by Interleukins
PIP3 activates AKT signaling
Intrinsic Pathway for Apoptosis
Chaperonin−mediated protein folding
SUMOylation
Pre−NOTCH Expression and Processing
UMAP 1
UMAP 2

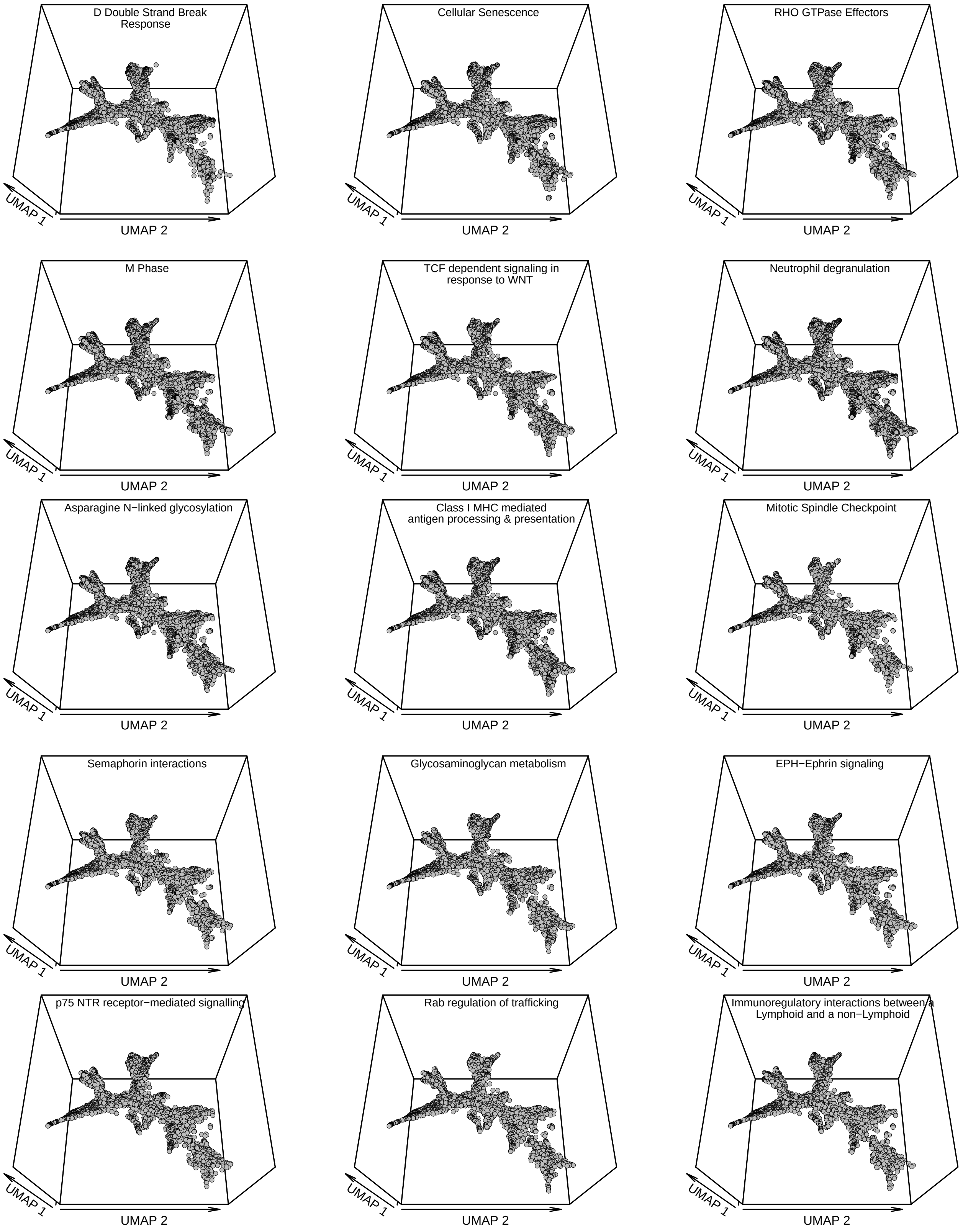
D Double Strand Break Response
UMAP 1
UMAP 2
Cellular Senescence
UMAP 1
UMAP 2
RHO GTPase Effectors
UMAP 1
UMAP 2
M Phase
UMAP 1
UMAP 2
TCF dependent signaling in response to WNT
UMAP 1
UMAP 2
Neutrophil degranulation
UMAP 1
UMAP 2
Asparagine N−linked glycosylation
UMAP 1
UMAP 2
Class I MHC mediated antigen processing & presentation
UMAP 1
UMAP 2
Mitotic Spindle Checkpoint
UMAP 1
UMAP 2
Semaphorin interactions
UMAP 1
UMAP 2
Glycosaminoglycan metabolism
UMAP 1
UMAP 2
EPH−Ephrin signaling
UMAP 1
UMAP 2
p75 NTR receptor−mediated signalling
UMAP 1
UMAP 2
Rab regulation of trafficking
UMAP 1
UMAP 2
Immunoregulatory interactions between a Lymphoid and a non−Lymphoid
UMAP 1
UMAP 2

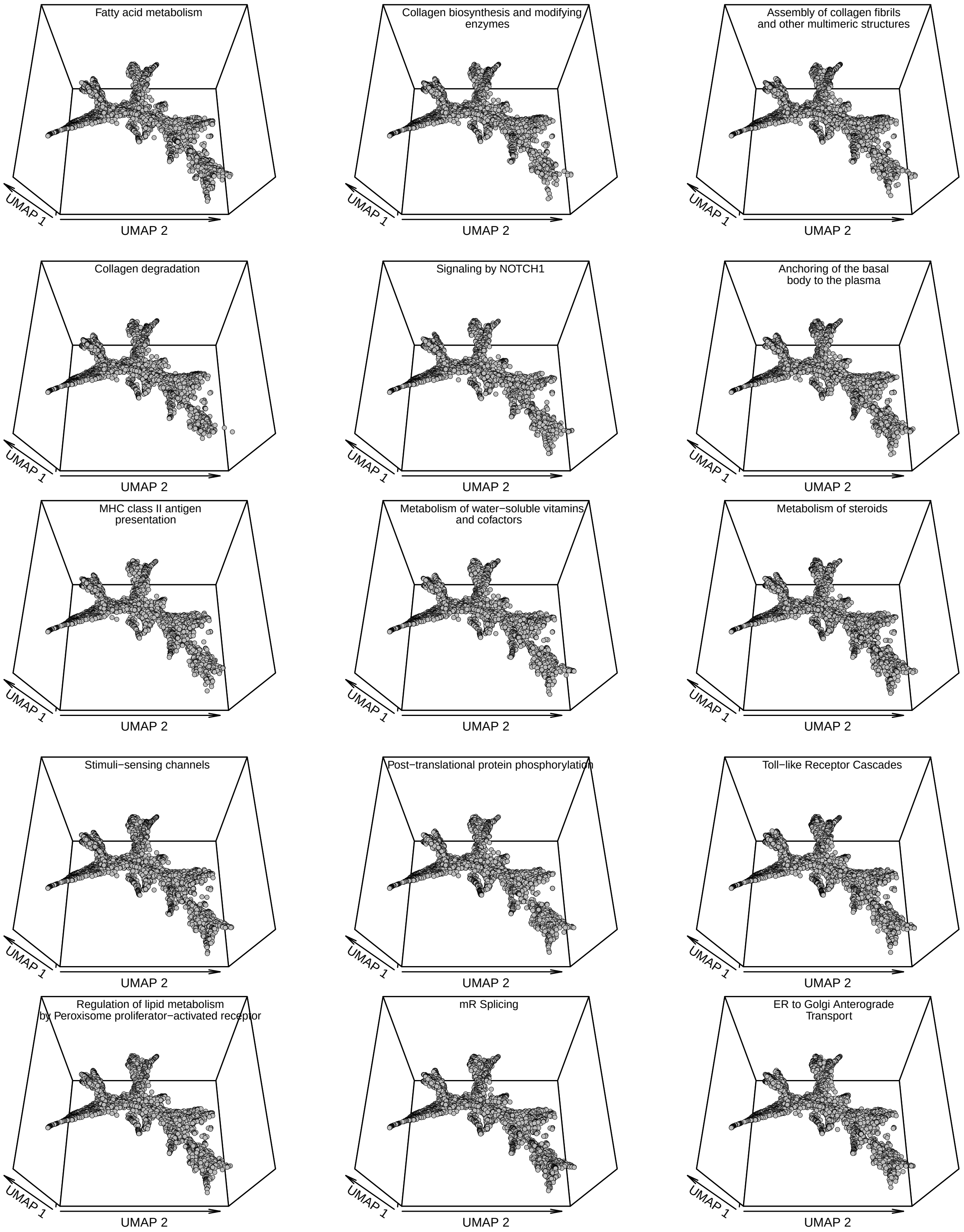
Fatty acid metabolism
UMAP 1
UMAP 2
Collagen biosynthesis and modifying enzymes
UMAP 1
UMAP 2
Assembly of collagen fibrils and other multimeric structures
UMAP 1
UMAP 2
Collagen degradation
UMAP 1
UMAP 2
Signaling by NOTCH1
UMAP 1
UMAP 2
Anchoring of the basal body to the plasma
UMAP 1
UMAP 2
MHC class II antigen presentation
UMAP 1
UMAP 2
Metabolism of water−soluble vitamins and cofactors
UMAP 1
UMAP 2
Metabolism of steroids
UMAP 1
UMAP 2
Stimuli−sensing channels
UMAP 1
UMAP 2
Post−translational protein phosphorylation
UMAP 1
UMAP 2
Toll−like Receptor Cascades
UMAP 1
UMAP 2
Regulation of lipid metabolism by Peroxisome proliferator−activated receptor
UMAP 1
UMAP 2
mR Splicing
UMAP 1
UMAP 2
ER to Golgi Anterograde Transport
UMAP 1
UMAP 2

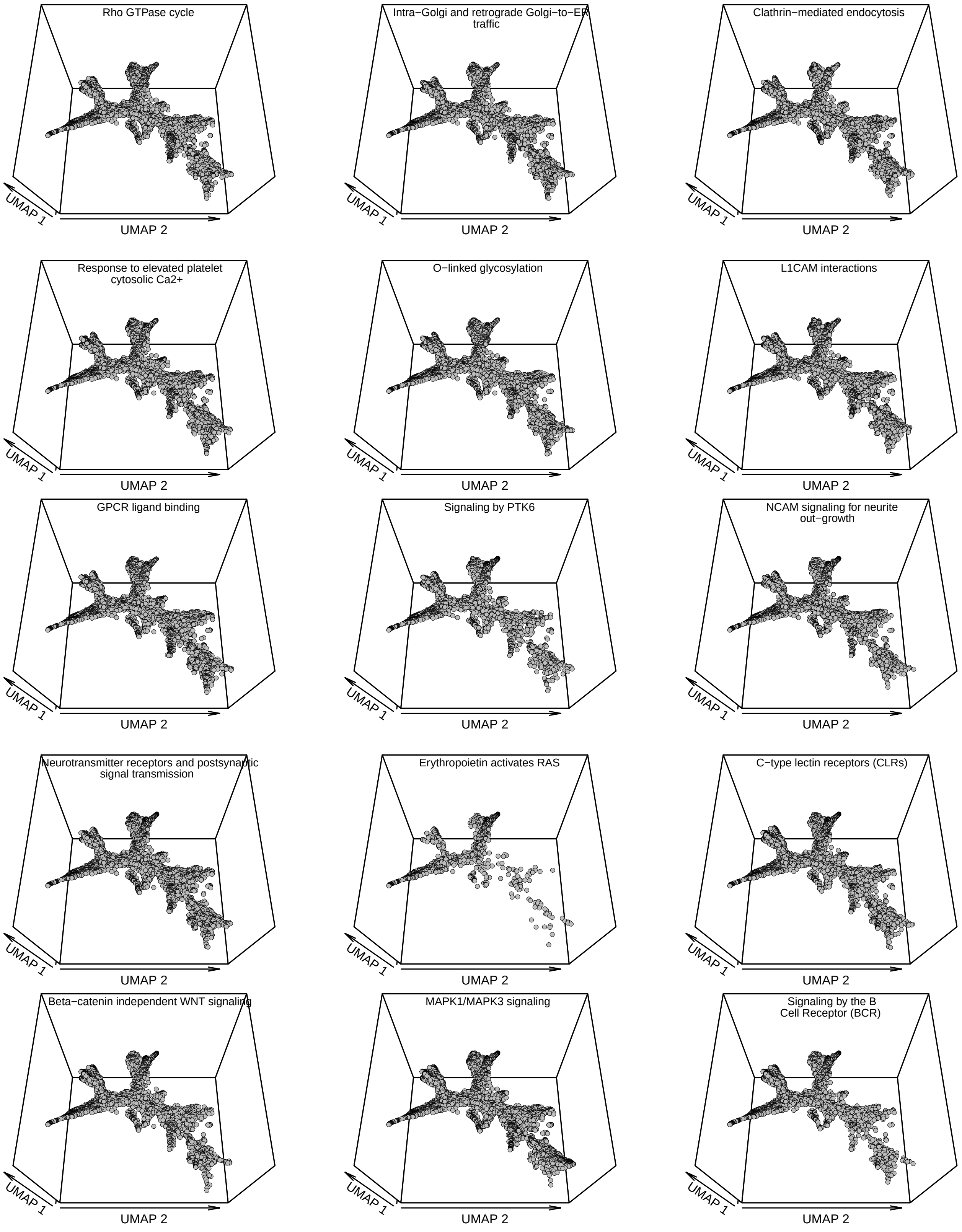
Rho GTPase cycle
Intra−Golgi and retrograde Golgi−to−ER traffic
Clathrin−mediated endocytosis
Response to elevated platelet cytosolic Ca2+
O−linked glycosylation
L1CAM interactions
GPCR ligand binding
Signaling by PTK6
NCAM signaling for neurite out−growth
Neurotransmitter receptors and postsynaptic signal transmission
Erythropoietin activates RAS
C−type lectin receptors (CLRs)
Beta−catenin independent WNT signaling
MAPK1/MAPK3 signaling
Signaling by the B Cell Receptor (BCR)
UMAP 1
UMAP 2

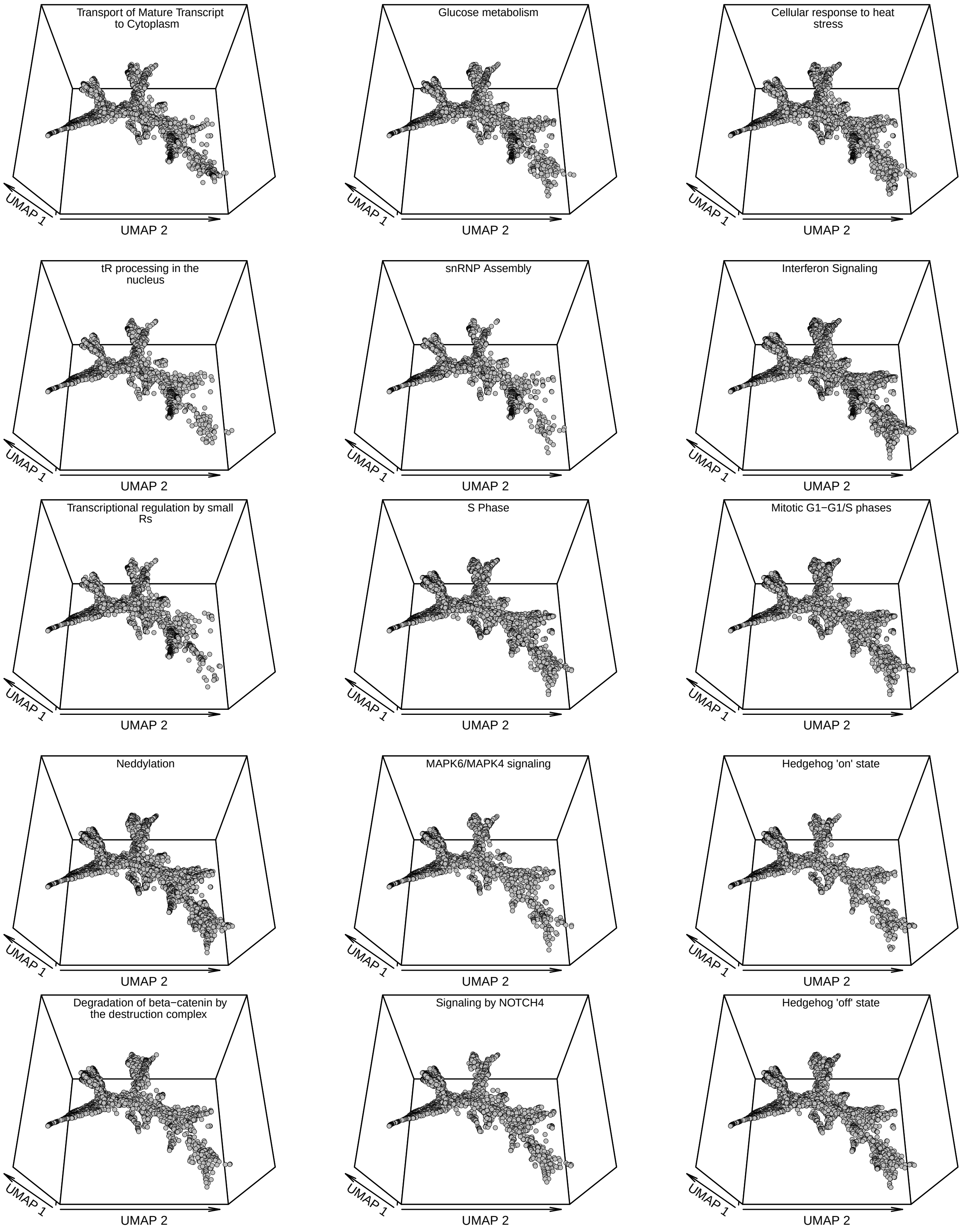
Transport of Mature Transcript to Cytoplasm
UMAP 1
UMAP 2
Glucose metabolism
UMAP 1
UMAP 2
Cellular response to heat stress
UMAP 1
UMAP 2
tR processing in the nucleus
UMAP 1
UMAP 2
snRNP Assembly
UMAP 1
UMAP 2
Interferon Signaling
UMAP 1
UMAP 2
Transcriptional regulation by small Rs
UMAP 1
UMAP 2
S Phase
UMAP 1
UMAP 2
Mitotic G1−G1/S phases
UMAP 1
UMAP 2
Neddylation
UMAP 1
UMAP 2
MAPK6/MAPK4 signaling
UMAP 1
UMAP 2
Hedgehog 'on' state
UMAP 1
UMAP 2
Degradation of beta−catenin by the destruction complex
UMAP 1
UMAP 2
Signaling by NOTCH4
UMAP 1
UMAP 2
Hedgehog 'off' state
UMAP 1
UMAP 2

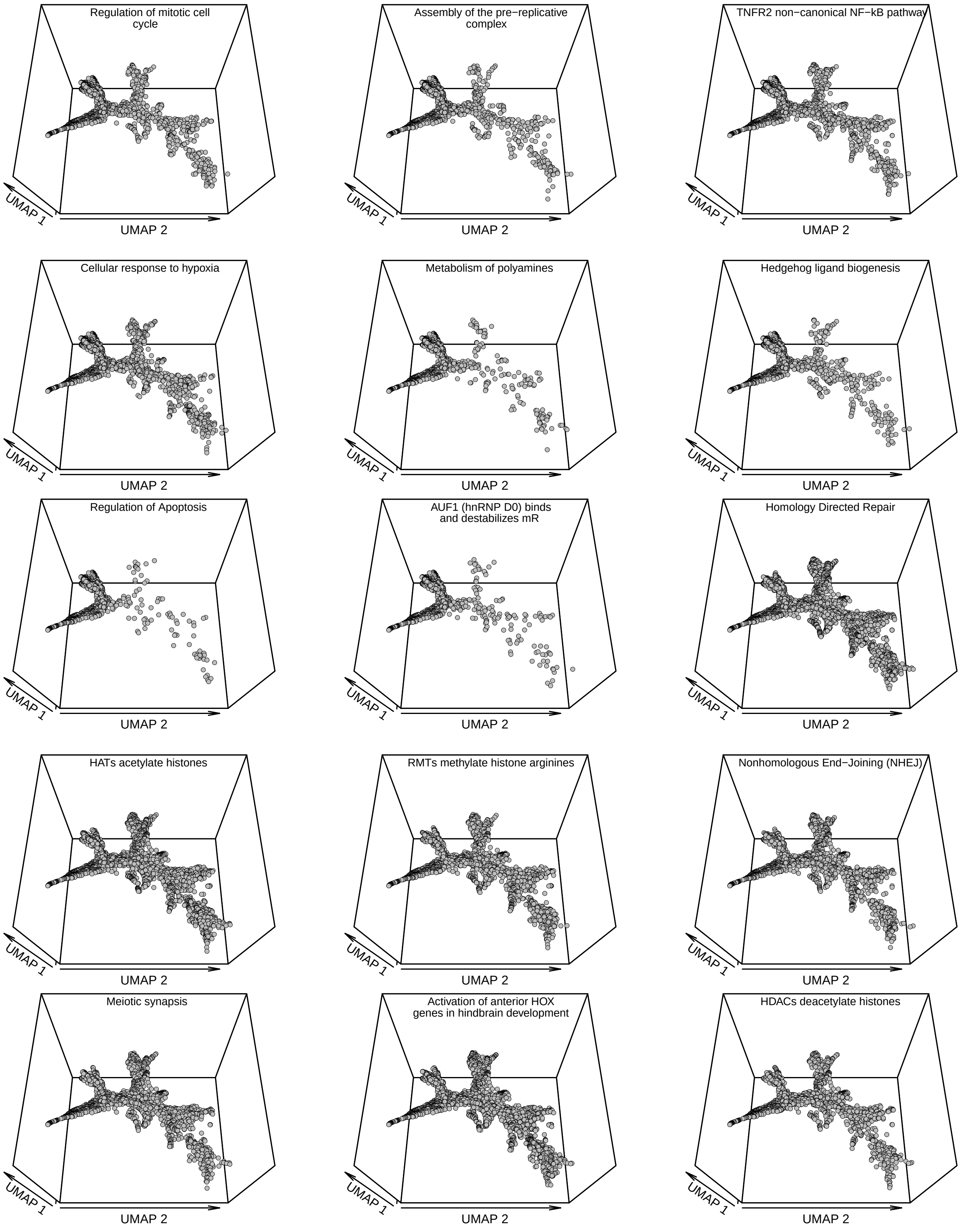
Regulation of mitotic cell cycle
UMAP 1
UMAP 2
Assembly of the pre−replicative complex
UMAP 1
UMAP 2
TNFR2 non−canonical NF−kB pathway
UMAP 1
UMAP 2
Cellular response to hypoxia
UMAP 1
UMAP 2
Metabolism of polyamines
UMAP 1
UMAP 2
Hedgehog ligand biogenesis
UMAP 1
UMAP 2
Regulation of Apoptosis
UMAP 1
UMAP 2
AUF1 (hnRNP D0) binds and destabilizes mR
UMAP 1
UMAP 2
Homology Directed Repair
UMAP 1
UMAP 2
HATs acetylate histones
UMAP 1
UMAP 2
RMTs methylate histone arginines
UMAP 1
UMAP 2
Nonhomologous End−Joining (NHEJ)
UMAP 1
UMAP 2
Meiotic synapsis
UMAP 1
UMAP 2
Activation of anterior HOX genes in hindbrain development
UMAP 1
UMAP 2
HDACs deacetylate histones
UMAP 1
UMAP 2

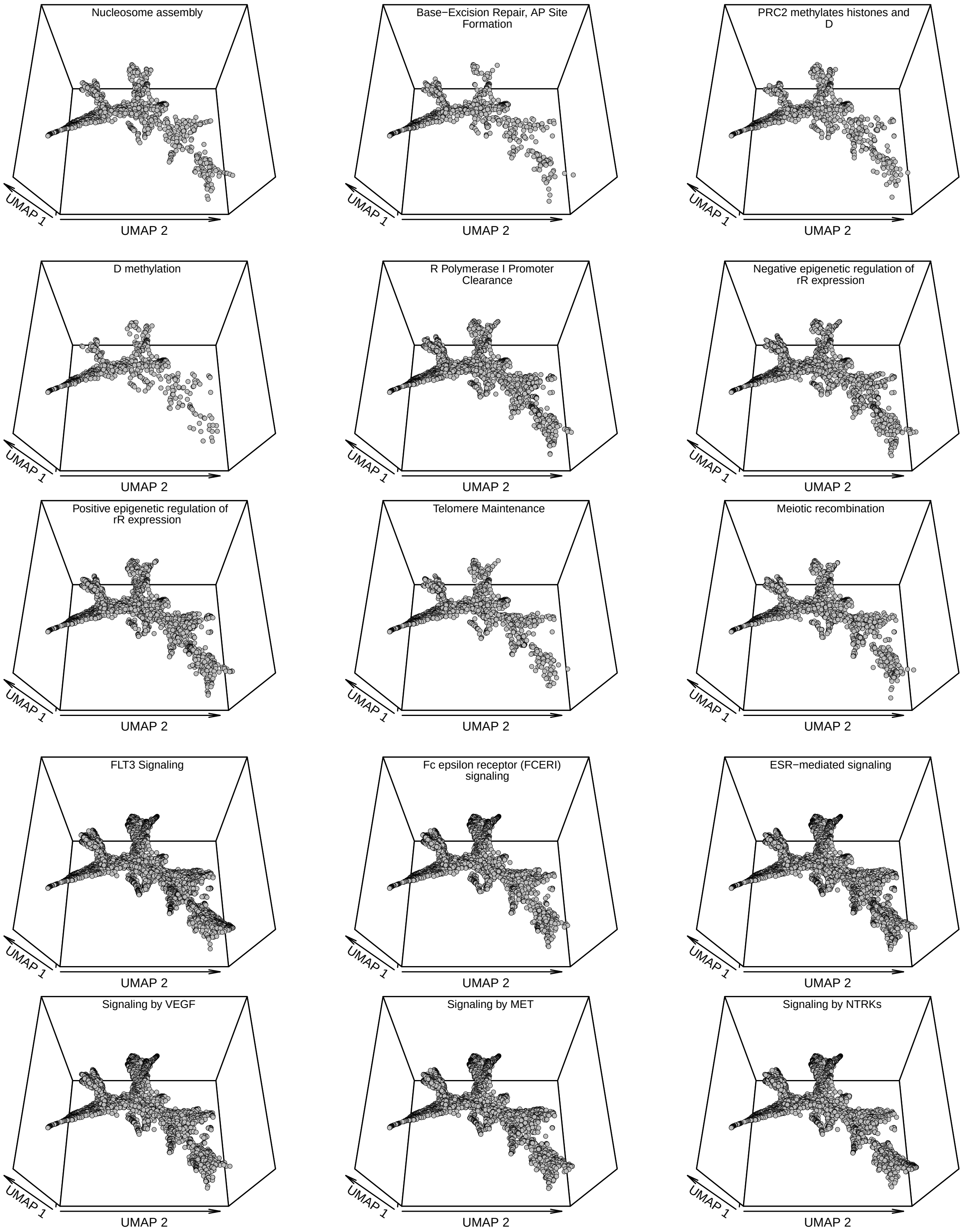
Nucleosome assembly
UMAP 1
UMAP 2
Base−Excision Repair, AP Site Formation
UMAP 1
UMAP 2
PRC2 methylates histones and D
UMAP 1
UMAP 2
D methylation
UMAP 1
UMAP 2
R Polymerase I Promoter Clearance
UMAP 1
UMAP 2
Negative epigenetic regulation of rR expression
UMAP 1
UMAP 2
Positive epigenetic regulation of rR expression
UMAP 1
UMAP 2
Telomere Maintenance
UMAP 1
UMAP 2
Meiotic recombination
UMAP 1
UMAP 2
FLT3 Signaling
UMAP 1
UMAP 2
Fc epsilon receptor (FCERI) signaling
UMAP 1
UMAP 2
ESR−mediated signaling
UMAP 1
UMAP 2
Signaling by VEGF
UMAP 1
UMAP 2
Signaling by MET
UMAP 1
UMAP 2
Signaling by NTRKs
UMAP 1
UMAP 2

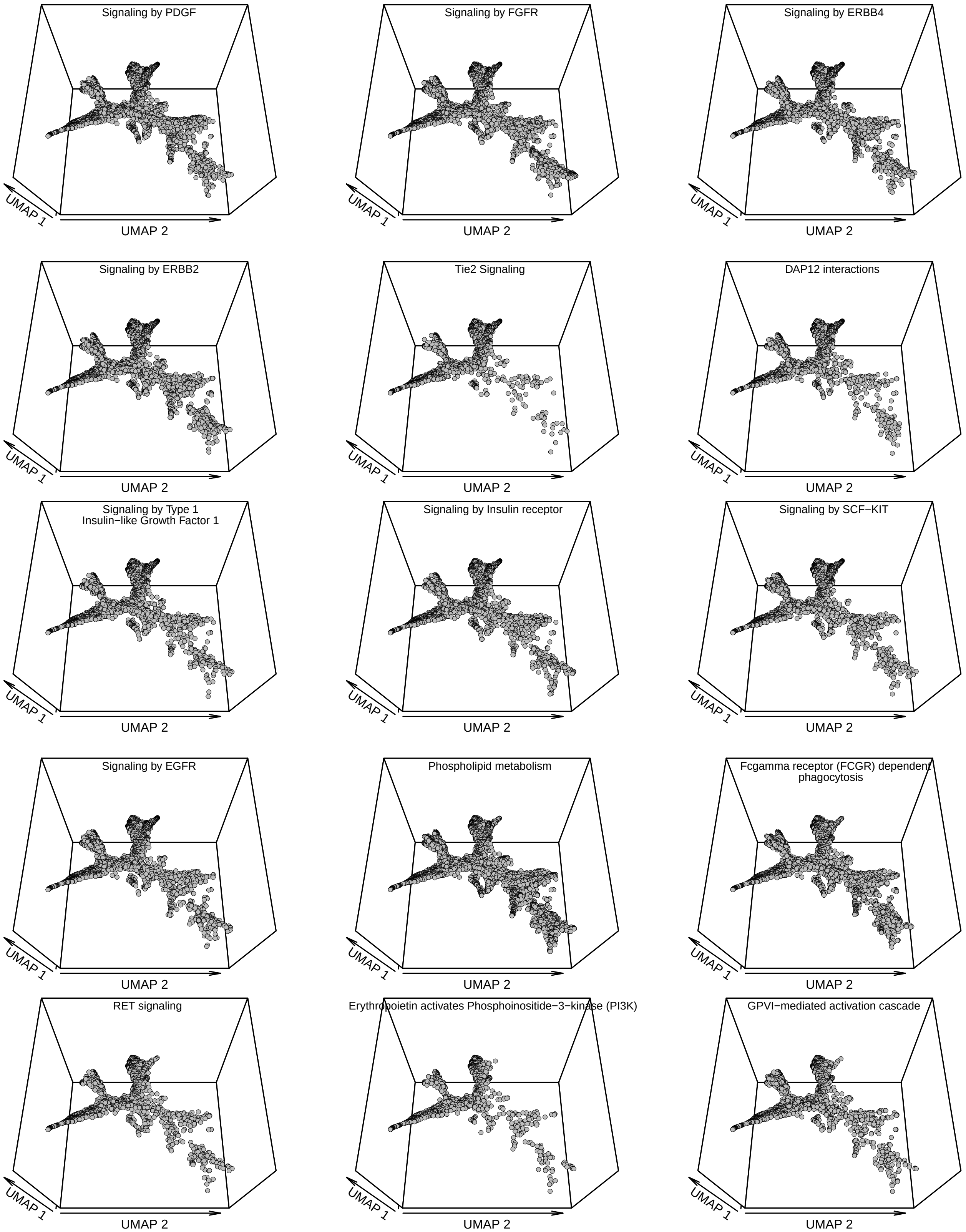

Signaling by PDGF
Signaling by FGFR
Signaling by ERBB4
Signaling by ERBB2
Tie2 Signaling
DAP12 interactions
Signaling by Type 1 Insulin−like Growth Factor 1
Signaling by Insulin receptor
Signaling by SCF−KIT
Signaling by EGFR
Phospholipid metabolism
Fcgamma receptor (FCGR) dependent phagocytosis
RET signaling
Erythropoietin activates Phosphoinositide−3−kinase (PI3K)
GPVI−mediated activation cascade
UMAP 1
UMAP 2

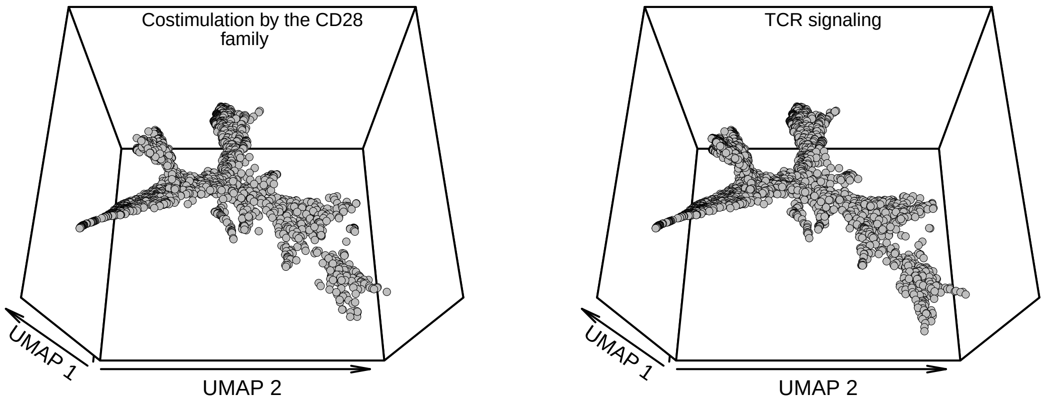


**Supplementary Figure S4. Projection of pathway disruptions onto the UMAP.** Each panel shows one of 377 Reactome pathways projected on the UMAP from **Figure 1D of the main text** ordered by the hierarchical pathway order (column) from **Figure 2 of the main text**.

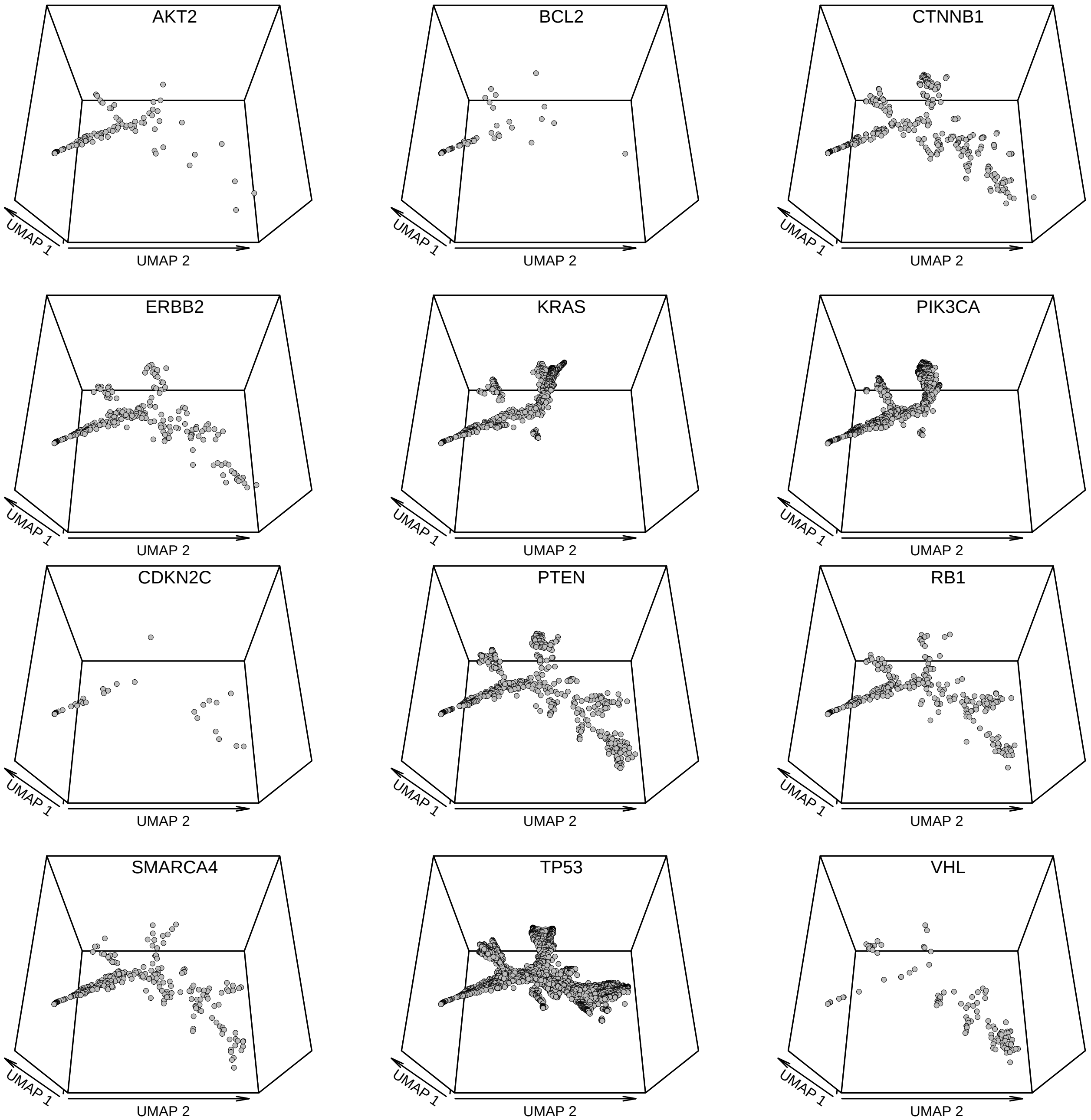


**Supplementary Figure S5. Distribution of oncogenes/tumor suppressors on UMAP clusters.** Each panel shows shows one of the twelve oncogenes/tumor suppressors projected onto the tumors of the UMAP from **Figure 2 of the main text**.
